# Microbiome diversity, intra-mucosal bacteria and immune integration within normal and asthmatic airway mucosa

**DOI:** 10.1101/2024.08.27.609874

**Authors:** Miriam F. Moffatt, Tamiko Nishimura, Michael J. Cox, Claire McBrien, Conor Burke, Leah Cuthbertson, Keir Lewis, Richard Attanoos, Gwyneth Davies, Kian Fan Chung, Jan Lukas Robertus, Jonathan Ish-Horowicz, Orla O’Carroll, John M. Bozeman, Aisling McGowan, Julian M. Hopkin, G. Mark Lathrop, Yasser Riazalhosseini, William O.C. Cookson

## Abstract

Asthma is characterized by reduced bronchial bacterial diversity and airway mucosal disruption. We examined spatial distributions of microbial sequences and host mucosal transcripts in bronchial biopsies from healthy controls and adult asthmatics. Bacteria were discovered by 16S ribosomal RNA staining in the lamina propria of all biopsies, with counts positively associated to lumenal bacterial diversity. Weighted correlation network analysis identified fifteen co-expression networks, including distinct programs of adaptive and innate immunity in differing spatial distributions. Stromal bacterial counts correlated significantly with eight of the network eigenvectors in directions compatible with beneficial relationships. The results suggest that dysbiosis may affect mucosal immunity through impaired interactions beneath the epithelial border. Intra-mucosal companion bacteria may be a potential substrate for selective management of immunity in a wide range of diseases.

**One-Sentence Summary:** The lung microbiome extends within the airway mucosa and associates spatially and functionally with immune networks.

## Main Text

Asthma is an epithelial disorder in the intra-thoracic airways, characterised histologically by mucosal friability, thickening of the basal lamina, and inflammatory cell infiltrates (*1*). Its causes are unknown. Although once uncommon, it now affects 350 million individuals world-wide (*2*). Childhood and adult-onset syndromes are recognised. Eosinophilia and IgE-mediated allergy are more prominent in childhood asthma in westernized societies, but globally most asthma is non-eosinophilic (*2, 3*). Strong host factors have been identified by genome-wide association studies, and the products of asthma susceptibility genes are concentrated in the epithelium(*4*).

Microbiological factors are of central importance to the disease. The thoracic airways carry a conserved bacterial microbiome (*5*) that is thought to regulate mucosal immunity (*6–8*). Early life antibiotics double risk of childhood asthma (*9*). Protective effects of rural living are recognised in children (*10*) and in adults (*11*), and are attributed to benefits from a rich microbial environment. The asthmatic airway microbiome shows typical features of dysbiosis (*12*), with bacterial diversity loss and variable dominance of pathobionts such as *Haemophilus influenzae* (Hi), *Moraxella* spp. (*5, 13–15*), and *Tropheryma whipplei* (*16, 17*). Hi vaccination reduces viral-induced childhood asthmatic wheezing (*18*) and antimicrobial therapy with macrolides is effective in some adults with asthma (*15*).

Acute episodes (attacks) of asthma are precipitated by common respiratory viruses, particularly human rhinovirus (HRV)(*19, 20*), suggesting that the presence of chronic bacterial dysbiosis and dysregulated airway mucosal functions predispose usually benign virus infections to become impactful.

Although extrinsic dysbiosis also accompanies major disorders in the gut, skin and vagina, the mechanisms through which a diverse external microbiome may modulate immunity within the healthy mucosa are not well understood. We therefore spatially mapped interactions between endobronchial bacterial communities and immune cell transcription in the airway epithelium and underlying lamina propria.

Our study design is summarised in Fig. 1. We bronchoscoped 67 asthmatics and 44 healthy controls from three European centres (Table S1). We used 16S rRNA gene amplicon sequences to identify the bacterial microbiota in bronchial brushings to sample the mucus layer and underlying epithelial cells, classifying communities for dysbiotic features such as loss of diversity and evenness, and the abundance of potential pathogens. An experienced lung pathologist scored standard histologic features of bronchial biopsies blind to asthma status. We scored bacterial abundance within the mucosa of 44 biopsies after staining with a 16S rRNA *in situ* hybridization probe and measured human gene expression in epithelial and sub-epithelial compartments using NanoString’s GeoMx Digital Spatial Profiler platform. We used network analysis to identify co-regulated modules of transcripts (*21*) and their spatial distribution within the biopsies.

**Fig. 1.**
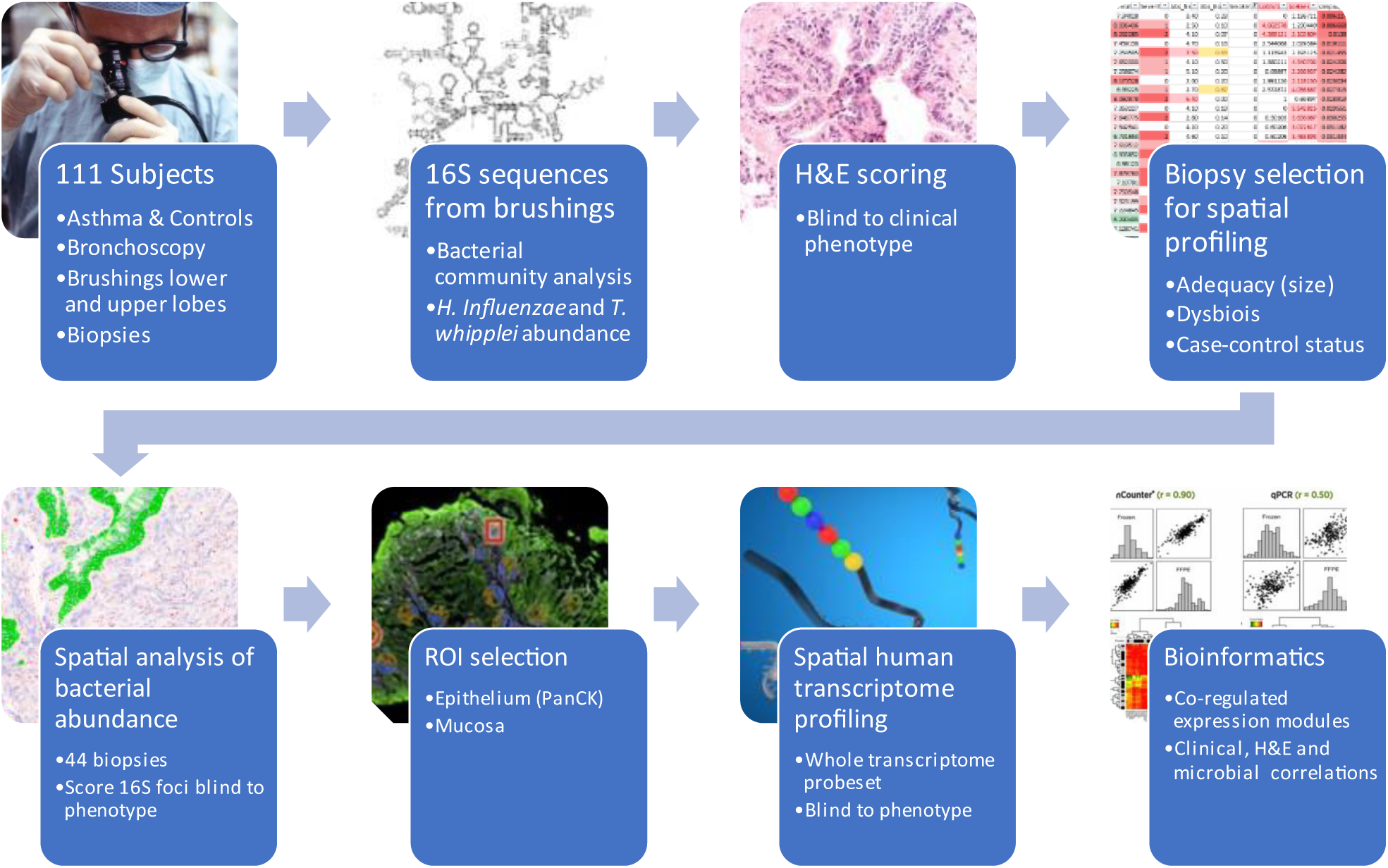
Study Design. Bronchial brushings and biopsies were taken during fibre-optic bronchoscopy of 67 asthmatics and 44 healthy controls. 16SrRNA gene sequencing quantified bacterial communities in bronchial brushings. All biopsies were scored for histological abnormalities, and an optimal subset of 44 biopsies taken through for GeoMx DSP Whole Transcriptome analysis. Bacterial content within biopsies was quantified by 16S *in situ* hybridisation staining. Regions of interest (ROI) were selected and segmented into epithelium or stromal compartments, and their transcriptomes quantified by NGS readout. After QC, network and spatial analysis of transcriptomes were related to clinical, microbial and image results.

We examined correlations between asthma-associated clinical, microbial and biopsy traits in the 111 subjects (Table S2). Significant associations to asthma included peripheral blood eosinophil counts (widely used as a clinical biomarker) (r=0.31, *P*<0.001), Shannon diversity (r=-0.20, *P*=0.009), and basement membrane thickening (r=0.32, *P*<0.001). Abundance of the pathogen *Tropheryma Whipplei* was also associated with asthma (r=0.17, P=0.03) whereas abundance of *Selenomonas* and *Actinomyces* spp., potential indicator species for microbial dysbiosis (*22*), were negatively associated (r=-0.20, *P*=0.01 and r=-0.16, *P*=0.04 respectively).

Inclusion of all associated variables in a forward stepwise logistic regression found independent predictors of asthma status to be basement membrane thickening (*P*=0.00008, Nagelkerke R^2^ 15.9%), Shannon diversity (*P*=0.00008, additional R^2^ 21.2%) and peripheral blood eosinophil counts (*P*=0.01, additional R^2^ 6.0%) (Table S3). These variables were therefore the focus for downstream analyses.

### Bacterial counts within mucosa

Biopsies slides from 44 subjects (chosen for specimen size and case-control status) were scanned microscopically on the GeoMx™ DSP instrument (at 20X). Pan-eubacterial rRNA was detected using the RNAscope® 16S rRNA probe, prior to immunofluorescence with PanCK, and the DNA dye SYTO13.

The results revealed 16S staining consistent with microbial infiltration in the biopsies beneath the airway epithelium (Fig. 2A-C, Fig. S1). The distribution of 16S foci was not uniform across the plane of section, suggesting they were not a consequence of secondary bacterial contamination. Comparison of the 16S rRNA probe staining with a scrambled version of the same probe on a serial section informed as a negative control (Fig. S3).

**Fig. 2.**
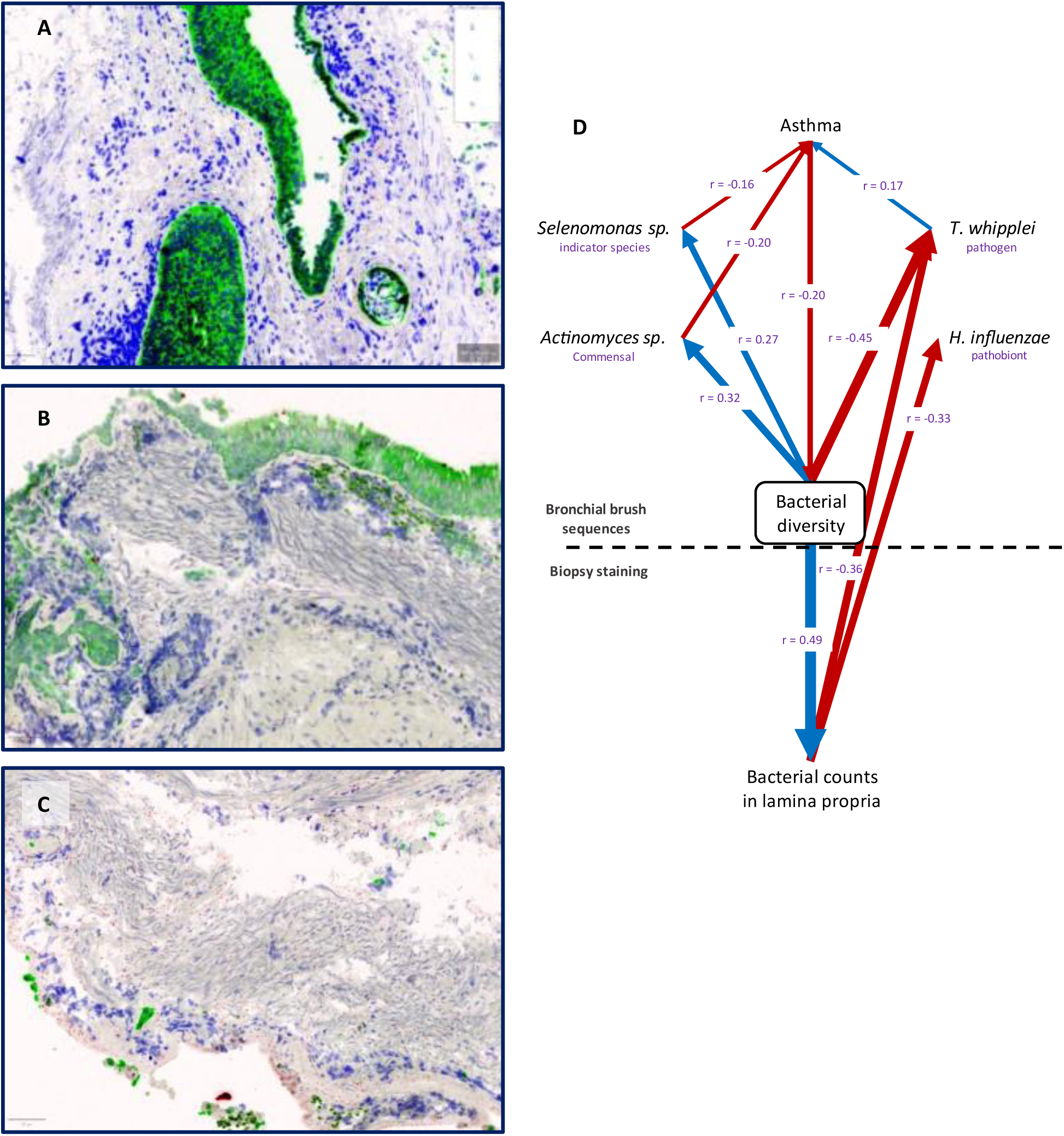
Microbial presence in airway biopsies. (**A**-**C**) 16S staining in airway epithelial biopsies. Green = PanCK (epithelial), blue = DNA (nuclear) and red= 16S rRNA gene probe (bacteria). Relative bacterial counts on a scale of 0-3 in epithelium, basement membrane, and stromal (lamina propria) compartments were derived independently from two observers without knowledge of disease status. Biopsy **A** is from a 41 years-old (y-o) non-asthmatic female (mean 16S abundance score: epithelium=0, basement membrane=1, stroma=3), **B** is from a 24 y-o asthmatic male (16S scores = 2,2,1), and **C** is from a 25 y-o asthmatic male (16S scores = 1.5,3,2). The full set of biopsies is found in Fig. S1. Basement membrane abnormalities are seen in **B** and **C**, and there is extensive epithelial denudation in **C**. (**D**) Pearson correlations between bacterial diversity and species abundance in the airway lumen (accessed through endobronchial brushings) and relative bacterial counts in the lamina propria of all airway biopsies studied by 16S rRNA staining. Blue indicates a positive correlation, red a negative correlation.

The abundance of microbial genomes in epithelium, basement membrane (BM) and lamina propria was graded on a scale of 0-3 independently by two observers, blind to clinical or histological status (Fig. S1 for all images scored). The degree of infiltration varied between biopsies but was present in all samples. The epithelial layer contained a lower abundance of 16S hits than the stroma (score 1.3±0.2 vs 2.1±0.2). Agreement between observers was significant for epithelial scores (Cohen’s Kappa=0.34, *P*=0.003), but higher for basement membrane and stromal scores (Cohen’s Kappa=0.89, *P*=0.000 and 0.74, *P*=0.000 respectively).

Stromal 16S counts were positively associated with lumenal bacterial diversity (r=0.49, *P*=0.000) and negatively associated with the presence of lumenal *H. influenzae* (r=-0.32, *P*=0.007) or *T. whipplei* (r=-0.36, *P*=0.003), suggesting that bacteria within the lamina propria were benign (Fig, 2D, Table S2). Subsequently we refer to them as intra-mucosal companion bacteria (IMCBs).

### Spatial profiling

GeoMx™ digital spatial profiling was performed with the human Whole Transcriptome Atlas (WTA) probe set with read-out by next-generation sequencing (Nanostring Technologies). Regions of interest (ROIs) were identified from each biopsy and segmented into epithelial (PanCK stained) and stromal sub-compartments. 2.64B unique sequence reads were generated from 205 ROIs. After quality control on the GeoMx platform, 12,487 transcripts were included for downstream analysis. The abundance of specific immune cells in each ROI was estimated using the SpatialDecon (*23*) module in the GeoMx analysis suite.

### Network analysis

In general, gene transcription levels are highly co-ordinated functionally within tissues as well as within individual cell types. Co-expressed modules of genes can be discovered by network analyses to provide systematic insights into the integrated activities of tissue or cellular transcriptomes (*21*). Modules can be related through their eigengene vectors to disease and related phenotypes (*24*), including, as we show here, spatial information from the GeoMx™ platform.

We therefore used weighted gene correlation network analysis (WGCNA)(*24*), identifying fifteen modules of highly co-expressed genes (Table S4). Networks were by convention named with colours. The grey module contains uncorrelated transcripts. Functions of hub genes were annotated from the NCBI gene website (https://www.ncbi.nlm.nih.gov/gene/) and limited literature searches (Table S4). Network module eigengenes were related to clinical traits, bacterial scores, and probe-derived putative immune cell counts using WGCNA (Fig. 3 and Table S2). Network eigengenes were not all independent and showed correlations within branches of their phylogenetic tree (Fig. S3).

**Fig. 3.**
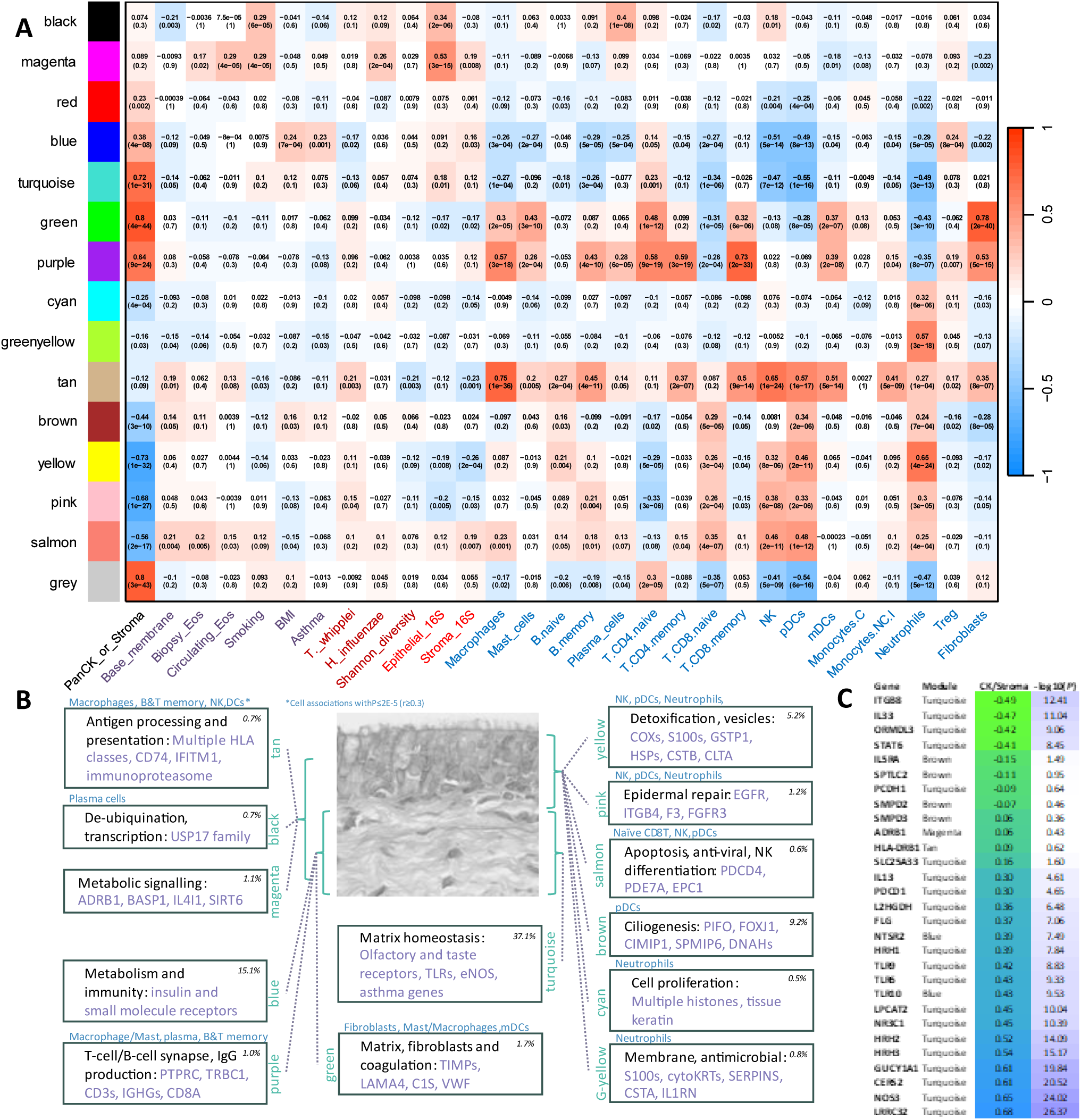
Spatial organization of mucosal transcript networks. (**A**) Phenotypic correlations with WGCNA network module eigengenes. The PanCK or Stroma variable (CKoS) provides localization within the biopsy (negative = epithelial, positive = stromal). Clinical and biopsy parameters are to the left and immune cell transcript abundances to the right. (**B**) Summary of module gene content and localization within the mucosa. (**C**) Localization of transcripts with known functions in asthma pathogenesis, and their module membership. Transcript localization may differ from that of the module eigengene, which is derived from all module members. Other asthma genes and therapeutic targets were either not detectable in the data (*IL1RL1*, *IL4*, *TLR1*, *TLR3*, *TLR4, TLR8*, *IL5*, *IL18R1*, *RORA*) or were unconnected and within the grey module (*TSLP*, *RELA*, *SMAD3* and *GSDMB*).

Spatial context to networks was given by the PanCK_or_Stroma (CKoS) variable (1= when measured in PanCK ROI, 2=stromal ROI), identifying diffuse (non-significant correlation with CKoS), epithelial (negative correlation) and lamina propria (positive correlation) localisations for the network eigenvectors as well as individual transcripts.

### Diffuse modules

The tan, black, and magenta modules were without significant CKoS associations (Fig. 3A) and putative epithelial or stromal localisation.

Tan module hubs included multiple MHC genes, including *HLA-A*, *HLA-B*, *HLA-C*, *HLA-DRB1*, *HLA-E*, *HLA-F* and *HLA-DPA1*, the invariant chain *CD74*, the immunoproteasome subunits *PSMB8*, *PSMB9* and *PSMB10*, the antigen transporter *TAP2* and its binding protein *TAPBP* (Fig. 3B, Table S1). *STAT1*, *IFITM1*, *IFITM3*, and *IRF1* encode major regulators of interferon and anti-viral responses. The tan module eigenvector correlated with multiple immune cell markers (Fig. 3A), including macrophages, B-memory, T-CD8 memory, NK, pDCs, and mDCs (all r ≥ 0.5, *P*≤10^-13^). These findings illustrate that co-expression of multiple HLA classes characterises immune surveillance through the airway mucosa.

Black module hub genes encode deubiquitination and transcription and were associated with plasma cell counts (r=0.4, *P*=10^-8^) (Table S1). Magenta hub genes encode metabolite signalling (glucose *BASP1*, *PDZD2*; aldehyde *ALDH3B2*; epinephrine *ADRB1*, and L-phenylalanine *IL4I1*); senescence (*ATRX*, *SIRT6*); and aspects of inflammation (*PTPB1*, *ST3GAL2*, *ADAM15*, *MAPKBP1*, *SPATA2L* and *EBI3*) (Table S1). The magenta module was not associated with cell-specific markers (Fig. 3A).

### Epithelial modules

Yellow, pink, brown, cyan and greenyellow modules were strongly epithelial and their gene membership had plausible roles in epithelial activities (Fig. 3 and Table S1).

Yellow hub genes encode cytochrome oxidases (*COX6A1*, *COX7C* and *COX5A*), detoxification (*GSTP1*) and vesicle formation (*CDC42*, *CLTA*, *VAMP8* and *HSPA8*) (Table S1). The module eigenvector correlated strongly with neutrophil markers (r=0.65, *P*<10^-23^) (Fig. 3A).

Brown hub genes enact ciliogenesis (*PIFO*, *FOXJ1*, *CIMIP1*, *SPMIP6*, *DNAH*s) (Table S1), and pink hub genes may mediate epithelial border repair (*EFGR*, *ITGB4*, *F3*, *FGFR3*) (Table S1).

The greenyellow hub was *KRT13* (MM=0.95, *P*<10^-97^), which is the primary marker for recently described airway hillock injury-resistant stem cells (*25*). Hillock stem cell differentiation is essential for resurfacing denuded airway epithelium (*25*). We show here *KRT13* is co-expressed with genes encoding multiple cytokeratins (*KRT5*, *KRT6A*, *KRT13*, *KRT16*, *KRT23*, and *KRT24*); antimicrobial and keratin-associated *SPRR3* and *SPRR1B*; multiple serpins (*SERPINB1*,

*SERPINB2*, *SERPINB13* and *SERPINB5*), cystatin A (*CSTA*), and the interleukin 1 inhibitor *IL1RN* (Table S1).

The salmon module contained genes involved with apoptosis (*PDCD4*, *EPC1*, *IL24*, *BCL2L2*, *ERF*) and NK cell differentiation (*PDE7A*, *EPC1*, *FIZ1*, *TUSC2*, and *ZBTB7B*) (Table S1), and was associated with NK (r=0.46, *P*<10^-10^) and pDC (r=0.48, *P*=10^-12^) cell markers (Fig. 3A).

### Stromal modules

Within the stroma, the purple module contained multiple components of the canonical antigen-specific response, including genes encoding T-cell receptor components (*TRBC1*, *CD3E*, *CD3D*, *TRAC*, *CD8A*); the T-cell/B-cell synapse (B-cell antigen receptor *CD79A*: Fc receptor for IgM (*FCMR*)), integrin complex genes (*ITGA4*, *ITGB2*, *ITGAL*, *ITGAX* and *CD37*); and Immunoglobulin G heavy chains (IGHG3 and IGHG2) (Table S1). Immune cell associations included macrophages (r=0.57, *P*<10^-17^), plasma cells (r=0.43, *P*<10^-9^), T-CD4 naïve (r=0.58, *P*<10^-18^) T-CD4 memory (r=0.59, *P*<10^-18^) and T-CD8 memory cells (r=0.73, *P*<10^-32^) (Fig. 3A).

Green module genes encode multiple functions in matrix maintenance, including collagens (*COL6A2*, *COL6A1*, *COL4A1*, *COL4A2*) laminins and filaments (*LAMA4*, *VIM*, *DCN*); matrix binding (*LGALS1*, *SPARCL1*); protease inhibition (*TIMP2*, *TIMP3*, *A2M*) and coagulation (*SERPING1*, *C1S*, *VWF*) (Table S1). The IgA heavy chain (*IGHA1*) and its accessory *JCHAIN* were in this module. Strong associations were seen to fibroblasts (r=0.78, *P*<10^-39^), naïve T-CD8 cells (r=0.48, *P=*10^-12^), and mast cells (r=0.43, *P*<10^-9^) (Fig. 3A).

Genes within the blue module modulate metabolism including *POC1A* and *NEUROD1* (insulin); *NTSR2* (histamine); *PTGER2* (prostaglandin); *PTHLH* (calcium); *HTR1E* (serotonin); *CA6* (CO_2_); and *GRM8* (glutamate) (Table S1). Other genes encode aspects of innate immunity (*LRRC24*, *MARCO*, *MAPK9*, *CD72*, *PDCD1LG2*, *TPRG1*) (Table S1). The module eigenvector was associated with Treg markers (r=0.24, *P*<10^-3^) (Fig. 3A).

Red module hub genes had obscure functions, perhaps involving tyrosine kinase activities, and the module eigengene did not associate with immune cell types or clinical parameters (Table S1).

The turquoise module contained 37.1% of all transcripts, and its encoded biological functions appeared complex and multi-themed. Module members include striking numbers of olfactory receptors (10% of genes with MM >0.7), taste receptors (7 in module), *TLR9* and *TLR6*, and the cannabinoid receptor *CNR1* (Table S1). The module also contained T-cell associated-genes (*PTPN22*, *CCR7*, *FCRL4*, *KLRB1*, *KLRC4*, *GIMAP8*, *PIK3R5*, *CD160*) (Table S1), consistent with the module eigenvector association with naïve T-CD4 cell markers (r=0.23, *P*=10^-3^) (Fig. 3A).

### Trait and module associations

Possible functions of IMCBs may be inferred from associations between stromal 16S counts and network module eigengenes. Homeostatic roles for IMCBs are supported by negative associations with the tan (antigen presentation) module (Fig. 4A), the green (matrix and fibroblast) module, and yellow, pink, and cyan modules (encoding genes for detoxification, epidermal repair, and neutrophil markers). Stromal bacterial counts were positively associated with metabolically related magenta and blue modules. Shannon diversity of the lumenal bacteria was also negatively associated with the antigen presentation module (Fig. 4B).

**Fig. 4.**
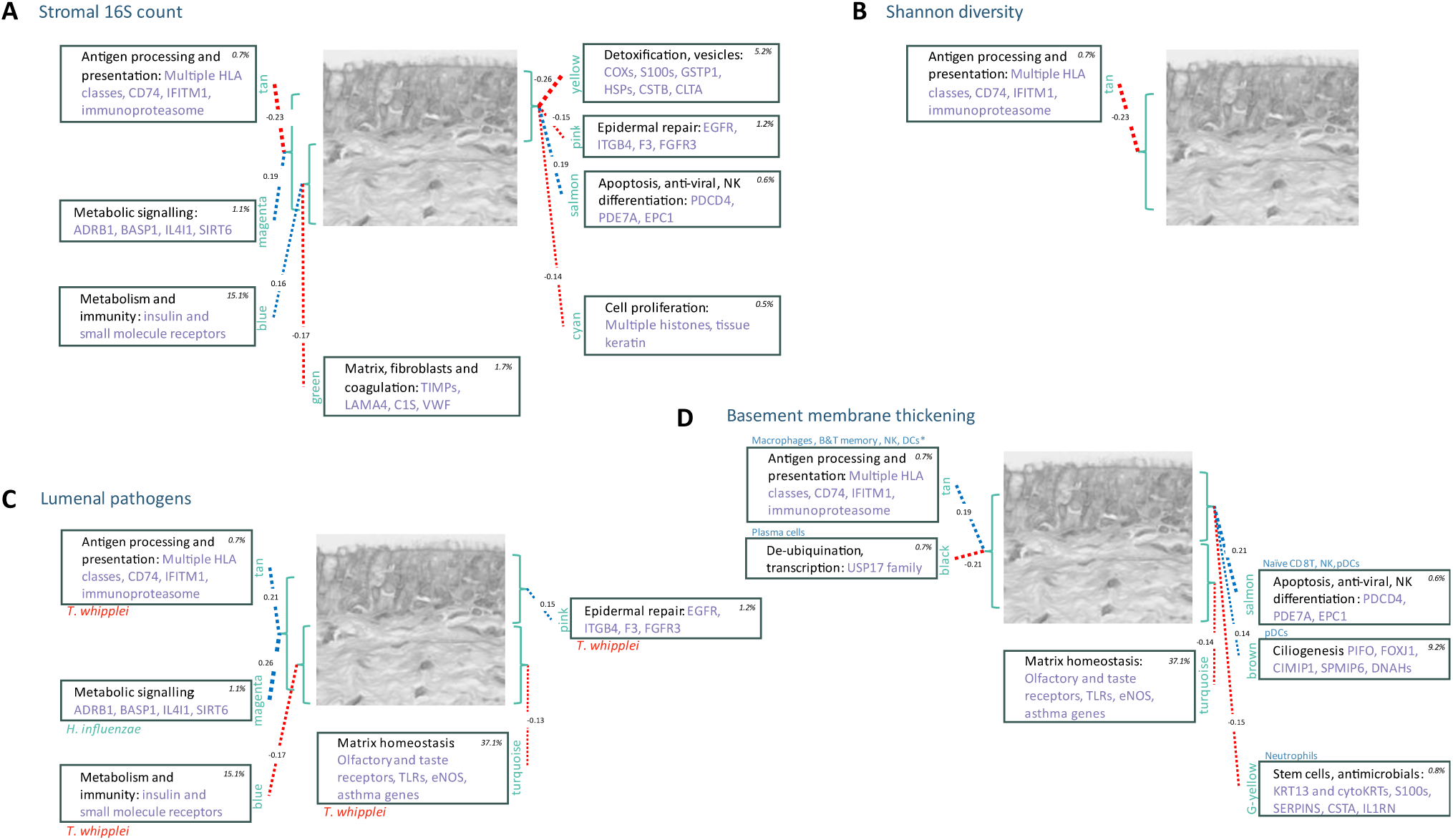
Phenotypic association with expression modules. Summary of associations between transcript networks and central phenotypes (derived from Fig. 3A). (**A**) Module associations with 16S probe bacterial counts from the biopsy stroma. Blue lines indicate positive associations, red lines negative. Line weight is proportional to the correlation coefficient. (**B**) Module associations to Shannon diversity of lumenal microbiome. (**C**) Associations to lumenal *T. whipplei* or *H. influenzae* abundances. (**D**) Module associations to basement membrane thickening, scored from H&E sections.

By contrast, the lumenal abundance of the pathogen *T. whipplei* was positively related to antigen processing (tan module) and epidermal repair (pink), whilst negatively correlated with blue module metabolism and immunity and turquoise module matrix homeostasis (Fig. 4C). Lumenal *H. influenzae* were associated with enhanced metabolic signaling and the magenta module, suggesting selective responses to different pathogens.

Basement membrane thickening, confirmed as a central pathologic marker of asthma (*1*) in our data, was positively associated with the modules mediating antigen presentation, apoptosis and ciliogenesis (Fig. 4D), consistent with upregulation of these processes in asthmatics. A negative relationship was observed to the modules with putative functions in membrane and matrix homeostasis (green-yellow, turquoise and black).

### Therapies and biomarkers

Current asthma treatments are structured progressively according to severity of symptoms (*26*), with biomarkers recommended to guide biologic therapy in severe disease (*27*). Biomarkers in common practice are blood eosinophils counts and exhaled nitrous oxide (FeNO) levels (*27*).

However, in our subjects, peripheral eosinophil counts independently predicted only 6% of the variation between asthma and non-asthma, less than airway basement membrane abnormalities (16%) or Shannon diversity in airway brushings (21%) (Table S3). FeNO is used as a marker for inflammation, but we found endothelial NO synthetase (*NOS3*, *eNOS*) and the NO receptor *GUCY1A1* were strongly stromal, (Fig. 3C) distant from basement membrane pathology.

Therapies for mild-moderate asthma include glucocorticoids, beta-adrenergic agonists, and antihistamines. The glucocorticoid receptor (*NR3C1*) was stromal (r=0.45, P<10^-9^) (Fig. 3C), as were histamine receptors (*HRH1*, *HRH2*, *HRH3* and *NTSR2*: r=0.39-0.54, *P*<10^-6^), whereas the beta-adrenergic receptor was not localized. Each of these receptors would be accessible with diffusible small molecule therapies, in line with current clinical practice.

Asthma GWAS susceptibility genes point to effective therapeutic choices (*4, 28*). *IL33* is a GWAS top hit which showed strong epithelial localization with CKoS (r=-0.47, P<10^-11^) (Fig. 3C). IL33 is a major alarmin and current target for biologic therapy, but its epithelial presence may suggest inhaled rather than systemic administration of antibody-based agents to be optimal.

*ORMDL3* is the major childhood asthma locus (*4, 29*), strongly predisposing to human rhinovirus (HRV)-induced asthma attacks (*30*) and regulating sphingolipid synthesis. It was strongly located to epithelium (r=-0.42, P<10^-9^), suggesting it may be a target for inhaled small molecule intervention to prevent or ameliorate the initial stages of acute asthma.

The results indicate improvements in biomarker design that take into account the spatial distribution of disease processes are possible. These might include the abundance of indicator species as a surrogate for diversity, and, although requiring vigilance, airway biopsy in severe cases has the potential to expand the utility and range of biologic and small molecule therapies.

## Discussion

The presence of IMCBs within the airway lamina propria is consistent with pathological, microbiological and transcriptomic elements of our study. IMCB abundances correlated positively lumenal bacterial diversity and negatively with pathogen counts (*T. whipplei* and *H. influenzae*). The correlation with lumenal diversity suggests a subset of commensals are resident within the airway mucosa, as has been established with bacterial communities within normal human skin (*31, 32*). Intramural bacteria are also visible in bowel mucosa (*33*), and their diversity within the stroma of lung cancer predicts better outcomes (*34*). IMCBs may therefore provide direct access for healthy microbial communities to influence inflammatory tone.

Our results should be interpreted in light of the study limitations. Our subjects were adults with a range of asthma severity and ages of onset, but not all asthma variants (particularly neutrophil or *H. influenzae* dominated) would have been included. The airway mucosa covers several m^2^ of area and our biopsies are of the order of mm^2^. Although some biopsies were too small or damaged to analyse histologically, there was no evidence of systemic dropout related to disease status or microbial profile. Lumenal microbial communities were accessed by sample brushings from upper and lower lobe bronchi, whereas biopsies were taken from the middle lobe. However, we have shown previously that regional differences in commensal pathobiont abundances in these subjects were modest (*35*), and were not significantly associated here with biopsy findings.

Our focus has been to understand airway bacterial dysbiosis, and our study did not address events that accompany asthma attacks or the major effects of HRV infection (*19, 20*). Airway biopsies are hazardous in the presence of bronchospasm, and so information on exacerbations may need to come from controlled experimental models (*36*).

Care needs to be taken in attributing cause or effect from positive or negative eigenvector correlations with disease phenotypes. Individual transcript localization may differ from that of the module eigengene, which is derived from all module members, and neighboring networks on the module phylogenetic tree may share gene content (Fig. S2, Table S4). Nevertheless, modules identified by WGCNA were bound by strong and highly significant correlations, with the MM of network hubs typically >0.85 (or <-0.85) and *P*<10^-54^, so that it is possible to draw confident insights into the integrated spatial organisation of airway mucosal immunity.

For example, multiple components of classical immune pathways for antigen processing and presentation were observed in the tan module. Antigen surveillance was shown to involve co-ordinated expression of multiple HLA classes, to be less active with abundant IMCBs yet stimulated in the presence of lumenal *T. whipplei* (Fig. 4C). Additionally, many canonical adaptive immunity elements encoding the T-cell/B-cell synapse and IgG production were present in the stromal purple module. Strong abundance signals from naïve CD4+ and CD8+ T-cells perhaps identify a stockpile of uncommitted T cells for the antigen specific machinery.

The turquoise module, centred in the lamina propria and containing 36% of detectable transcripts, extends understanding of innate immune integration. Capacities for pathogen and metabolite recognition are indicated by a membership that includes *TLR5*, *TLR6* and *TLR9* as well as numerous olfactory and taste receptors (*37*). *ORMDL3*, *STAT6* and *IL33* are major asthma susceptibility genes with prominent roles in innate antiviral immunity (*30, 38, 39*), and other module genes regulate synthesis of the potent anti-viral metabolite NO (*40*). Future investigation of stromal responses to acute viral infections may uncover mechanisms to explain why usually mild HRV infections have profound effects in patients with asthmatic bacterial dysbiosis.

Microbial diversity is known to contribute to microbiome community stability and pathogen exclusion. Whole-community airway microbiome replacement could contribute to asthma care, as faecal transplant does in dysbiotic bowel diseases. However, more selective therapies, including engineered organisms, are possible if resident IMCBs are a conserved subset of surface communities (*32*). The recent isolation and genome-sequencing of a broad range of commensals from healthy airways (*35*) may therefore provide a founding resource to develop microbial therapies for airway diseases.

## Supporting information

Subject details

Correlation analyses

Stepwise predictions oaf asthma status

Membership of WGCNA modules

Visual scores for 16S foci

## Non-author contributions

We are very grateful to the patients and healthy subjects who consented to airway bronchoscopy and biopsy. Mark Andrews, Joy Creaser-Thomas, and Robin Ghosal contributed to bronchoscopies in Welsh Centres. Lorraine Lawrence managed biopsy sections with great skill. We thank the team of application scientists at Nanostring for their generous advice in optimizing their applications. We are grateful to Cecila Johansson for her critical review of our immunological findings.

## Author contributions

Conceptualization: MFM, MJC, CB, LC, KFC, JMH, WOCC

Methodology: MFM, MJC, LC, TN, JLR, YR, LL

Investigation: TN, CMB, CB, CL, KL, GD, JI-H, RA, JLR, KFC, OO’C, JMB, MFM, WOCC

Visualization: TN, JLR, RA

Funding acquisition: WOCC, MFM, GML

Project administration: AMG

Supervision: MFM, WOCC

Writing – original draft: WOCC, initial review & editing: MFM, TN, MJC.

## Data and materials availability

Images will be available in image data resource: http://idr.openmicroscopy.org/about/ Clinical and metadata will be available:

Spatial whole transcriptome digital spatial profiling (GeoMX) data generated in this study has been deposited as unfiltered and filtered R objects as well as raw data in the Gene Expression Omnibus (GEO) database under accession code GSEXXXXX. Whole transcriptome digital spatial profiling data are also available as a supplement to the manuscript in the formats of raw expression (DCC and PKC files) and processed expression (filtered for targets detected in at least 5% of the ROIs and Q3-normalized) data along with the annotation file for each ROI.

All data analysis scripts will be available online at https://zenodo.org/records/xxx

## Supplementary Materials

### Materials and Methods

#### Clinical protocols

Samples were collected through a multicentre, cross-sectional study of asthmatic adults and healthy controls (the microbial pathology of asthma study (Celtic Fire, CELF). Participants were recruited from Connolly Hospital, Dublin; The Royal Brompton Hospital, London; and Swansea University Medical School, Swansea. Ethical approval was granted by the London-Stanmore Research Ethics Committee (reference 14/LO/2063). All subjects provided written informed consent. Subject groups were: healthy subjects (non-smokers and current smokers; asthmatic patients taking short-acting beta agonists only (BTS Step 1)(*26*); asthmatics on moderate dose of inhaled corticosteroid (ICS) ± long-acting β-agonist LABA (BTS Step 2/3); asthmatics on high dose ICS (ICS dose >=1600 µg/day) + LABA ± other controllers (theophyllines, LTRA, LAMA) (BTS Step 4); and asthmatics on high dose ICS (ICS dose >=1600 µg/day) + LABA ± other controllers + oral prednisolone ± anti-IgE treatment (BTS Step 5). Severe asthma was defined as BTS step 4 or 5. Exclusion criteria were: Asthmatic subjects were non-smokers or ex-smokers with < 5 pack-years smoking; BMI>35; diagnosis of rheumatoid arthritis, allergic bronchopulmonary aspergillosis, or Churg-Strauss syndrome; drug therapy with beta-blockers, ACE inhibitors, anti-asthma immune modulators other than steroids; antibiotics within 4 weeks of study; acute exacerbations of asthma within past 4 weeks; history of an upper or lower respiratory infection (including common cold) within 4 weeks of baseline assessments; confounding occupations (such as baking); and significant vocal cord disorder.

A posterior oro-pharyngeal (ptOP) swab was taken from each participant immediately before the bronchoscopy commenced. During bronchoscopy, two bronchial brushings were taken from the left lower lobe (LLL) of each subject. If tolerated, two further brushes were taken from the left upper lobe (LUL).

Biopsies taken from the sub-segmental bronchi in the right middle lobe (RML) were shaken or coaxed from the end of the biopsy forceps into pre-labelled tubes. Samples for histology and digital spatial pathology were placed into 4% formalin and kept for subsequent paraffin embedding and H&E staining. Freezing was avoided.

All non-biopsy samples were stored at -80°C within 1 hour of collection. Those harvested at The Royal Brompton Hospital were transported stored directly to the Asmarley Centre for Genomic Medicine (ACGM) at the same site. Samples at other sites were stored locally at -80°C for a maximum of 6 months prior to transport to the ACGM on dry ice.

### DNA extraction and quantification

Microbial DNA extraction from brushing was carried out using a hexadecyltrimethylammonium bromide (CTAB) and bead-beating double extraction using phase lock tubes, as outlined in Cuthbertson *et al* 2020 (Protocols.io). DNA was stored at -20°C until processing. Microbial DNA quantification was carried out using a SYBR green 16S rRNA gene qPCR(*41*).

Within a Class 2 biological safety cabinet, each bronchial brushing was transferred directly into an LME tube. To control for contamination an empty LME tube (i.e., an extraction control) was added to each batch. The extraction control underwent the entire extraction process along with the samples. Eighteen randomly selected Scope Control Washes (SCWs) also underwent DNA extraction.

### Microbial 16S rRNA analyses

16S rRNA gene sequencing was performed on the Illumina MiSeq platform using dual barcode fusion primers and the V2 500 cycle sequencing kit. Sequencing was performed for the V4 region of the 16S rRNA gene as previously described(*22, 41*). Sampling and extraction controls, PCR negatives and mock communities were included on all sequencing runs. Sequences were quality trimmed to 200bp using trim-galore (Version 0.6.4) and joined with a maximum of 10% mismatch and a minimum of 150 base pair overlap using joined_paired_ends.py (Version 1.9.1). Data was quality checked using FASTX Toolkit (Version 0.0.14) prior to de-multiplexing.

Reads were dereplicated and open reference OTU clustering was performed in QIIME 2.0. Chimeric sequences were identified and removed, leaving borderline calls in the analysis. Phylogeny was aligned using mafft followed by consensus taxonomic classification. The Biom file, tre file and taxa identifications were exported for further analysis.

Processed data was transferred to R (Version 3.6.3) and uploaded into Phyloseq (Version 1.3). Reads unassigned or assigned to Archaea at the kingdom level were removed before further analysis along with reads identified as Chloroplast or Mitochondria. All OTUs with less than 20 reads (reads present in less than <2% of the samples (n = 1,174)) were removed from further analysis.

Contaminant OTUs were identified using Spearman’s correlation between bacterial biomass with number of reads per samples. OTUs were considered to be contaminants with a Benjamini-Hochberg corrected *P*-value of <0.05 and a correlation value of >0.2 and removed from further analysis. The “Prevalence” method in Decontam (Version 1.6) with a threshold of 0.1 identified a further 55 OTUs contaminant OTUs associated with negative controls. All OTUs identified were checked and found to be consistent with contamination(*42*).

OTU frequencies from 111 brushings from the left lower and 56 from the left upper lobe completed all quality checks and were passed to downstream analyses. OTU frequencies were transformed using log(x+1) prior to statistical analysis.

### Histology

Samples from 109 biopsies were cut and H&E stained and the glass-mounted slides scanned using the Hamamatsu NanoZoomer S360 Digital slide scanner and viewed with Hamamatsu NDP.view2 Image viewing software. Images were reviewed by an experienced lung histopathologist (RA) who scored the following features: Sample Adequacy (Yes/No); Bronchial epithelium (1=Normal, 2=Abnormal); Denudation (1/2); Epithelial hyperplasia (1/2); Goblet cell hyperplasia (1/2); Epithelial metaplasia (1/2); Basement Membrane (1/2): if abnormal, Thickened (1/2) and breached (1/2); Inflammation (judged on 10 high power fields) Absent (0) Present (sparse=1, confluent=2); and the presence of Macrophages (1/2); Giant cells/Granuloma (1/2); Lymphocytes (1/2); Plasma cells (1/2); Eosinophils (1/2); Neutrophils (1/2); and visible bacteria (1/2). Ninety-seven biopsies were adequate for interpretation: the remaining 12 were too small or damaged, or contained only cartilage or fibrin.

### GeoMx™ DSP with human WTA

GeoMx™ digital spatial profiling was performed with the human Whole Transcriptome Atlas (WTA) probe set with read-out by next-generation sequencing (Nanostring Technologies). Briefly, slides were mounted with 4 individual FFPE lung biopsy samples sectioned at 5 µm, dried overnight, and stored at 4°C in a desiccator until processing (within 2 weeks of sectioning). Slides were baked for an hour at 60°C, deparaffinized, and processed for detection of pan-eubacterial rRNA using RNAscope® Probe 16S-rRNA (Bio-Techne, 464461) and the Multiplex Fluorescent v2 Reagent Kit (Bio-Techne, 323100) following the manufacturer’s procedure. The r16S signal was developed using OPAL-620 fluorescent dye (Akoya, FP1495001KT), after which the slides were treated with a Horseradish Peroxidase Blocker for 15 min at 40°C, washed twice with wash buffer, and post-fixed in 10% neutral buffered formalin. Slides were then incubated overnight with the WTA probe set at 37°C in hybridization buffer, washed twice under stringent conditions (2XSSC/formamide, 25 min at 37°C), washed twice with 2XSSC, and blocked prior to immunofluorescence with PanCK-AlexaFluor-532, 1:100 (Novus, NBP2-33200AF532) together with SYTO13 DNA dye (Thermo, S7575) for 2 hrs in a humidity chamber. After washing 3 times in 2XSSC, slides were scanned on the GeoMx™ DSP instrument (at 20X) and WTA barcodes from PanCK+ or PanCK- (stroma) areas were released from defined regions-of-interest by UV light illumination and microfluidic collection. The UV-released barcodes from each of the 203 collected area-of-illuminations (AOIs) were PCR-amplified to add unique dual-index Illumina i5 and i7 indices and then bead-purified twice using AMPure XP beads (Beckman Coulter, A63881). Pooled libraries were sequenced on an Illumina NovaSeq S4 flowcell, generating 2.64B unique reads. FASTQ files were converted to DCC files and uploaded onto the GeoMx™ DSP for trimming, stitching, aligning, and deduplication.

Targets for which average counts across all segments were ≤10% of the counts for all probes to that gene and were not expressed above the Limit of Quantification in at least 1% of the AOIs (2 of 203 AOIs) were filtered out and data was upper-quartile normalized within each AOI for downstream analysis. In total, 12,487 targets (out of a total of 18,942) were included in the analysis. The results of downstream analyses were not sensitive to more or less stringent QC thresholds.

### Counting and validation of bacterial r16S

Images for each biopsy were generated by QuPath Version: 0.5.1 (*43*) with an inverted (white) background and printed on A4 paper with a standard magnification. Scoring 16S rRNA foci derived from ILO methodology for scoring pneumoconiosis in plain chest radiographs (*44*). After sorting images from least to most 16S rRNA foci in lamina propria, the abundance of foci in epithelium, basement membrane and lamina propria were each graded on a scale of 0-3 independently by two observers, blind to clinical or histological status (Fig. S1 for all images scored).

RNAscope fluorescence in-situ hybridization (ISH) was repeated as above for 9 biopsies, following the manufacturer’s procedure with the RNAscope® Probe 16S-rRNA (Bio-Techne, 464461). In parallel, a scrambled version of the same probe (ACD Cat no. 1144331-C1) was used on a serial section as a negative control. Slides were washed 3X in PBS and scanned on the GeoMx DSP instrument.

### Statistical analyses

Correlation and regression analyses were performed with IBM SPPS statistics v29.0.1.0 on a PC running Windows 10. WGCNA network analyses (*24*) was carried out in RStudio 2023.12.1 in the Windows desktop environment.

## Supplementary Figs

**Fig. S1. RNAscope staining for mucosal bacteria in airway biopsies.**

See file: Fig. S1. RNAscope staining for mucosal bacteria in airway biopsies.pdf

**Fig. S2. Negative controls with scrambled 16S rRNA probe.**

See file: Fig. S2. Negative controls with scrambled 16S rRNA probe.pdf

**Fig. S3. Network module eigengene phylogeny and correlations.**

See file: Fig. S3. Network module eigengene phylogeny and correlations.pdf

## Tables

**Table S1. Subject details and summary statistics.**

See file: Table S1 Subject details and summary statistics.pdf

**Table S2. Correlation analyses.**

See file: Table S2. Correlation analyses.pdf

**Table S3. Stepwise regression of asthma predictors.**

See file: Table S3. Stepwise regression of asthma predictors.pdf

**Table S4. Membership of WGCNA network modules.**

See file: Table S4. Membership of WGCNA network modules.xlsx (.pdf for initial submission)

**Table S5. DSP slides 16S visual scores for two readers.**

See file: Table S5. DSP slides 16S visual scoring.pdf

## Data

**Data S1. Clinical and microbial metadata.**

See file: Moffatt et al 16S mapfile with histology 16Score 2024_08_12.csv

All slides viewed in QuPath

Invert background

Turn off Cy3 to visualise bacteria in epithelium

Scale bar always 50µm

Score number of 16S objects 0-3

Three compartments:

Epithelium (if no epithelium score as missing)

Basement Membrane (BM)

Stroma (lamina propria)

Score sheets for both readers in Supplementary information:

Table S5. DSP slides 16S visual scoring

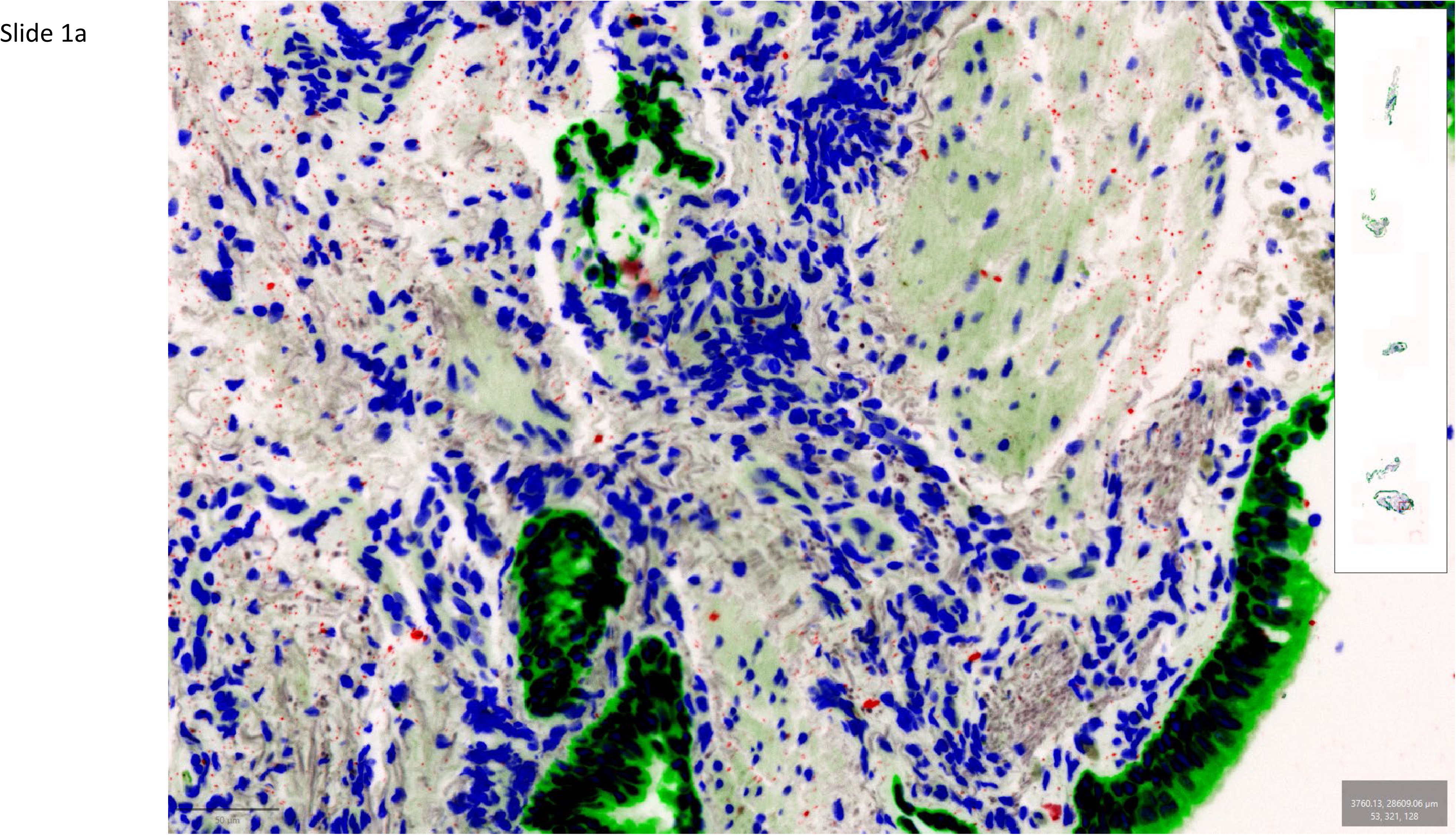
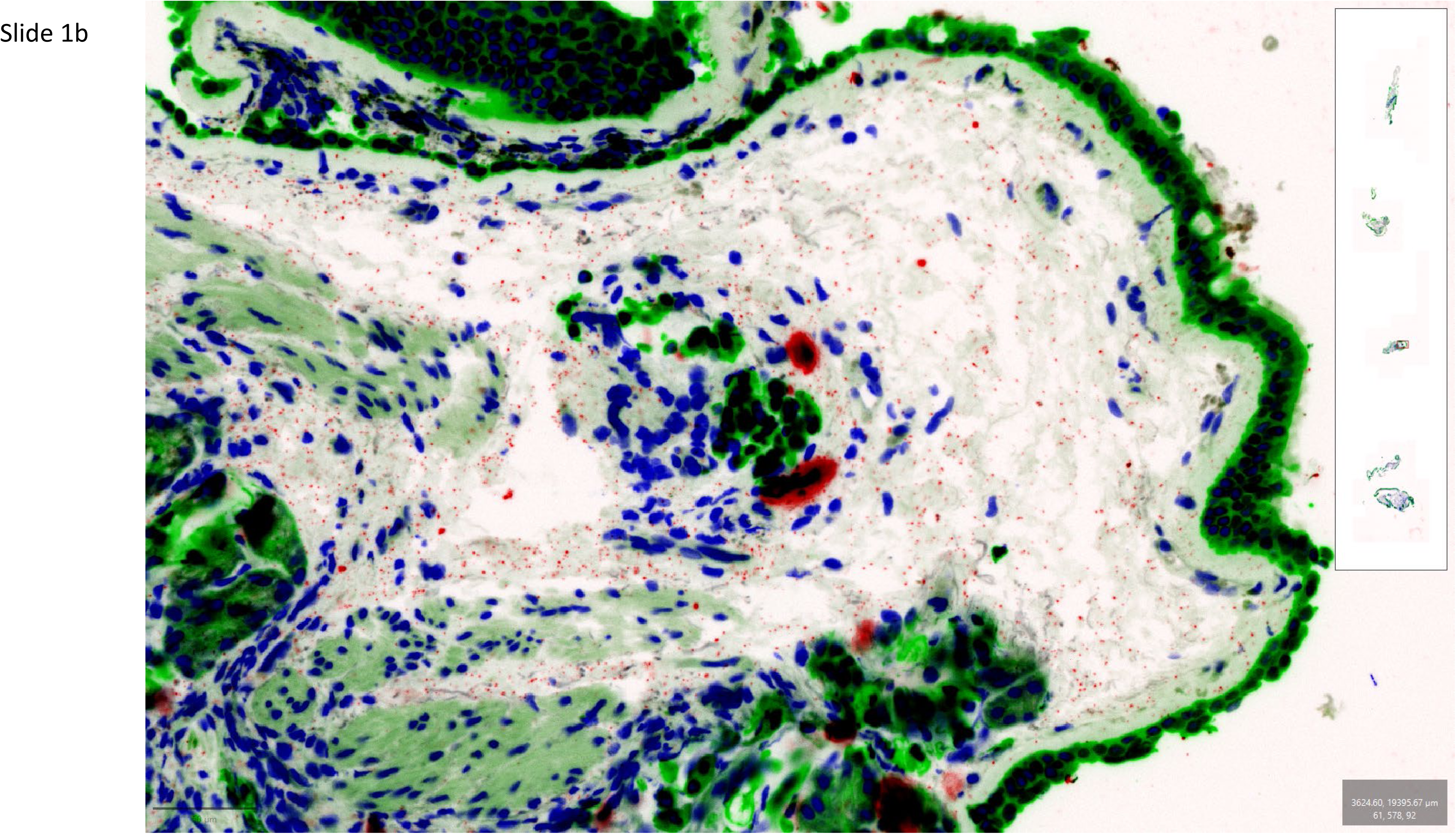
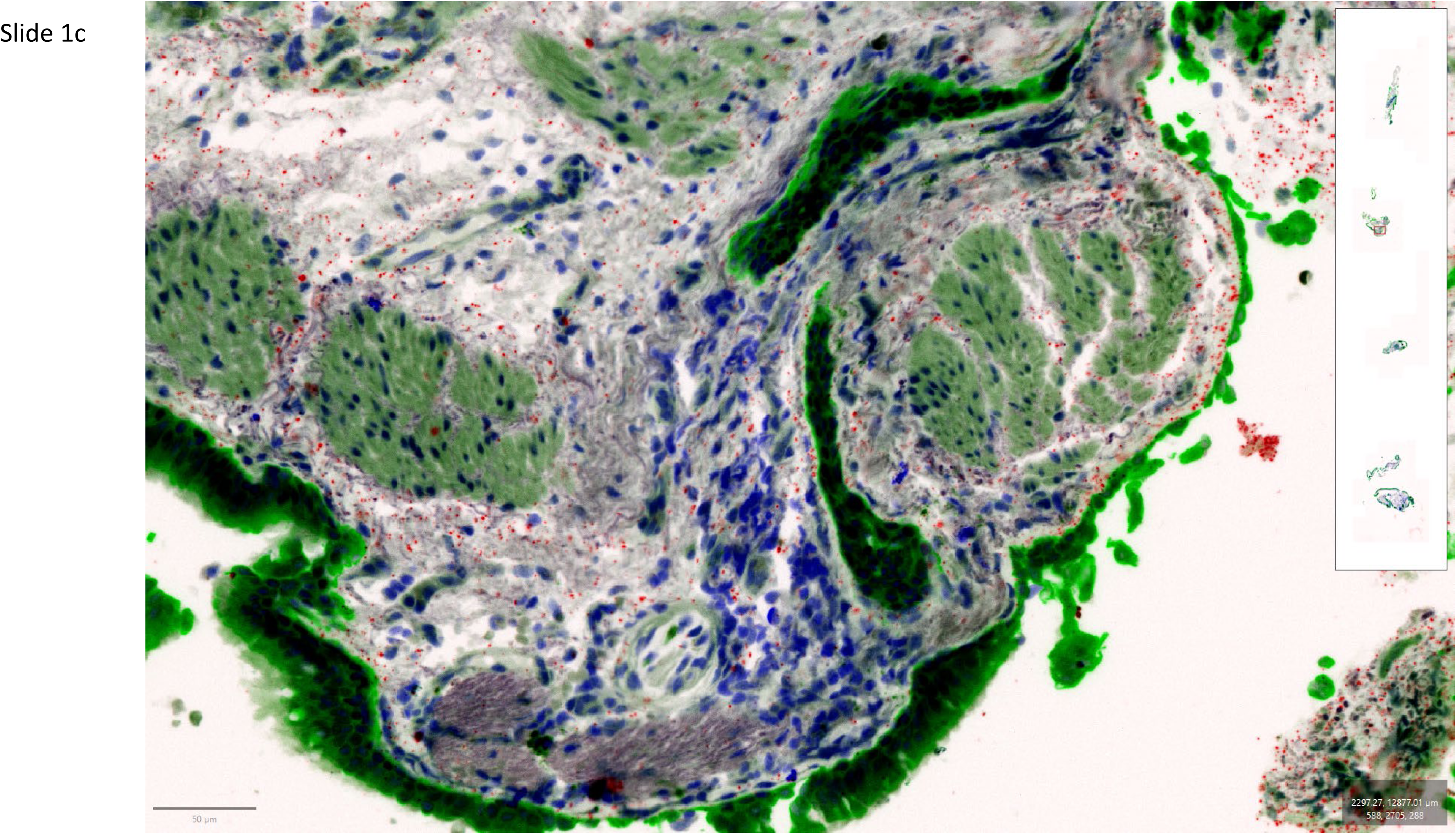
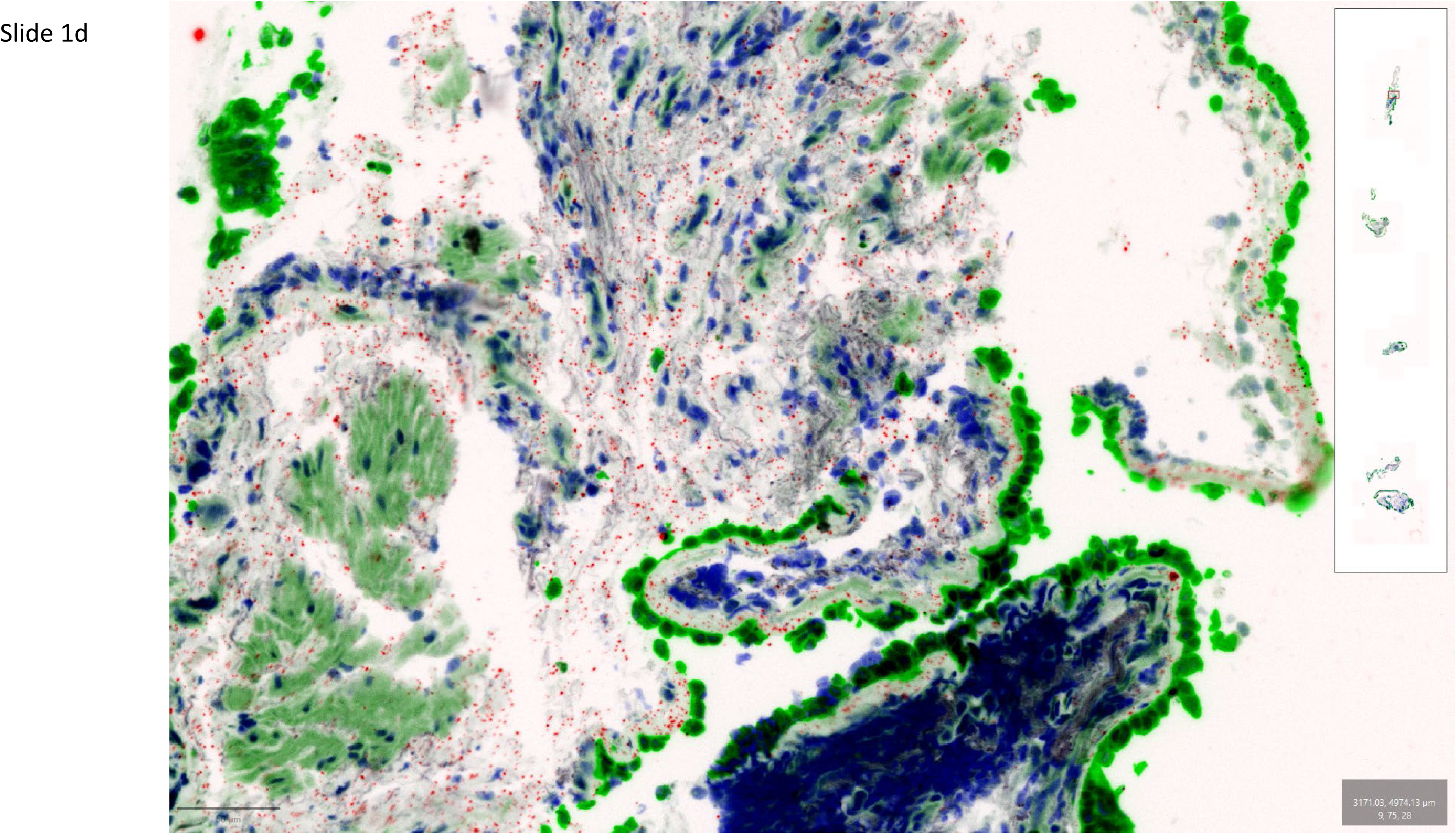
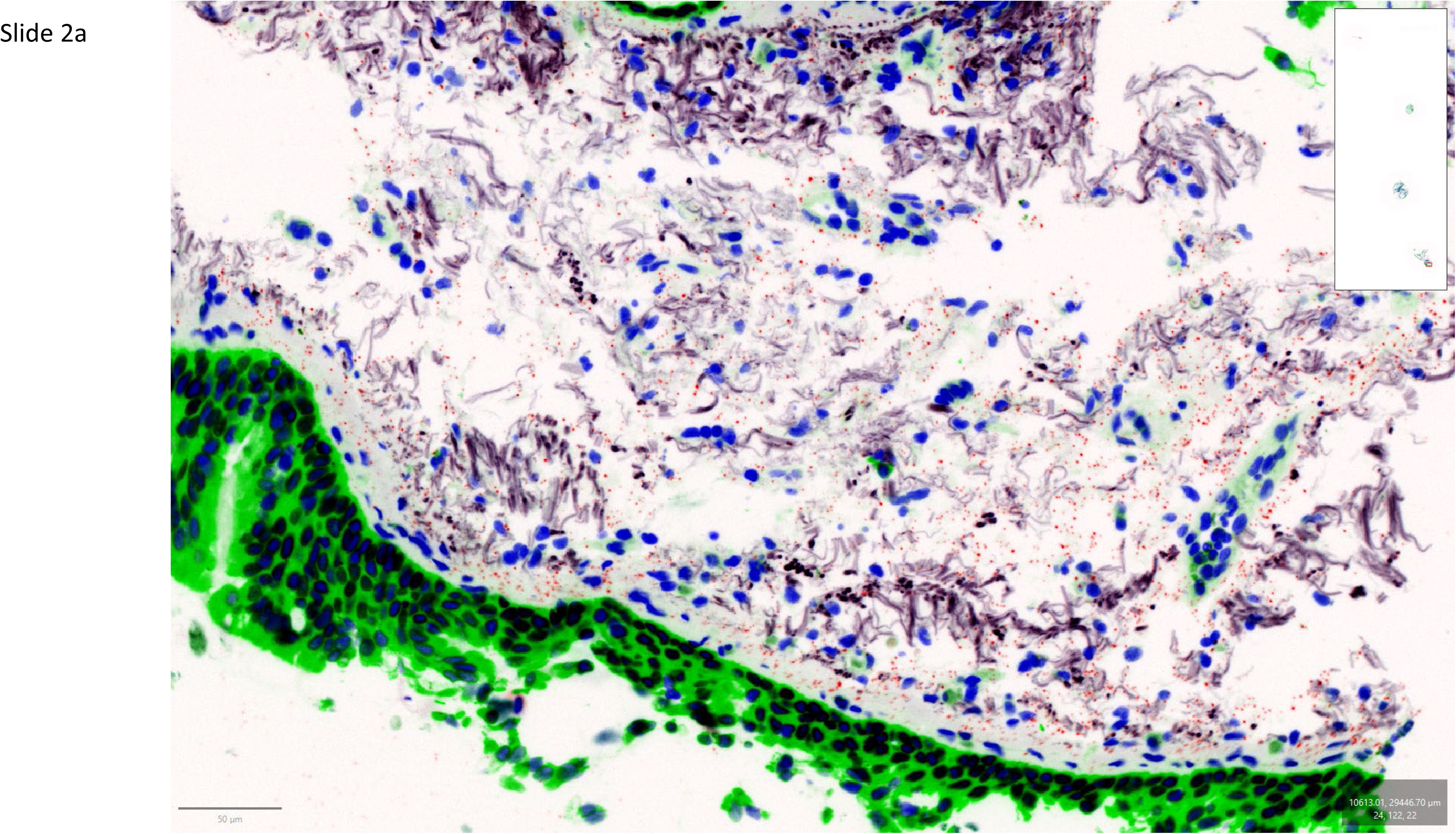
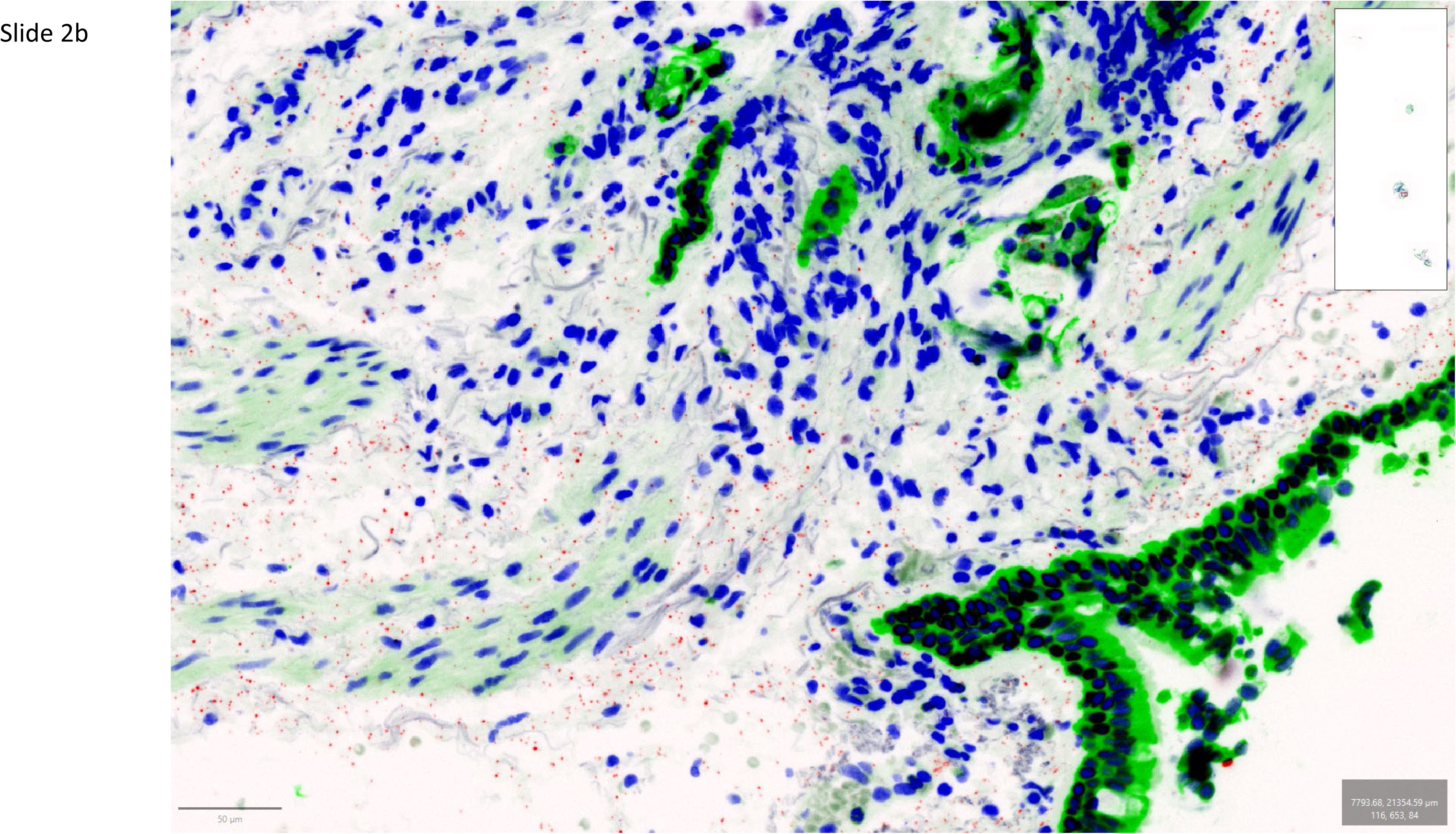
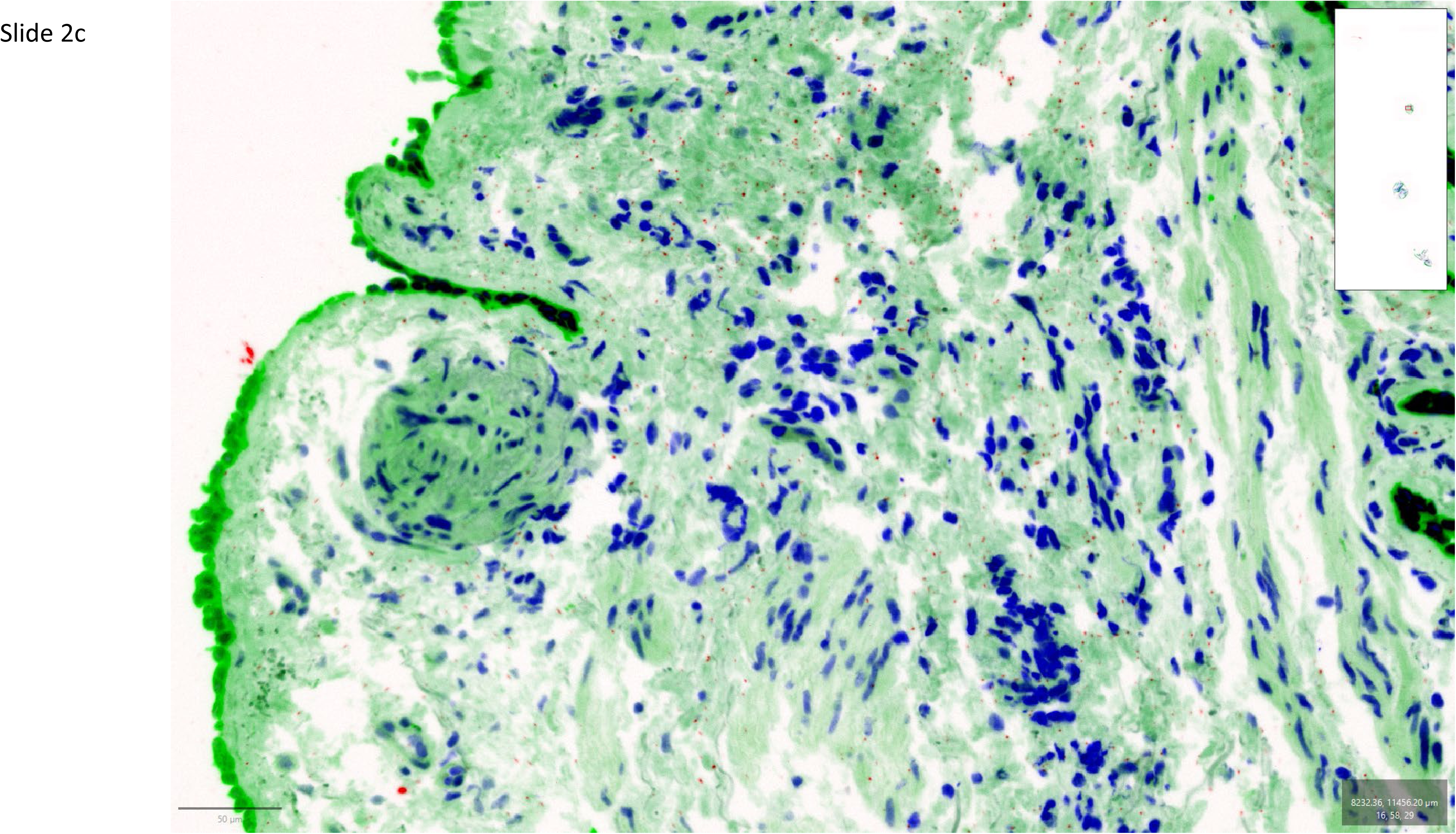
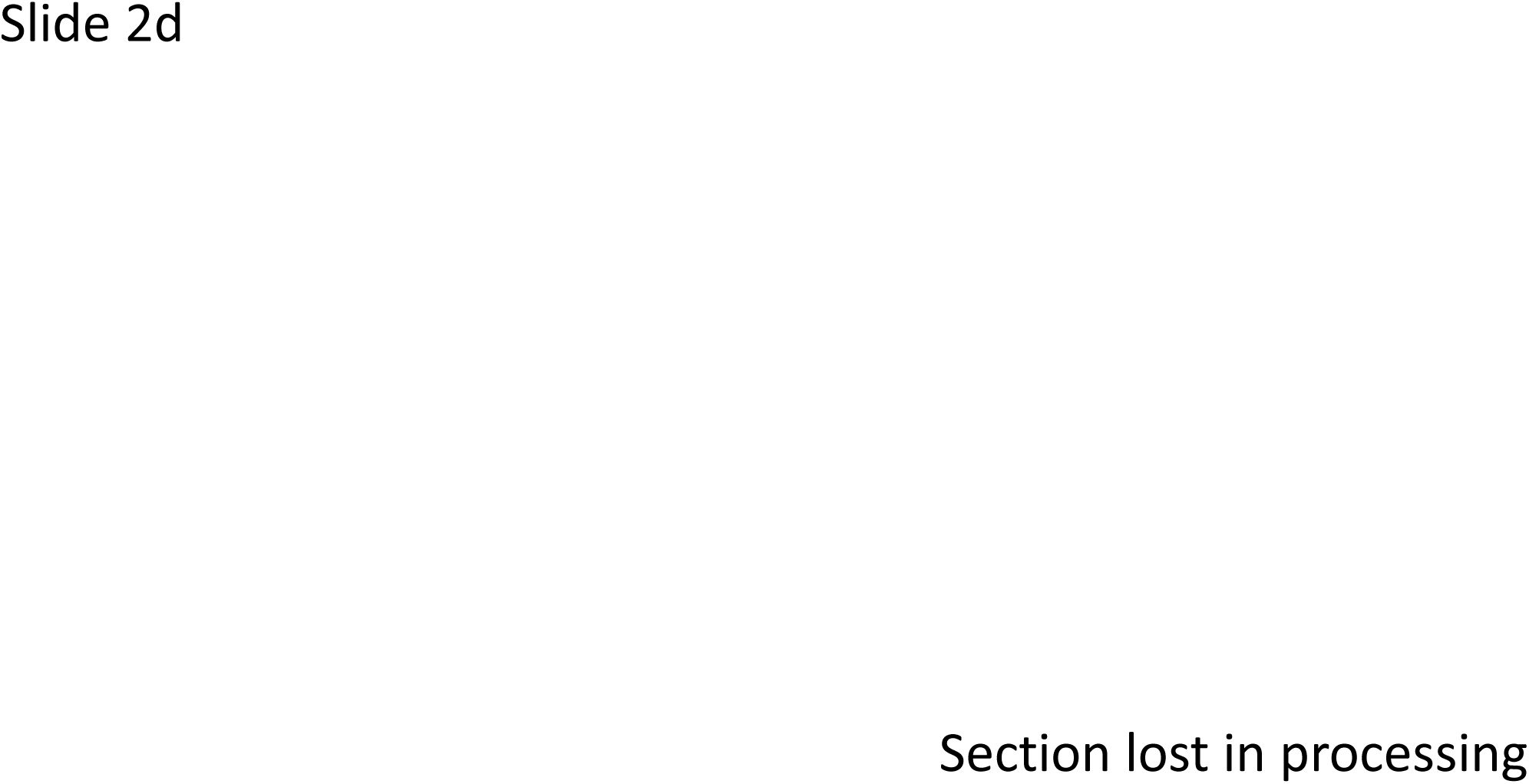

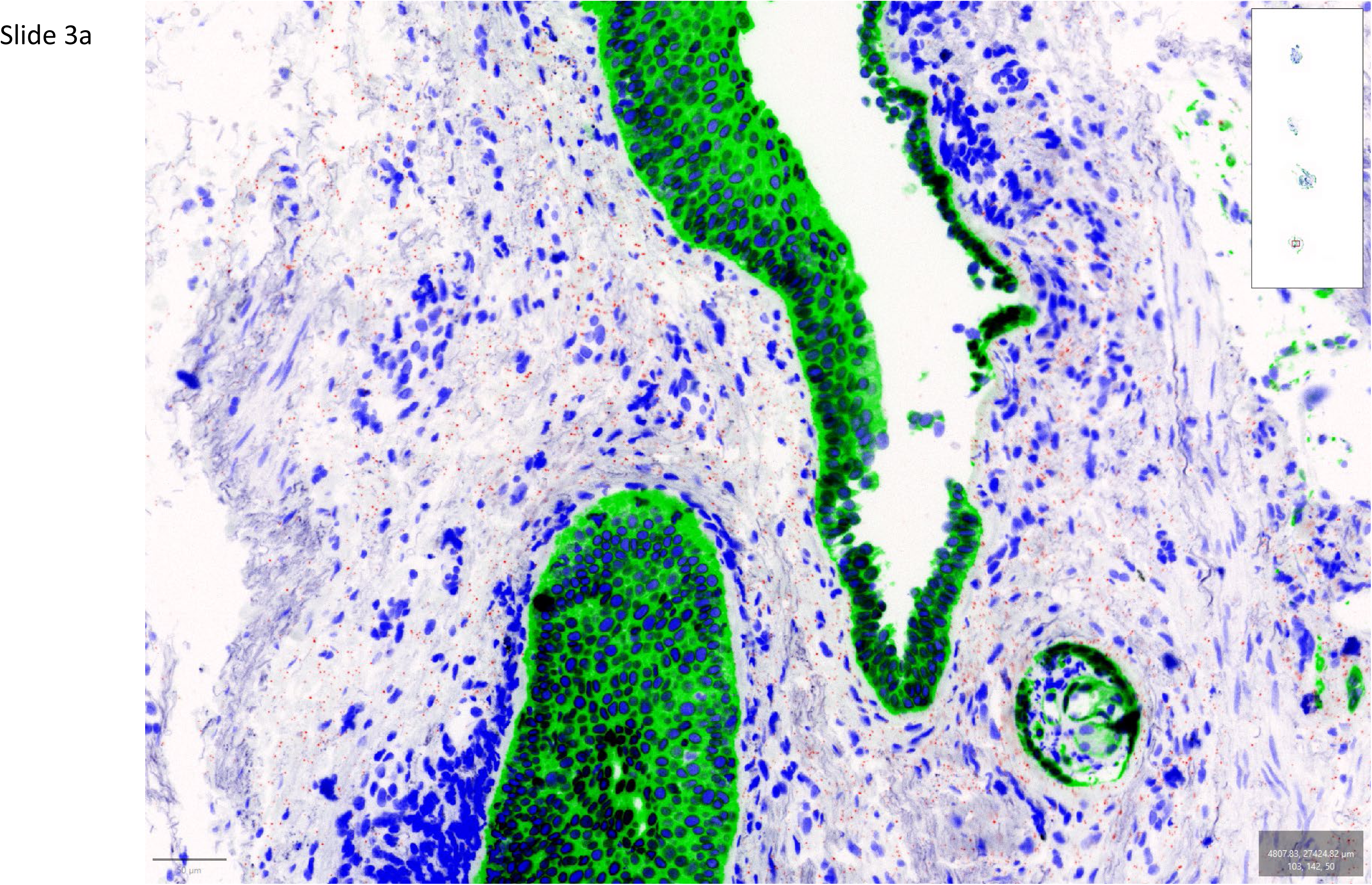
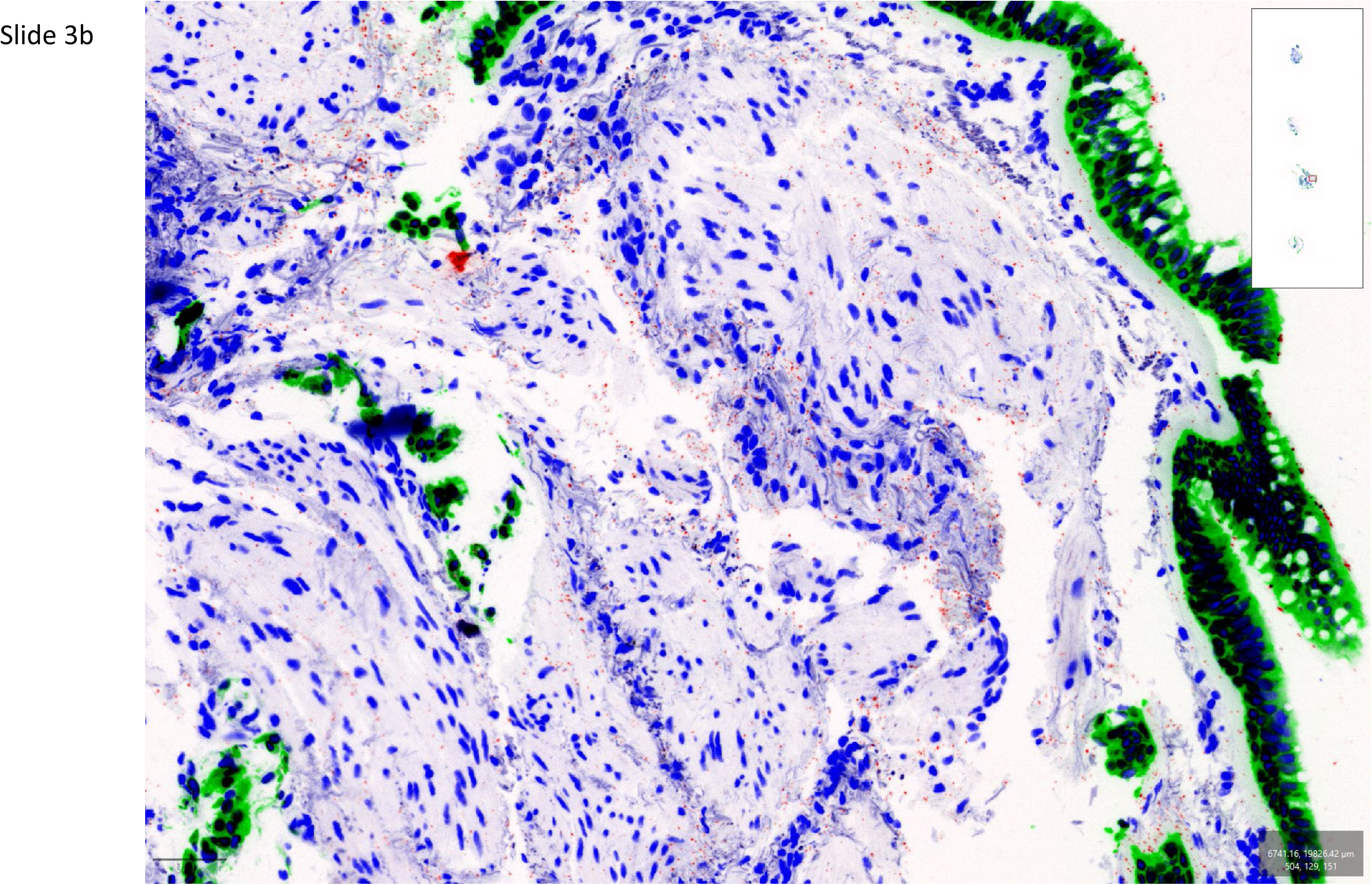
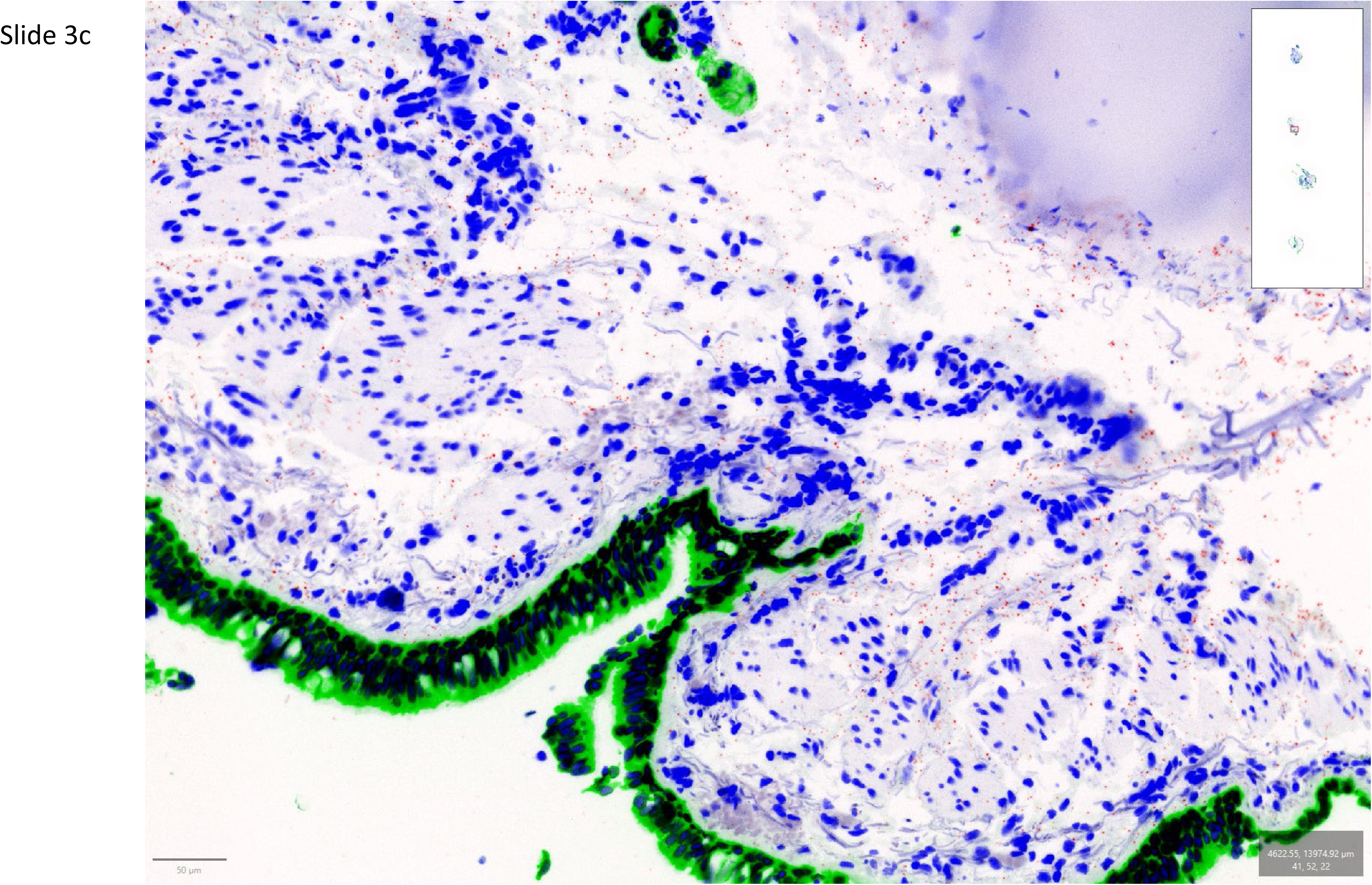
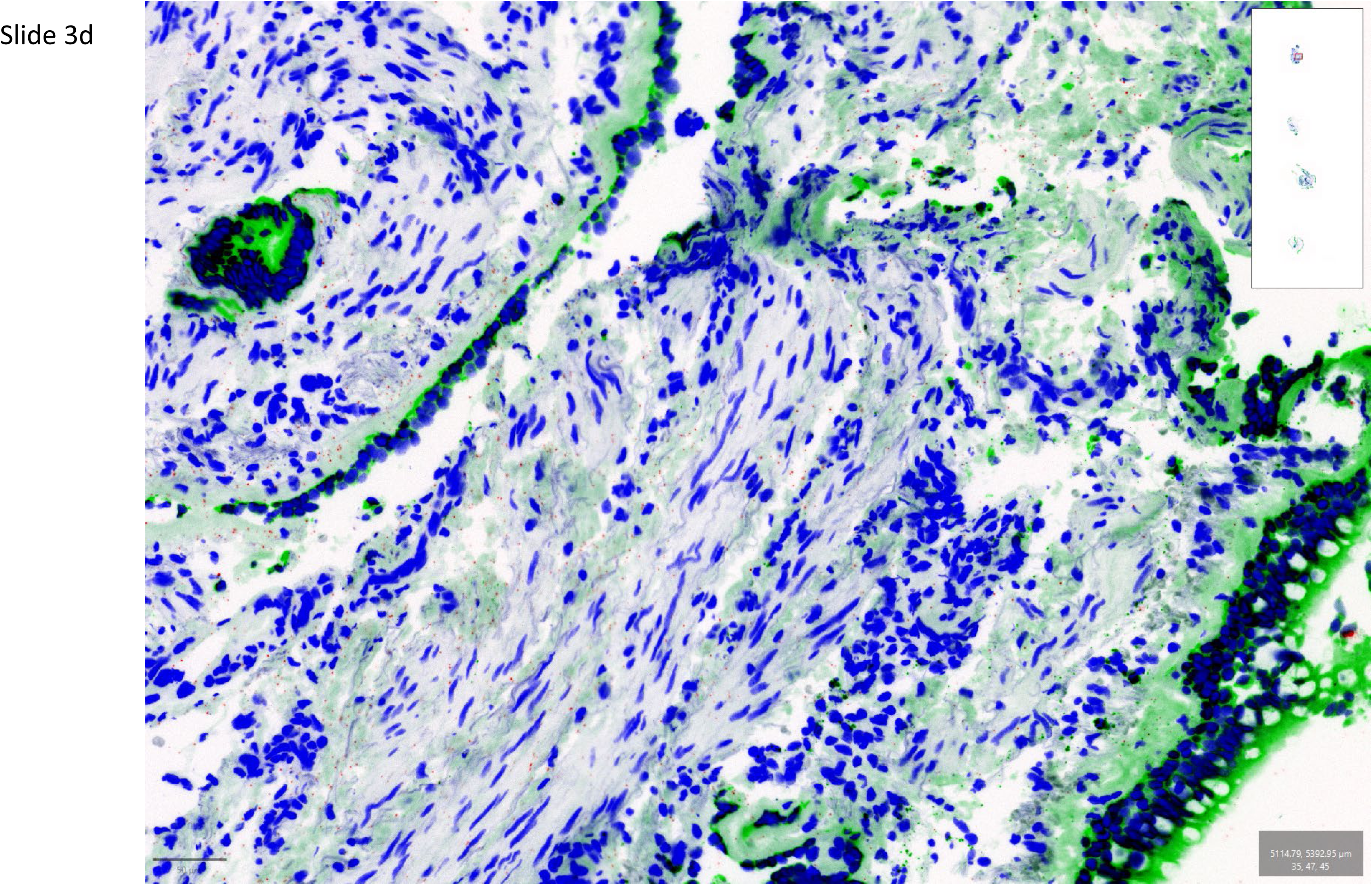
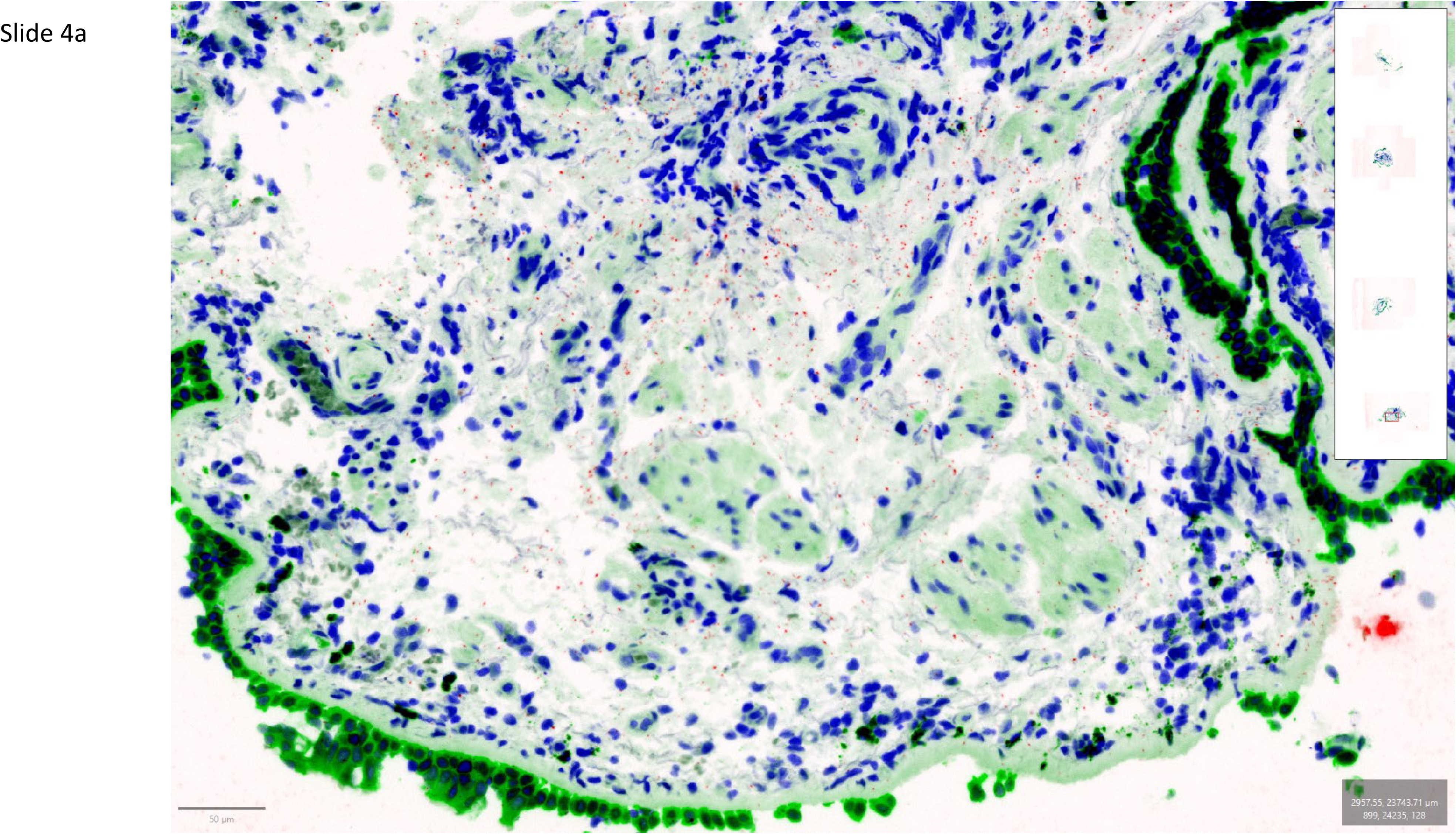
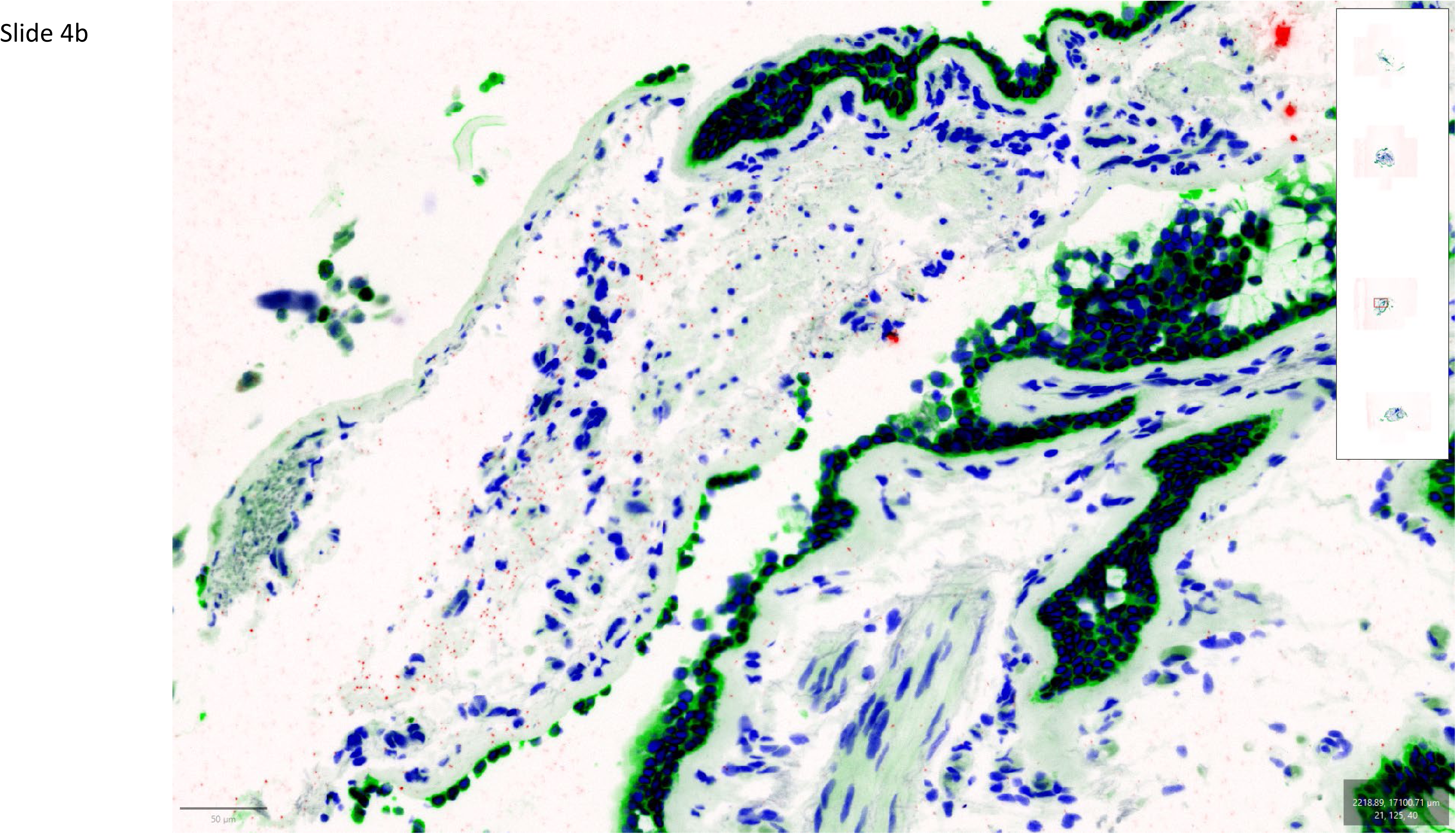
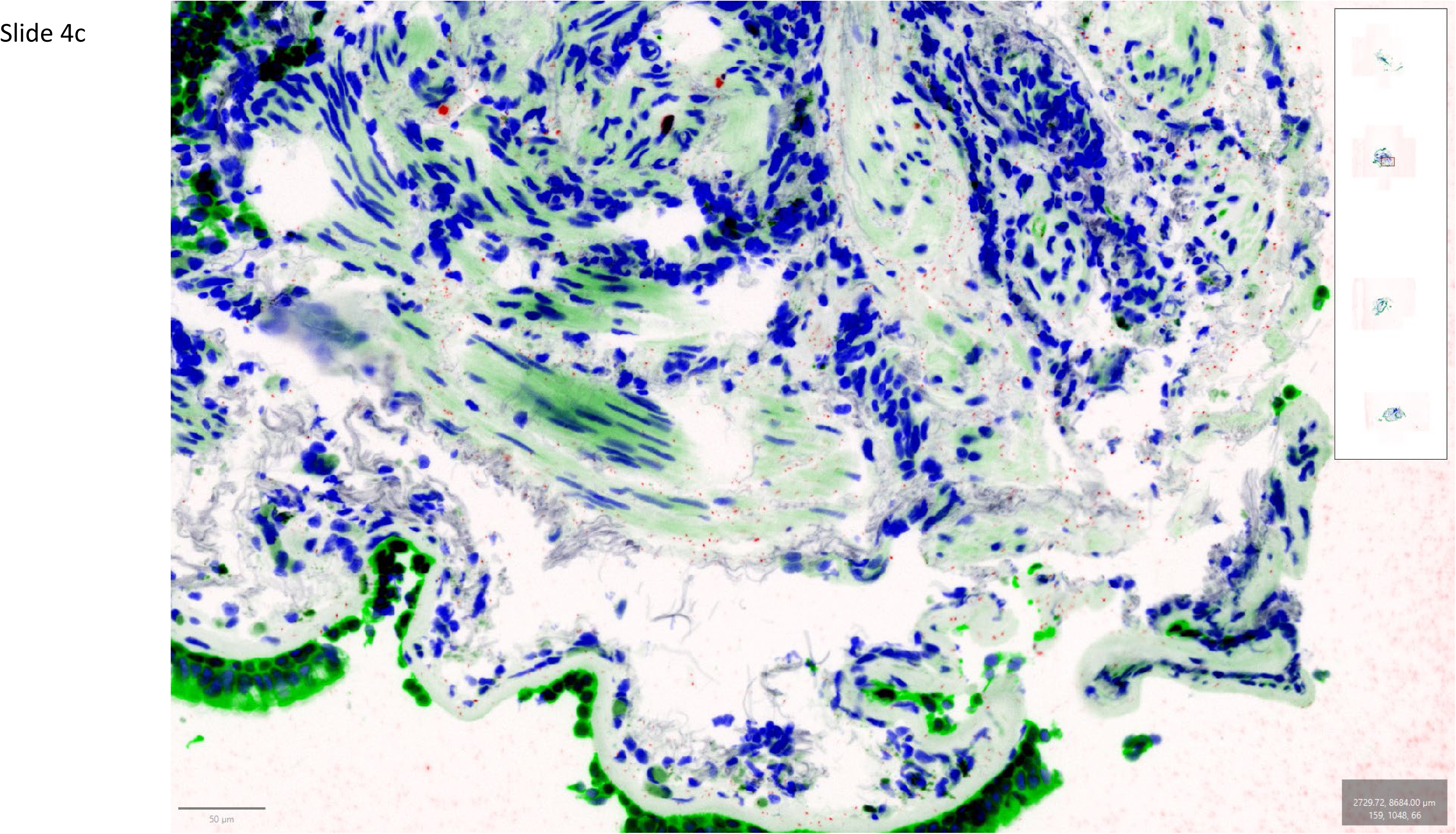
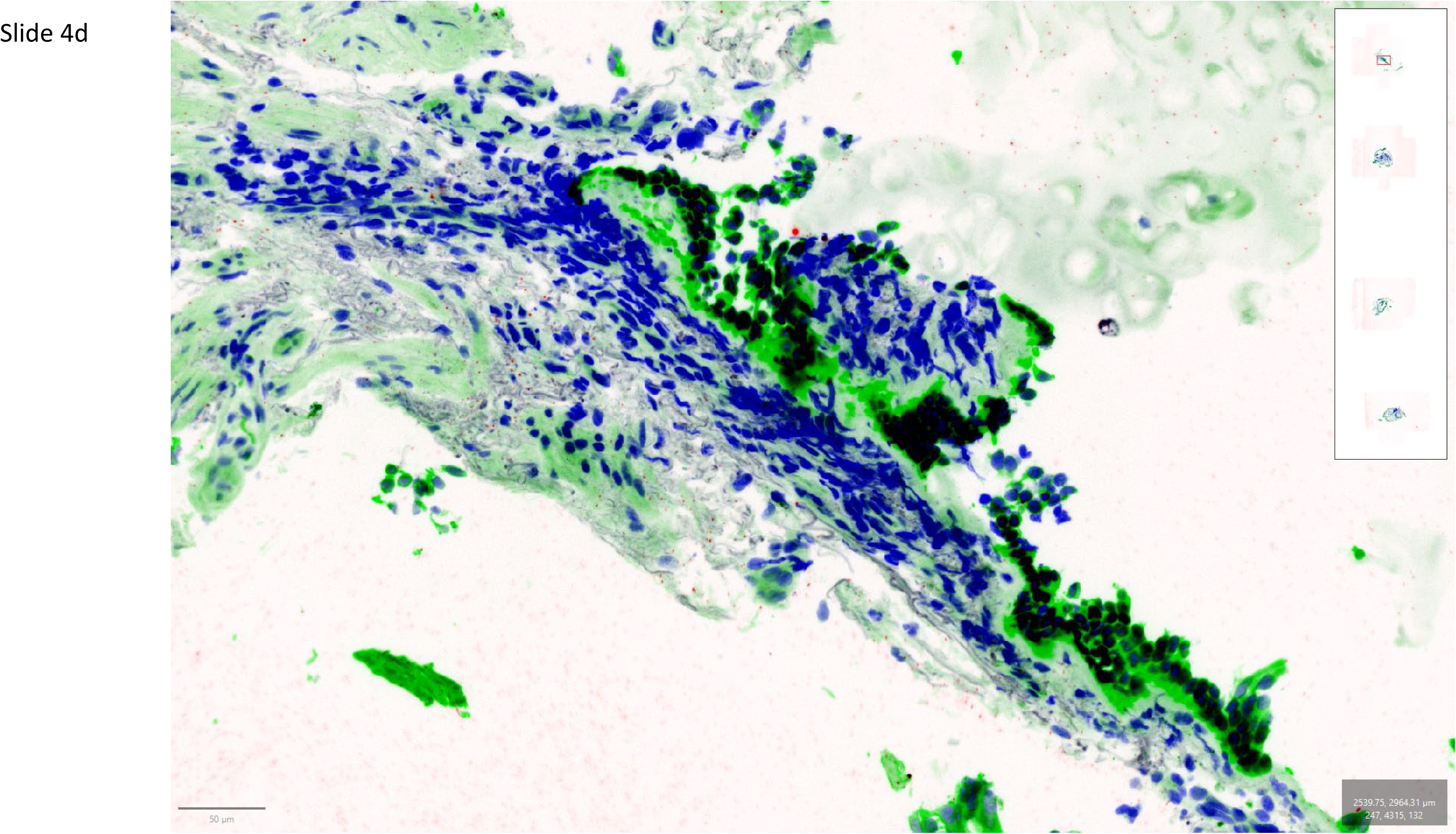
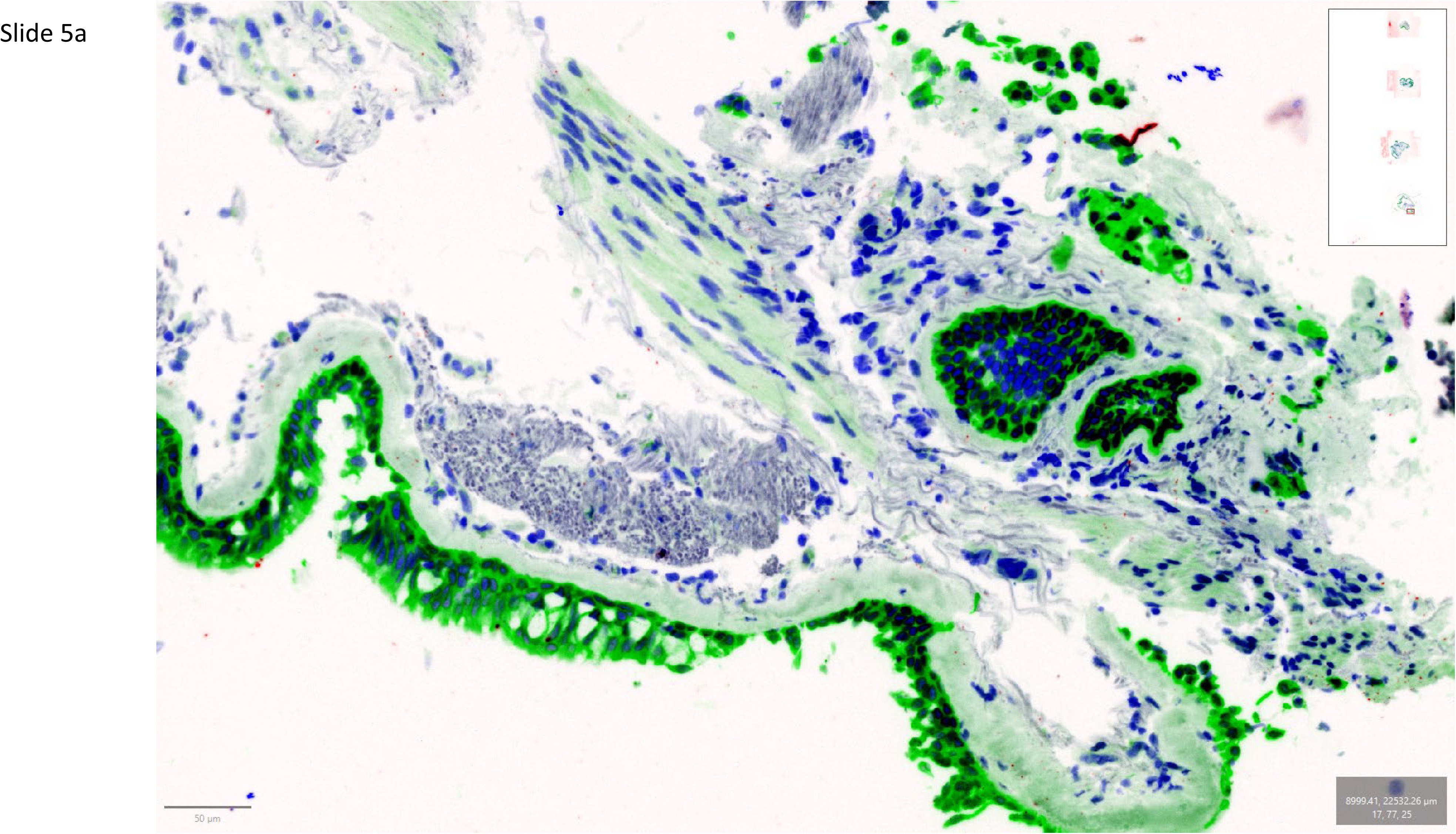
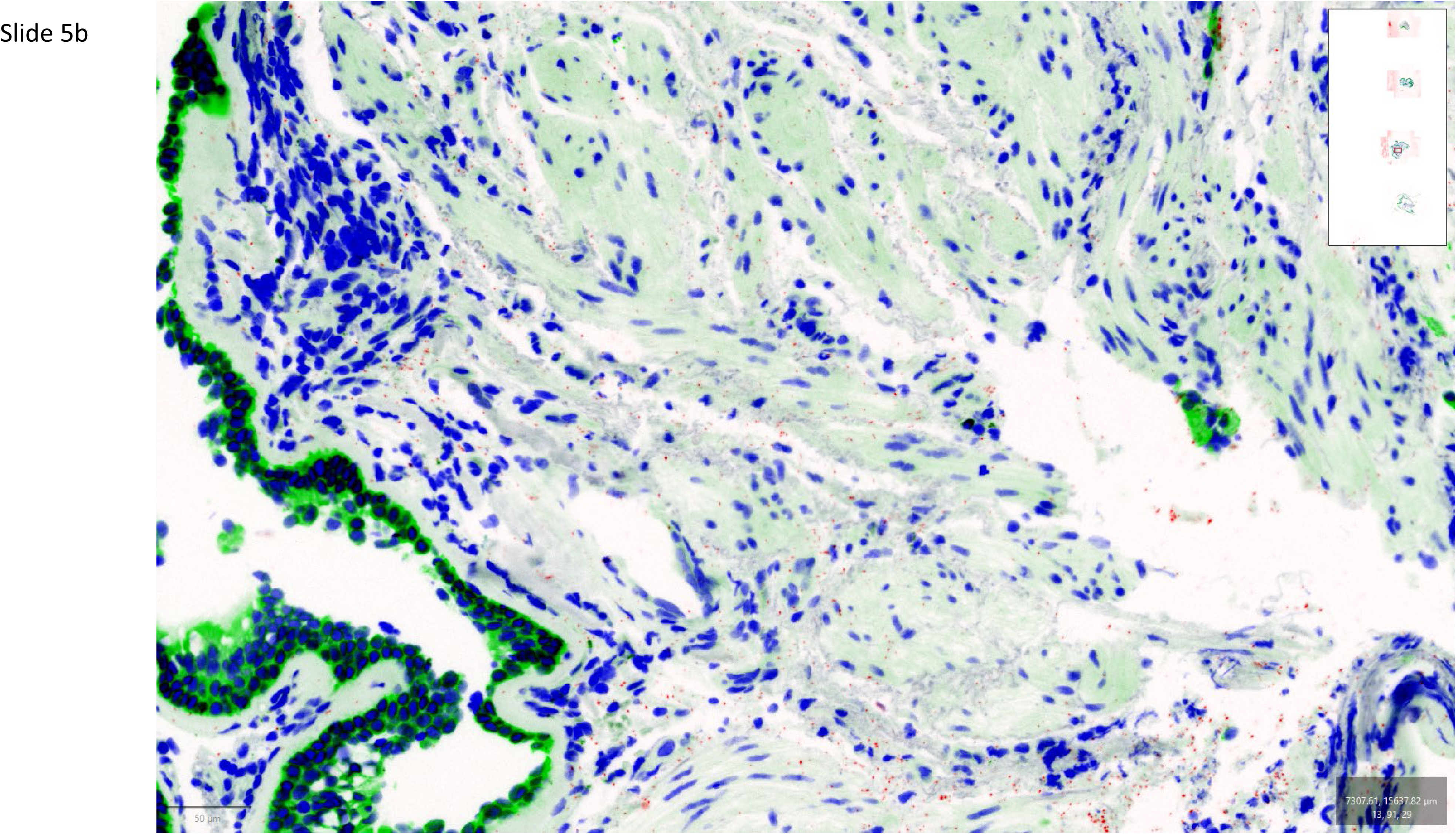
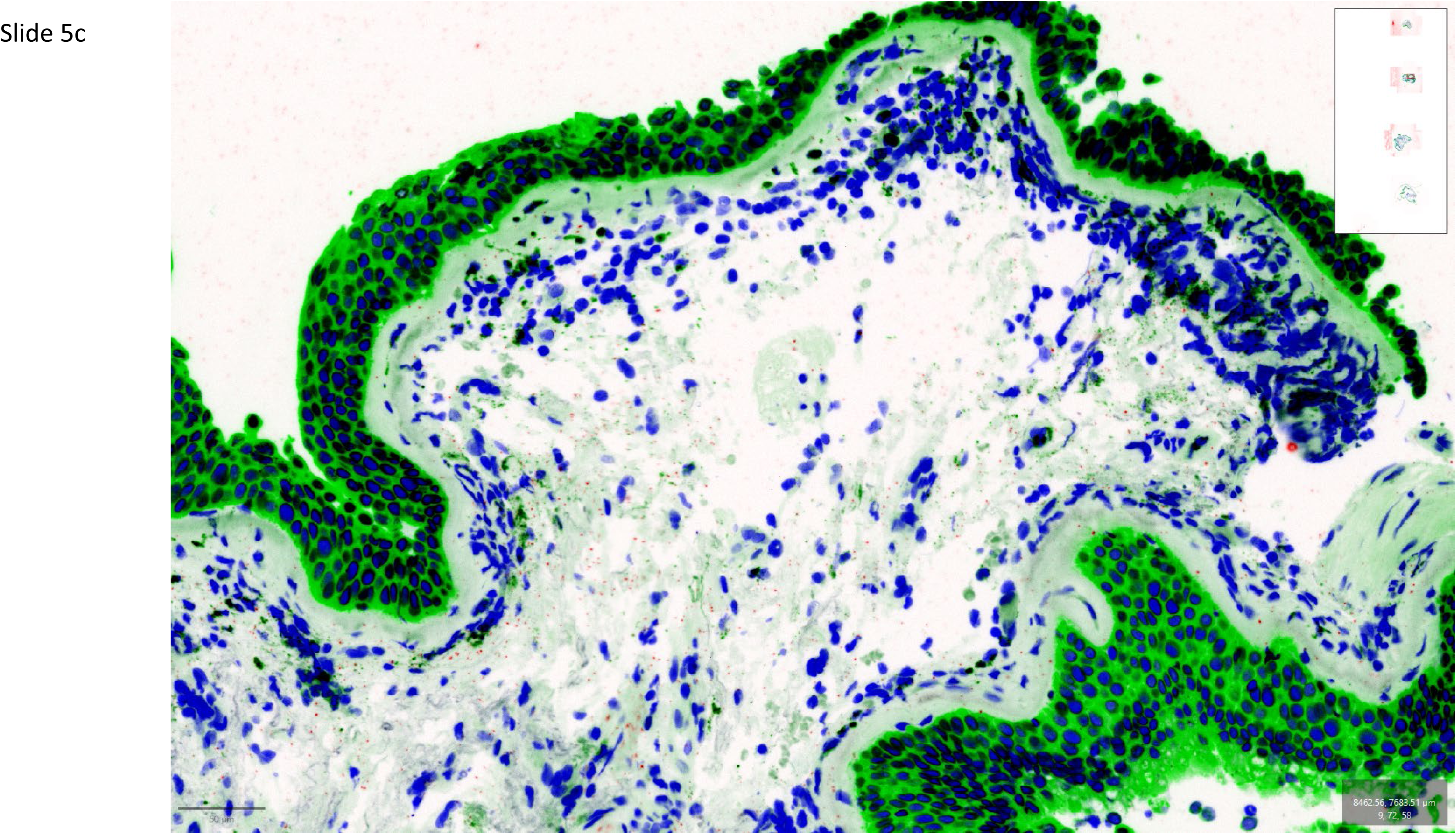
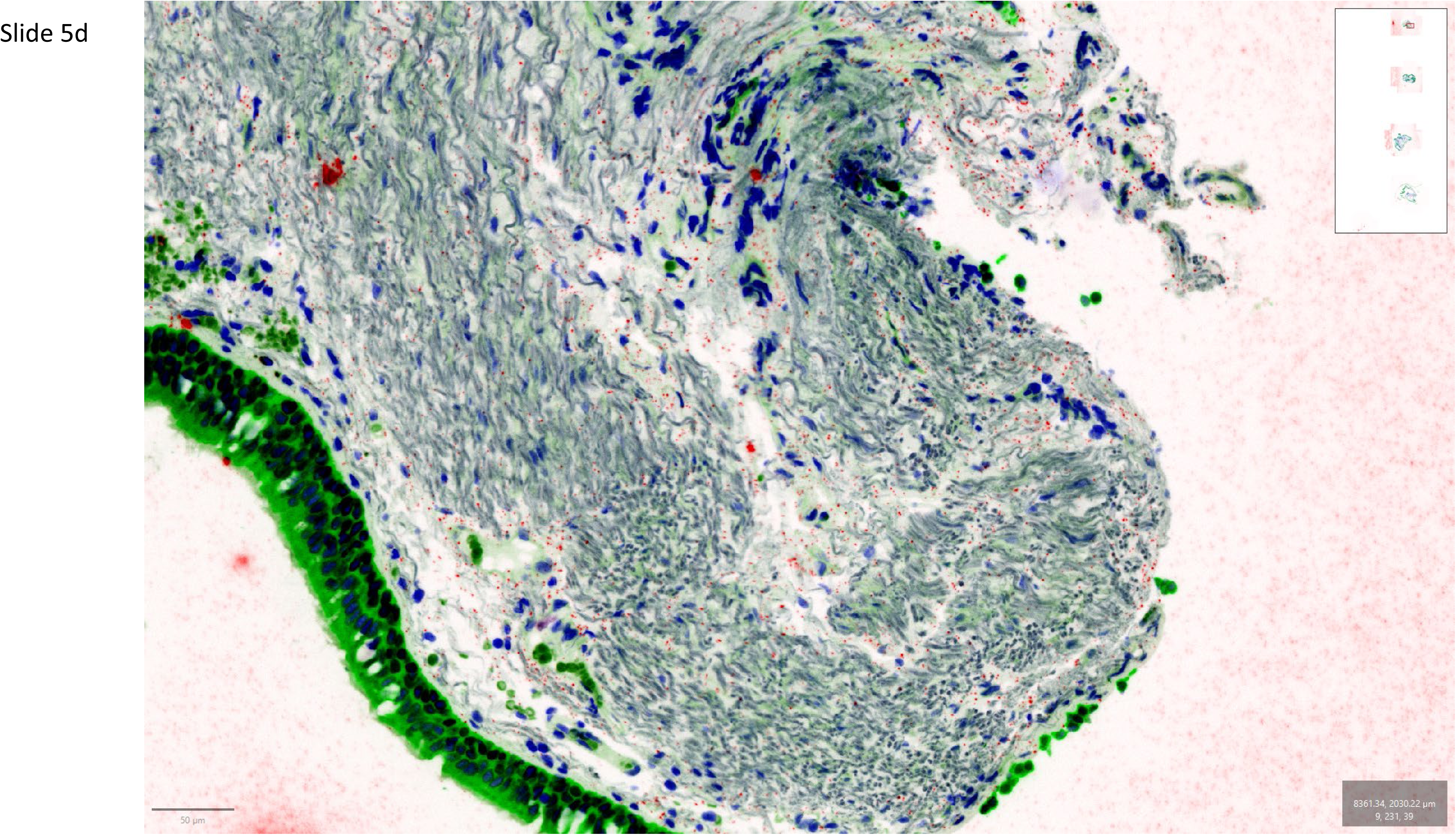
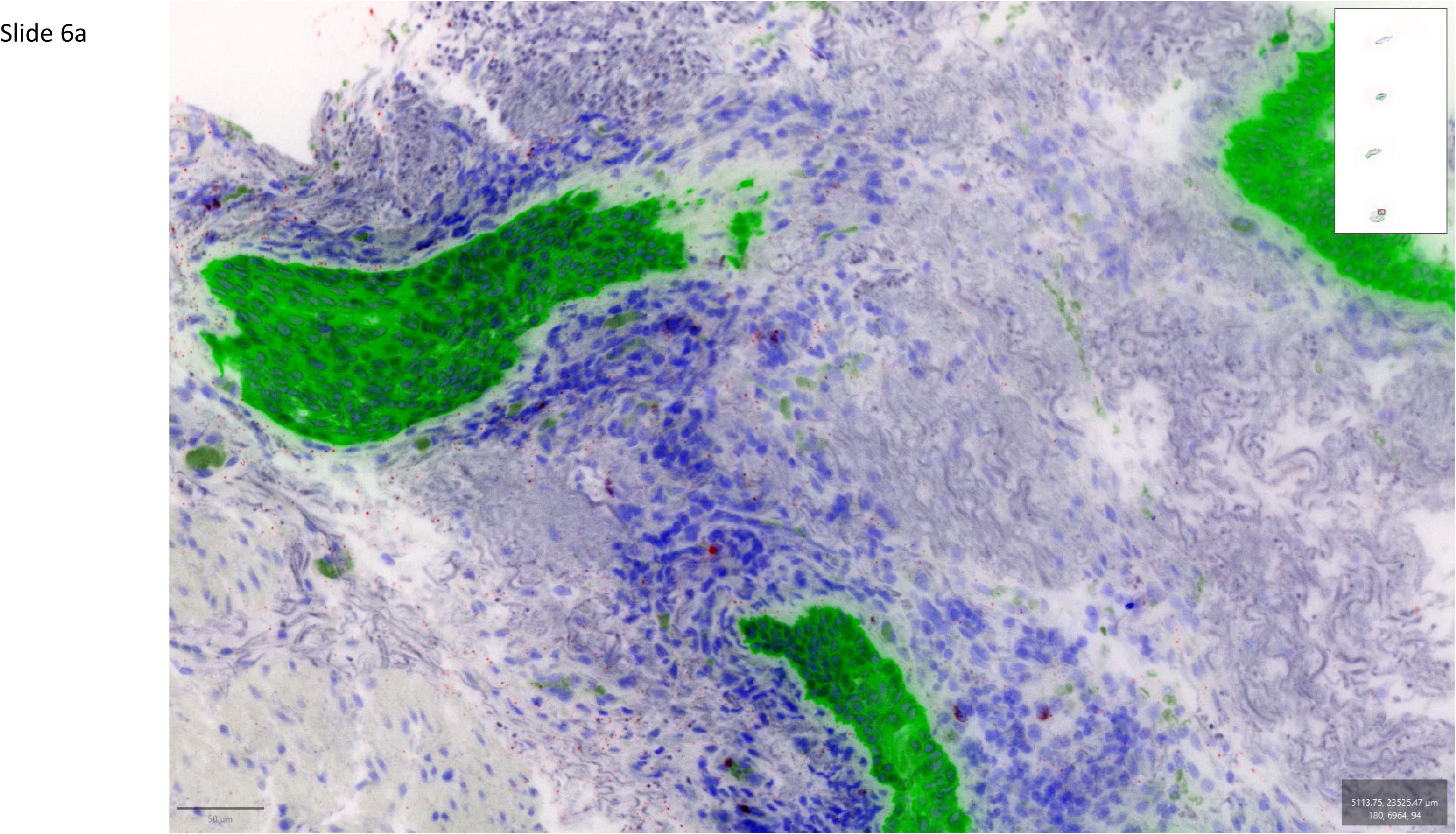
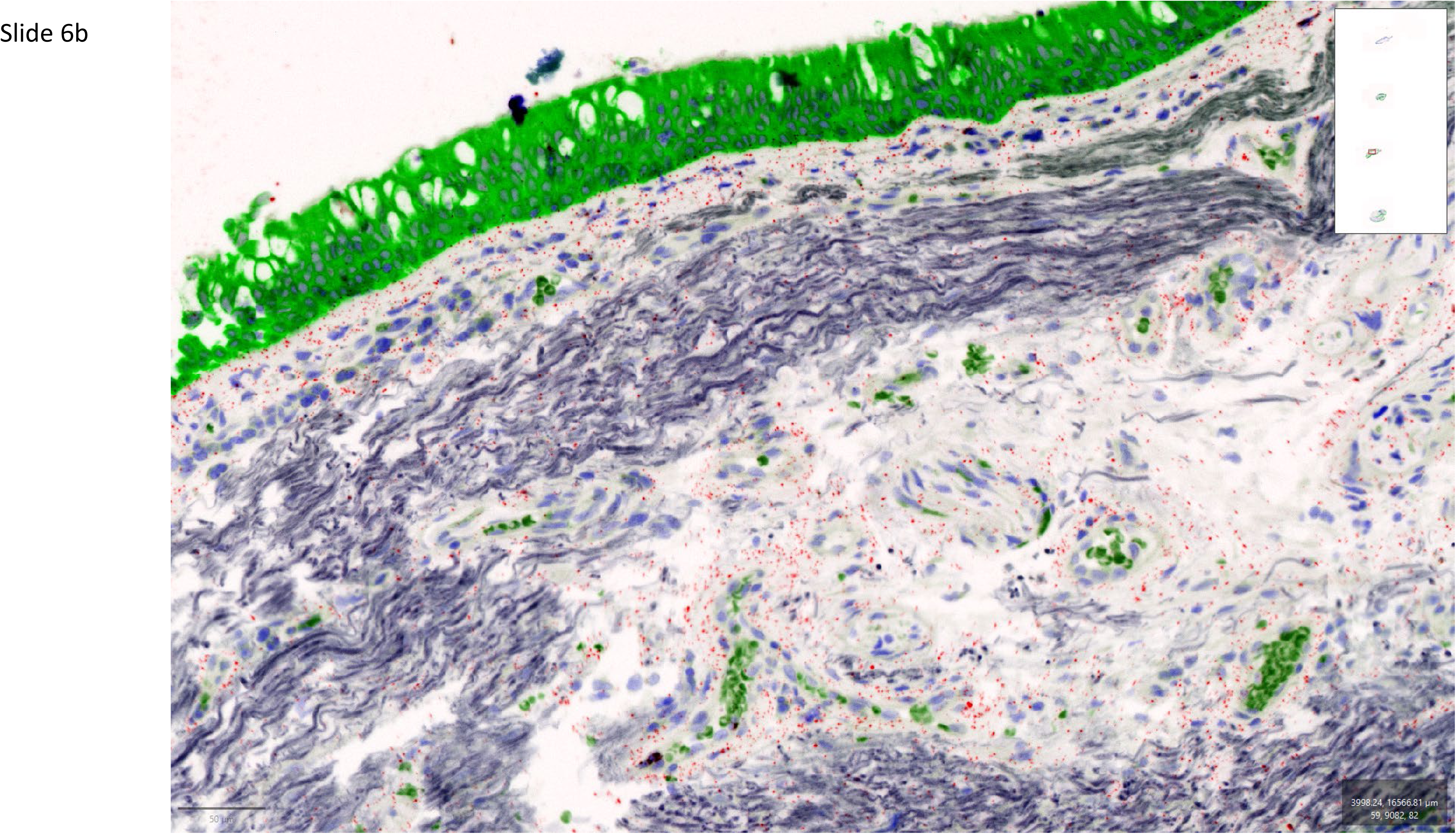
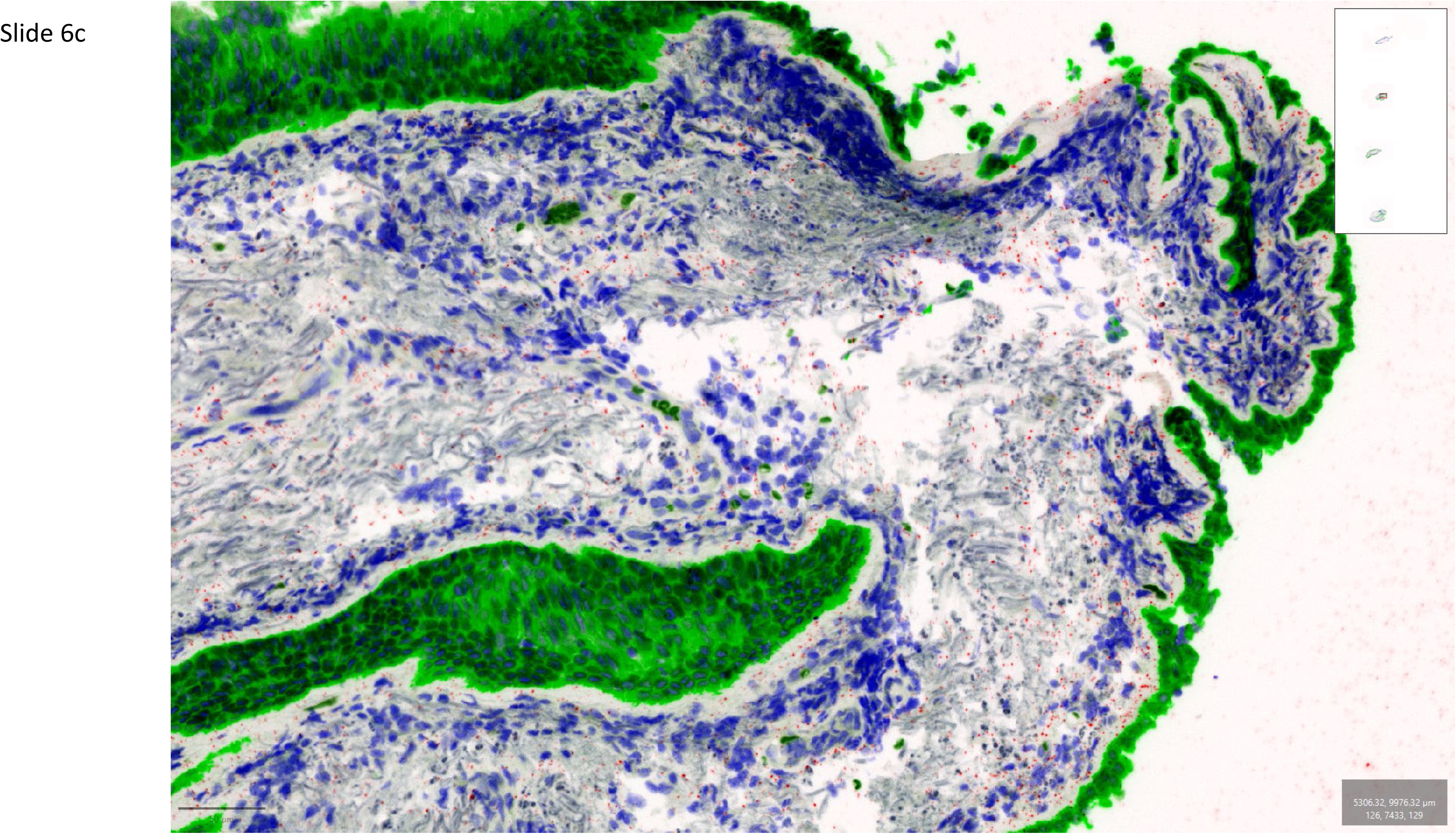
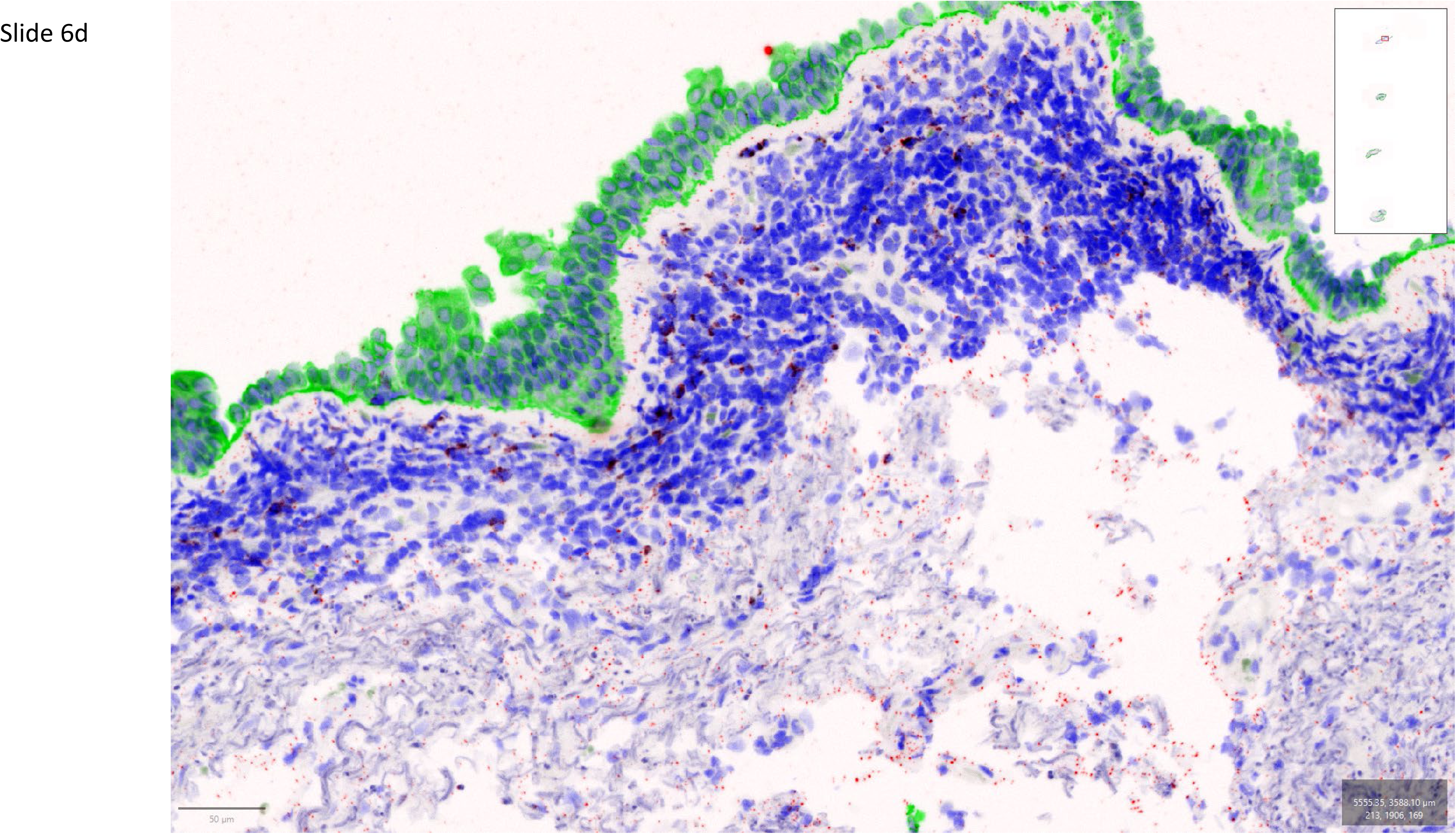
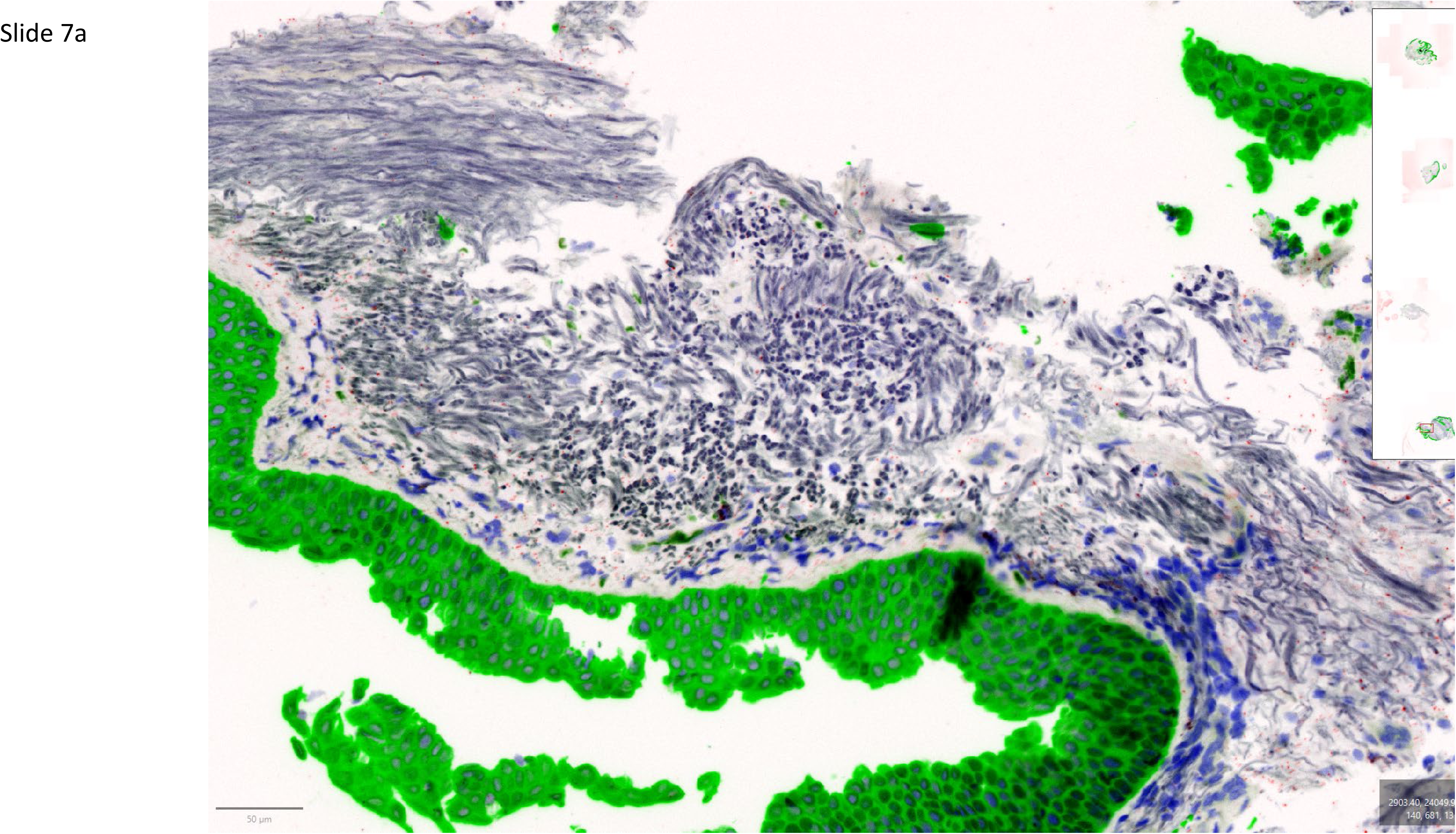
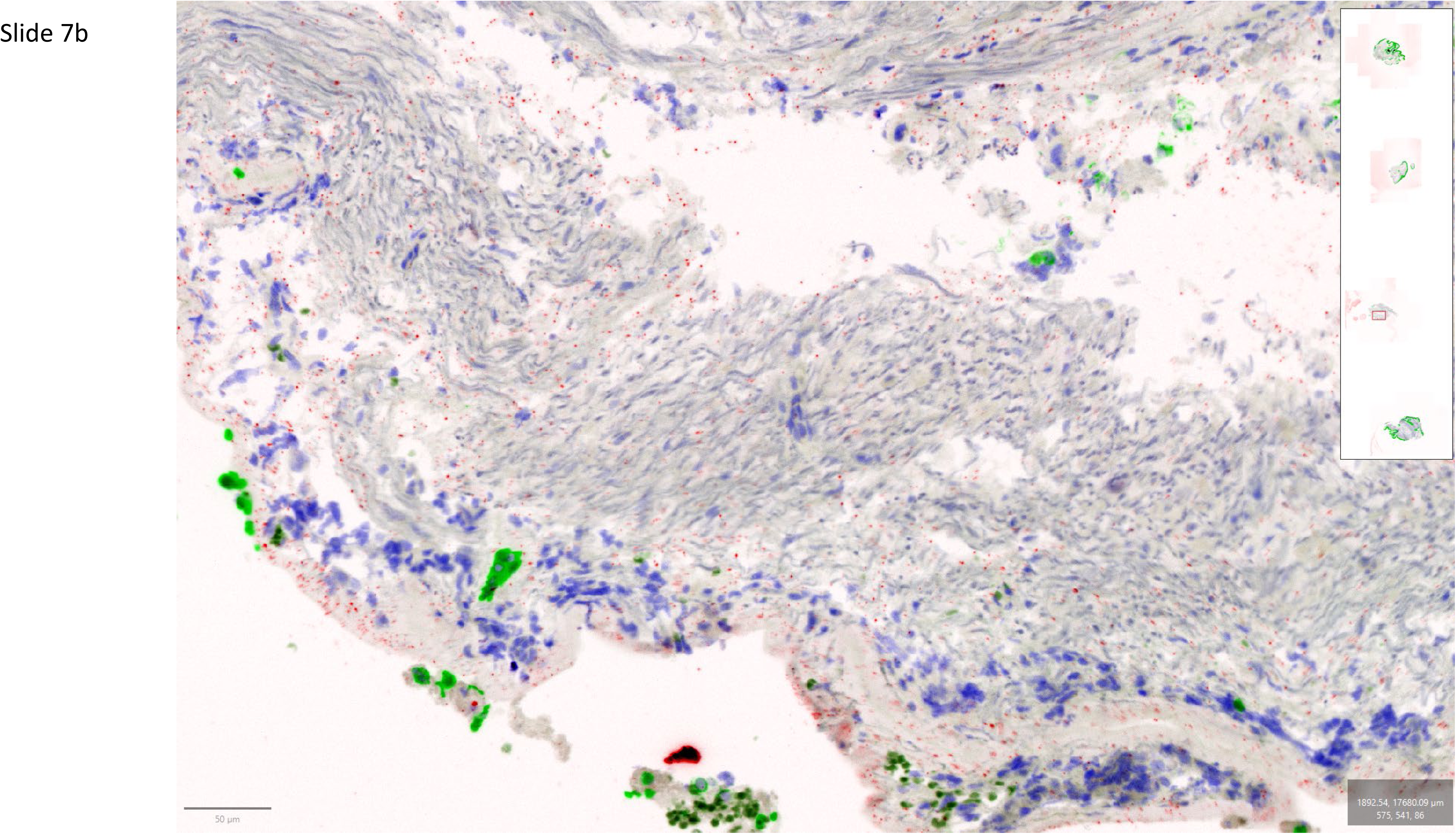
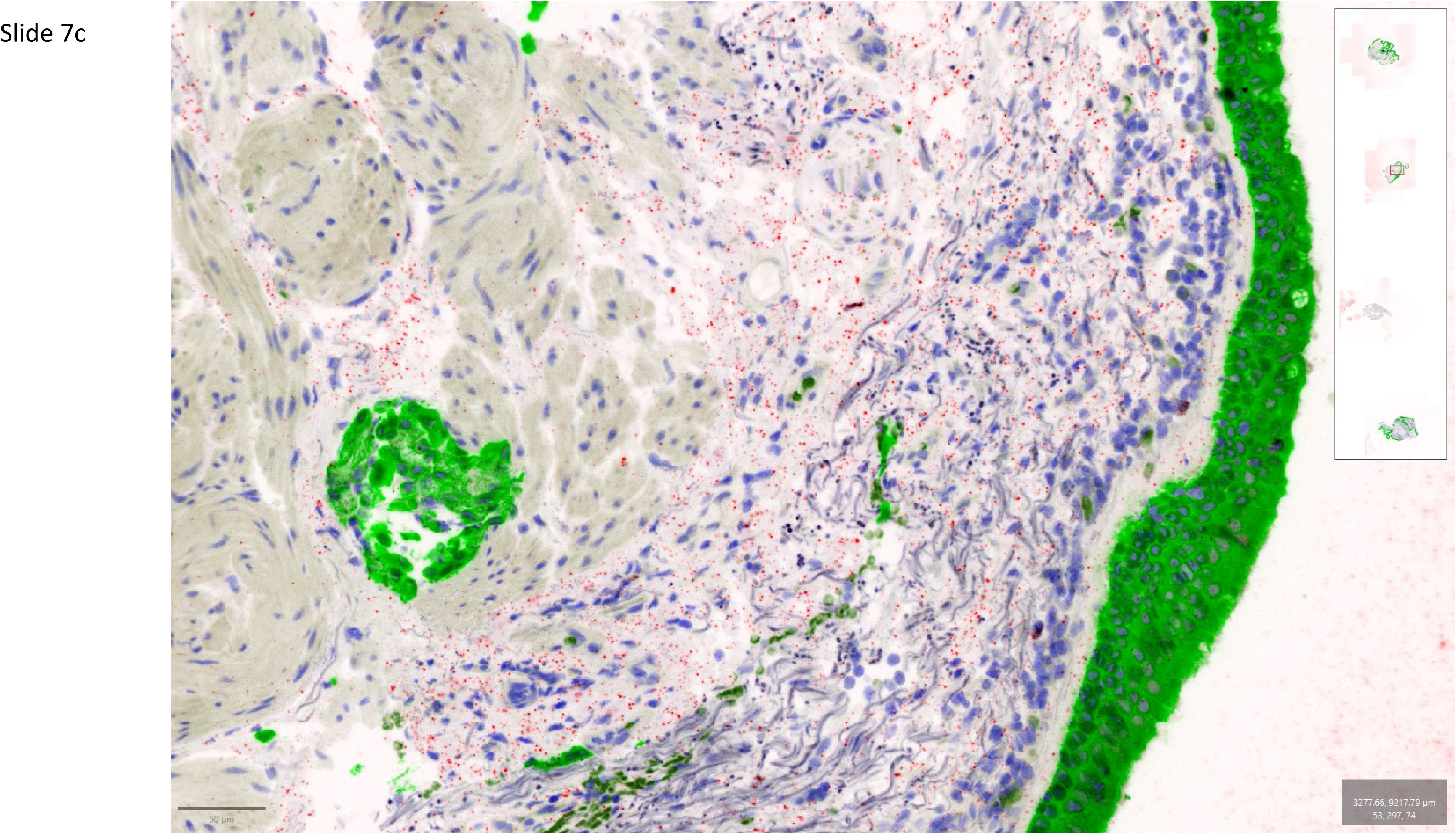
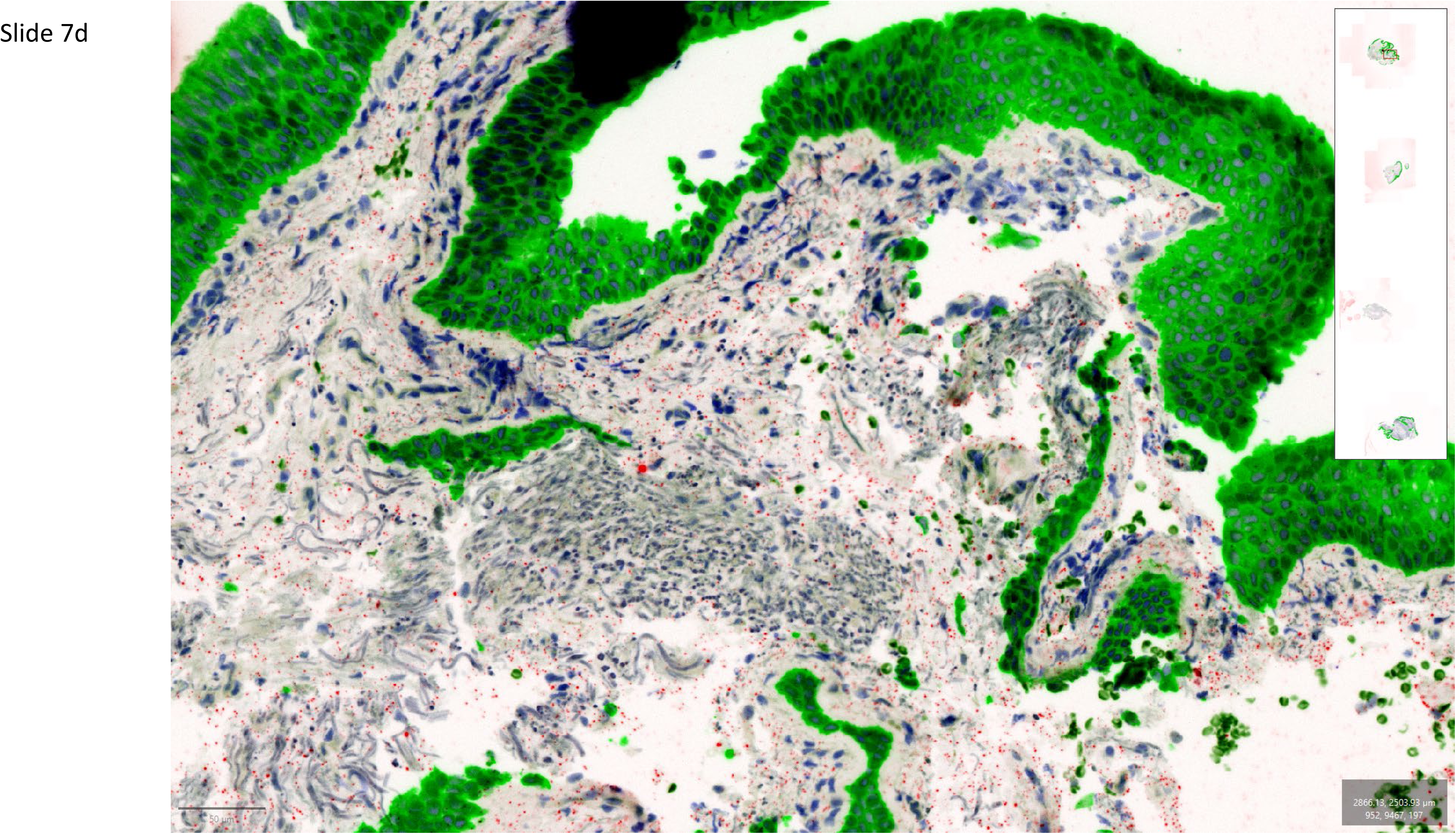

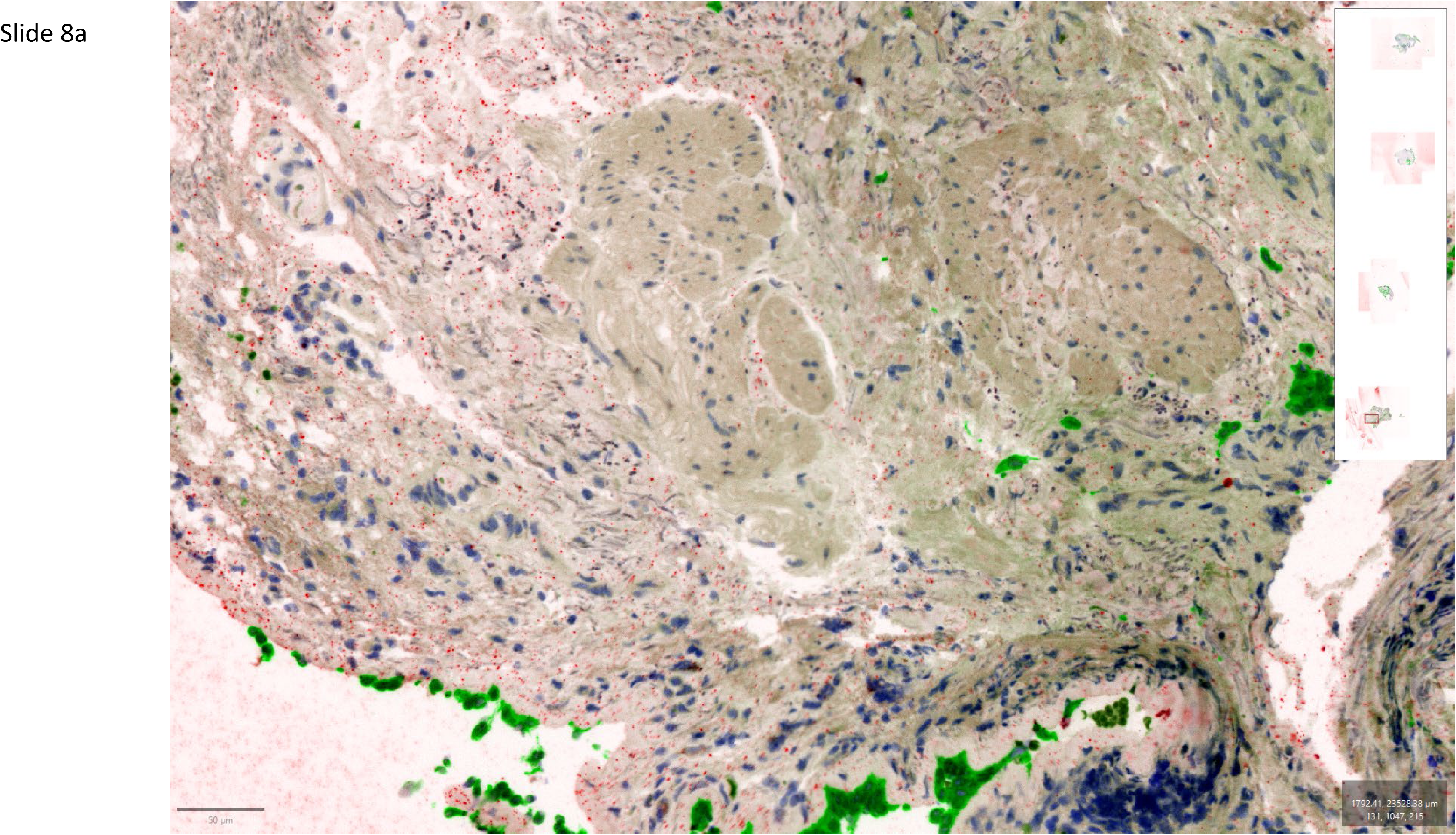
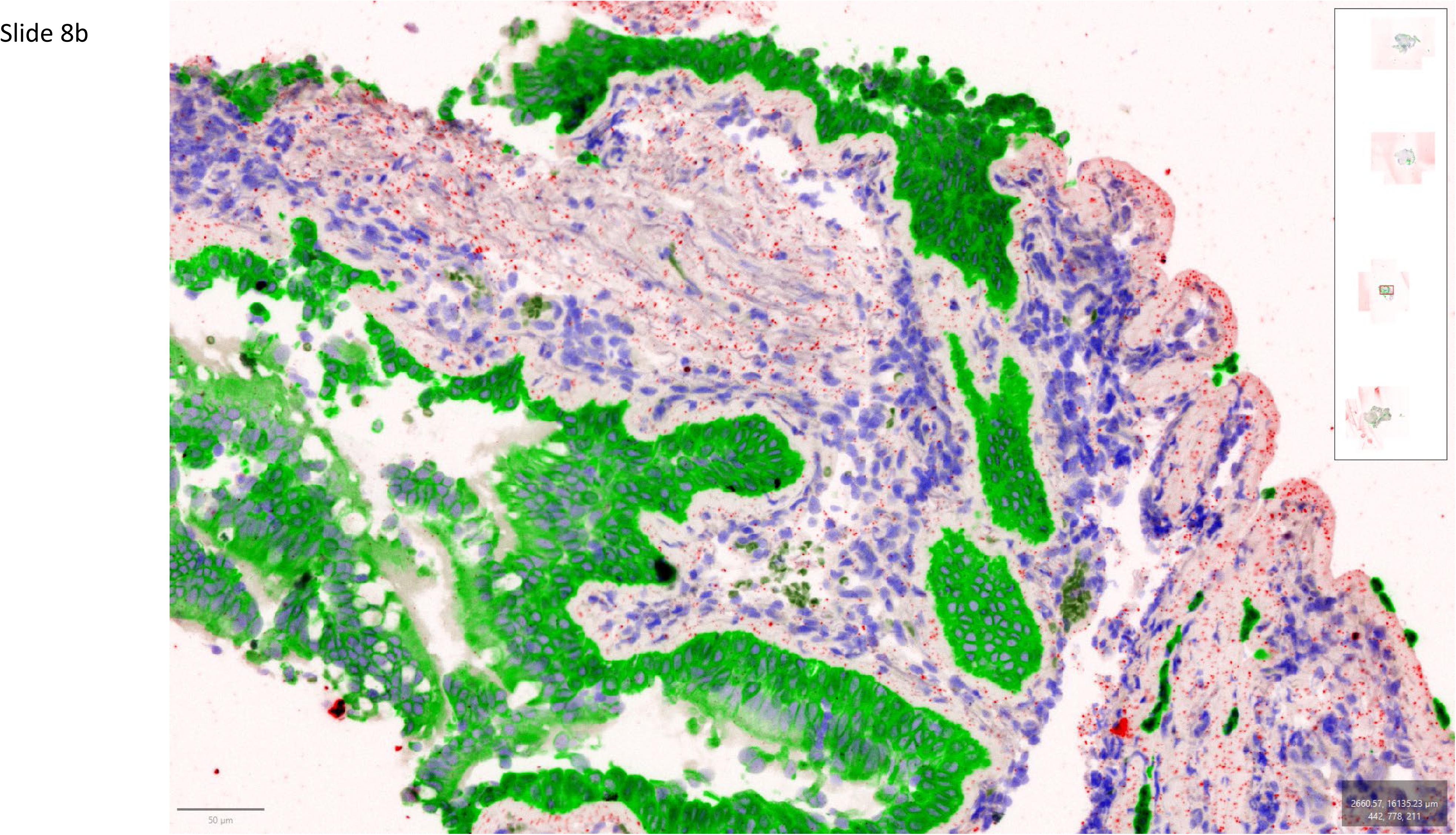
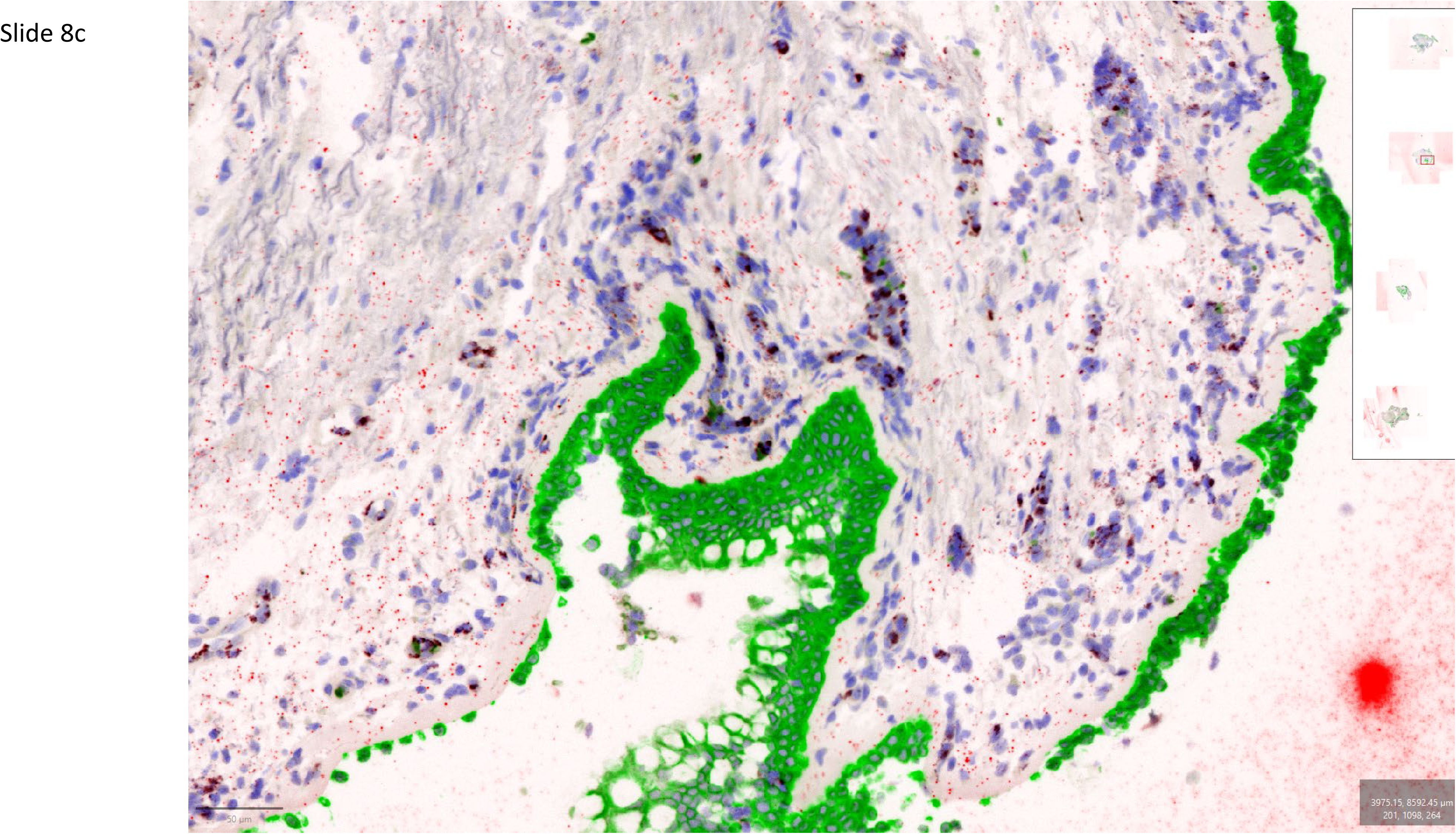
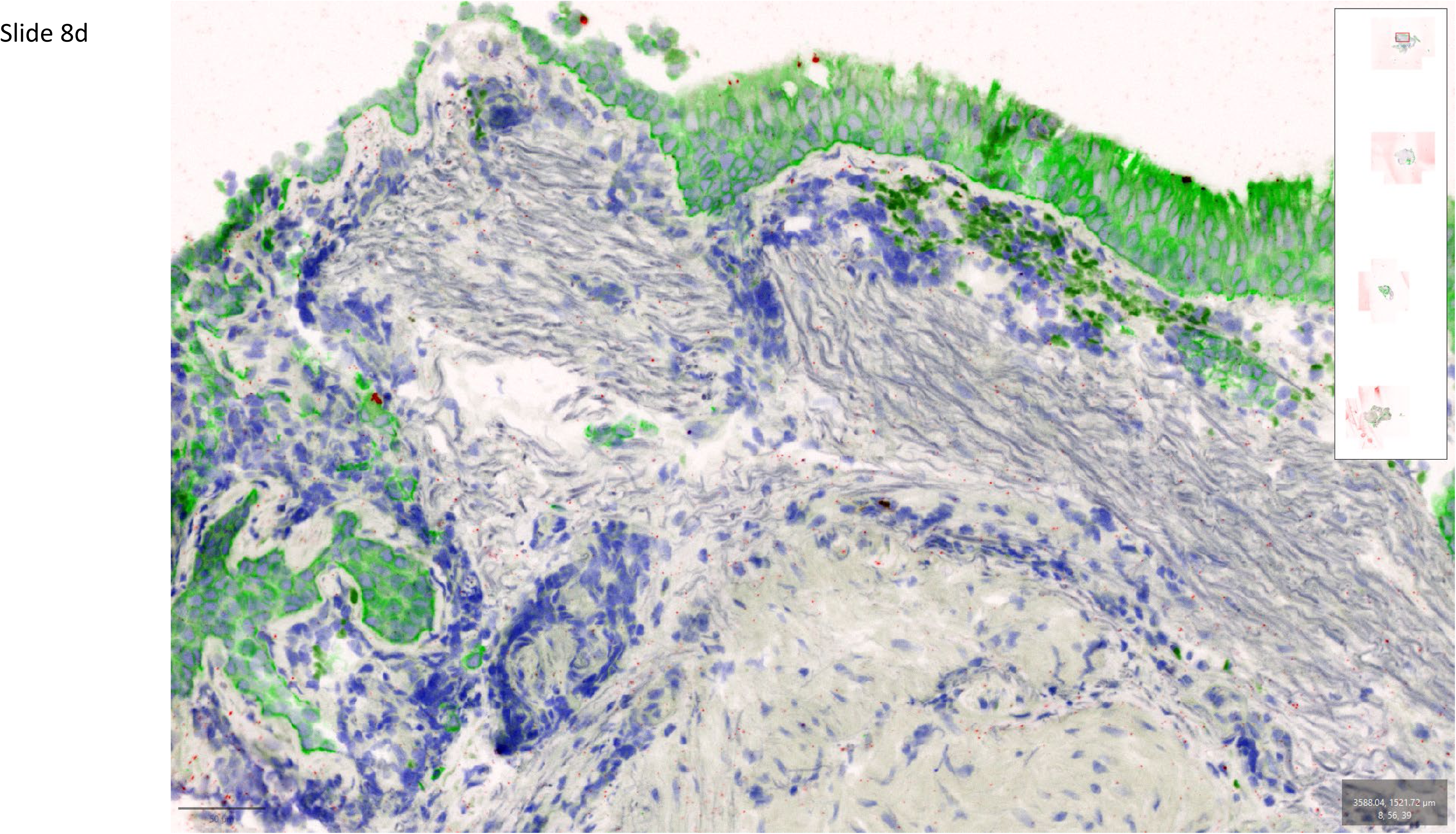
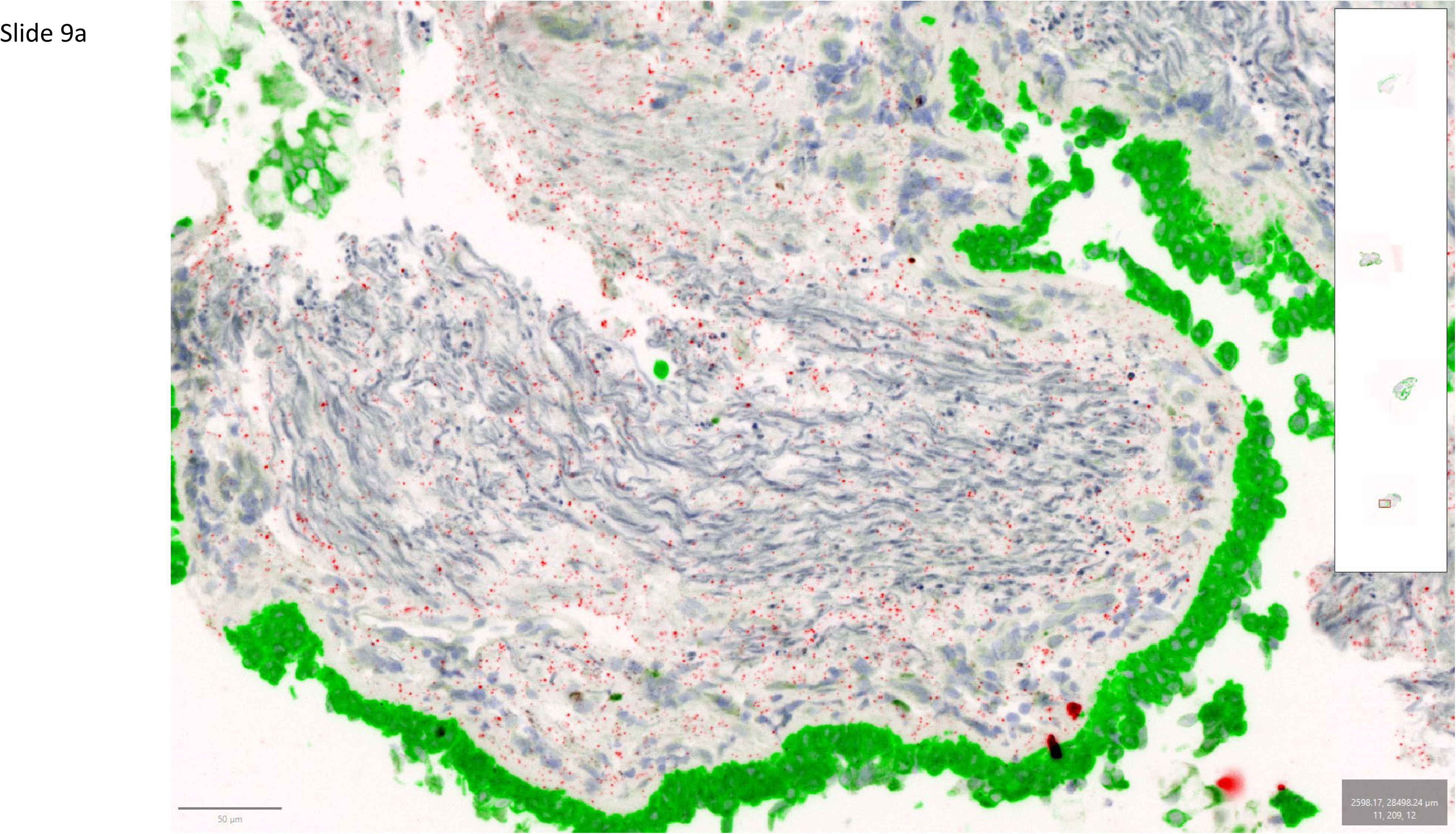
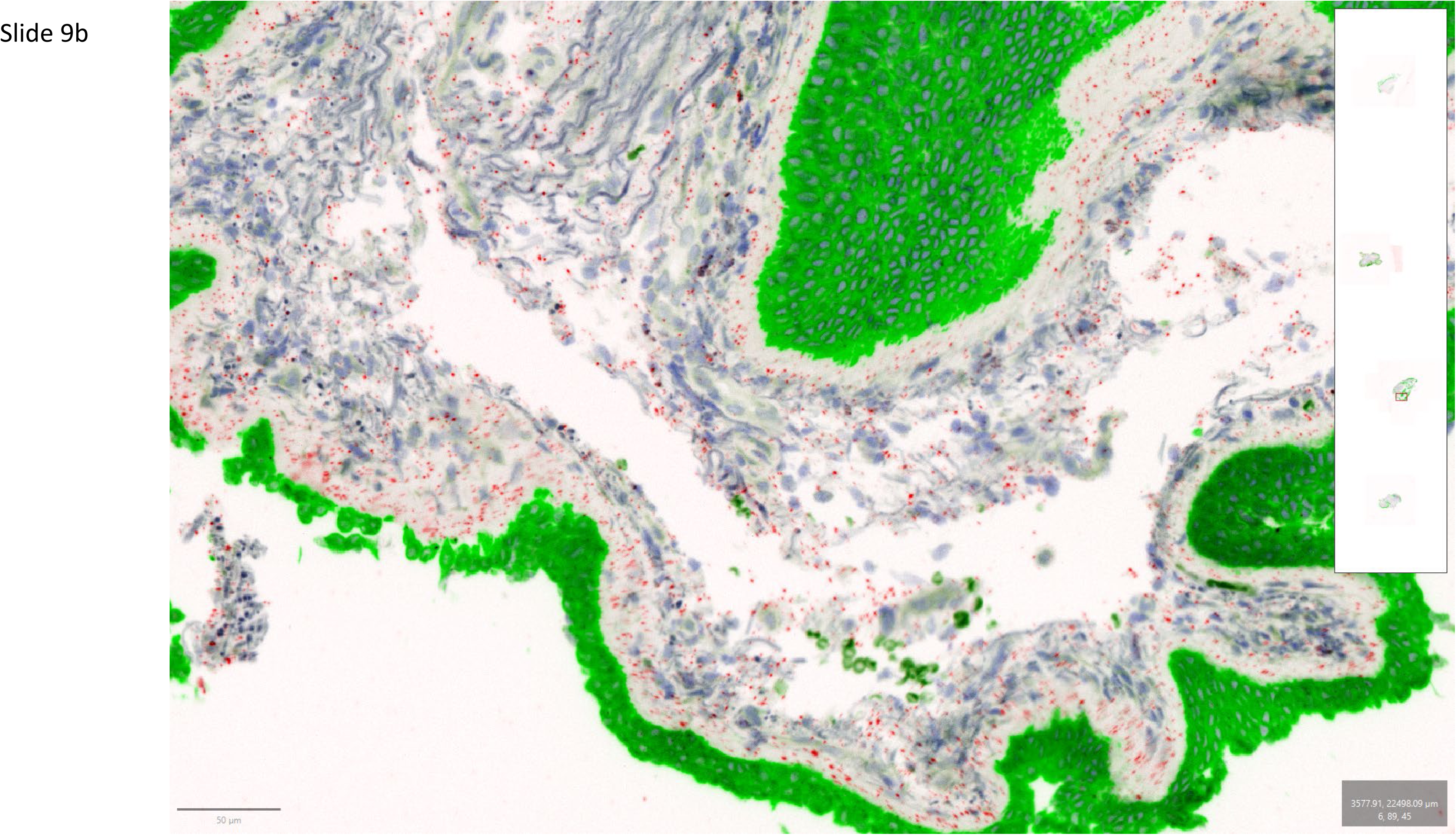
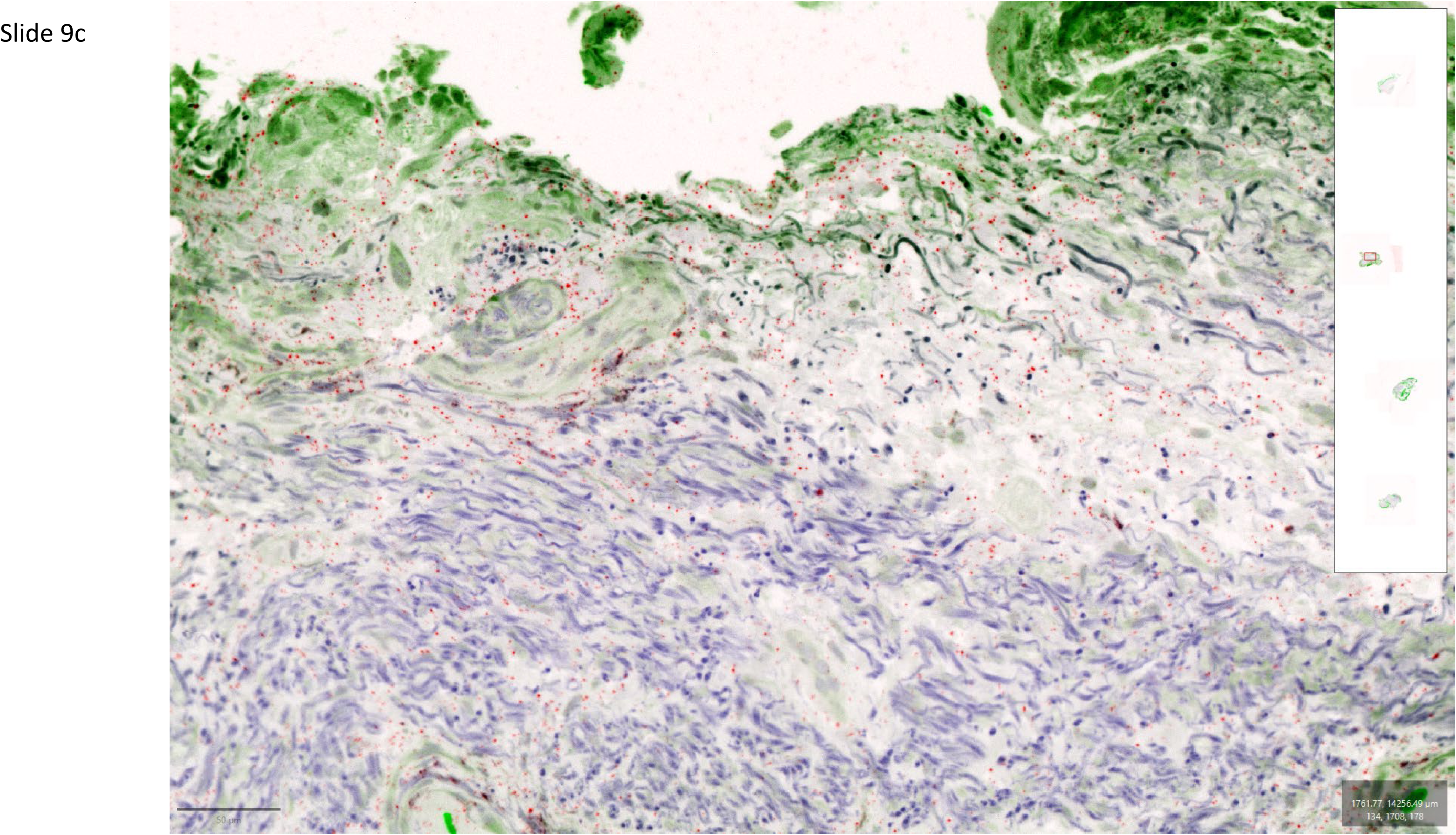
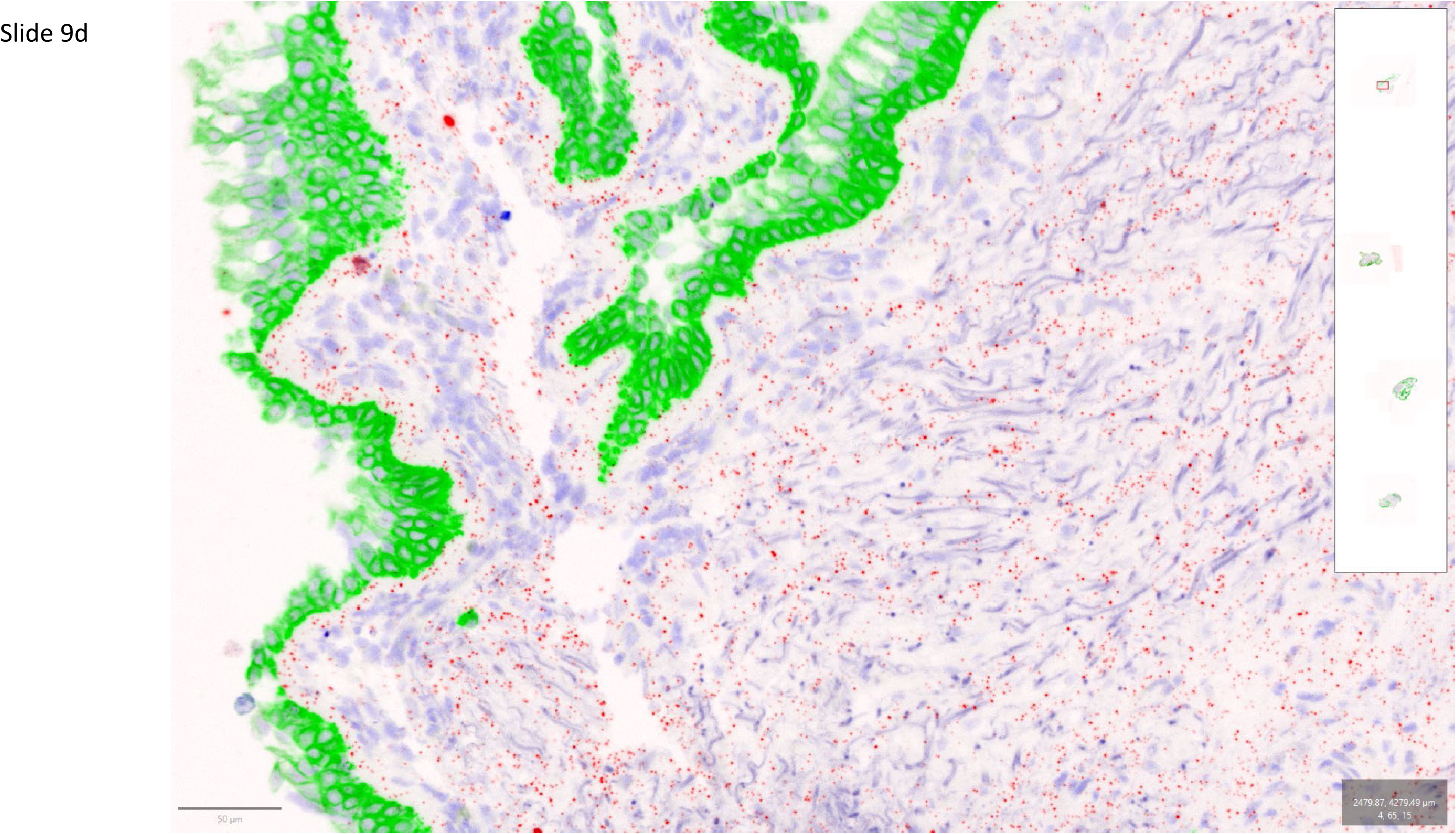
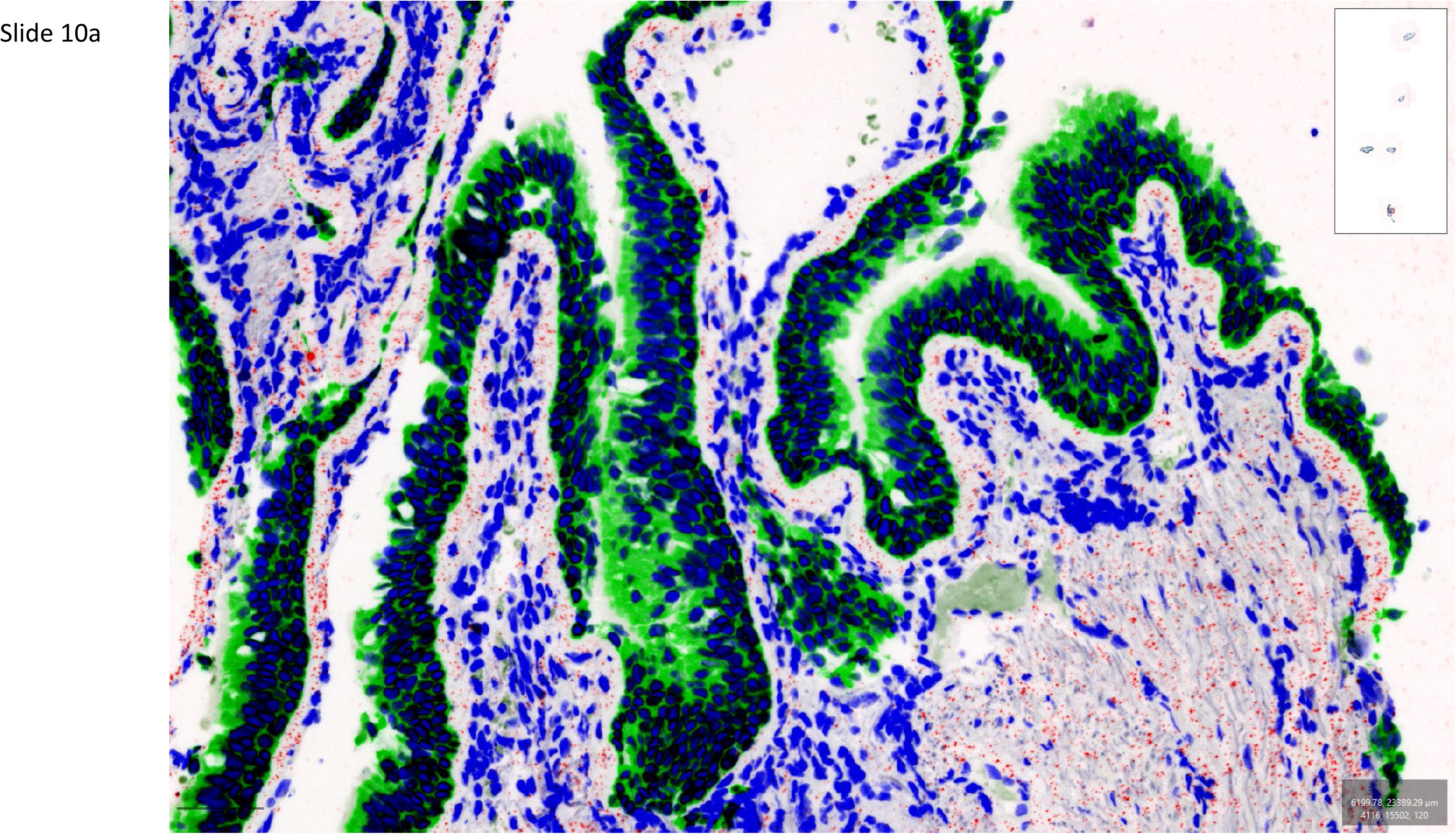
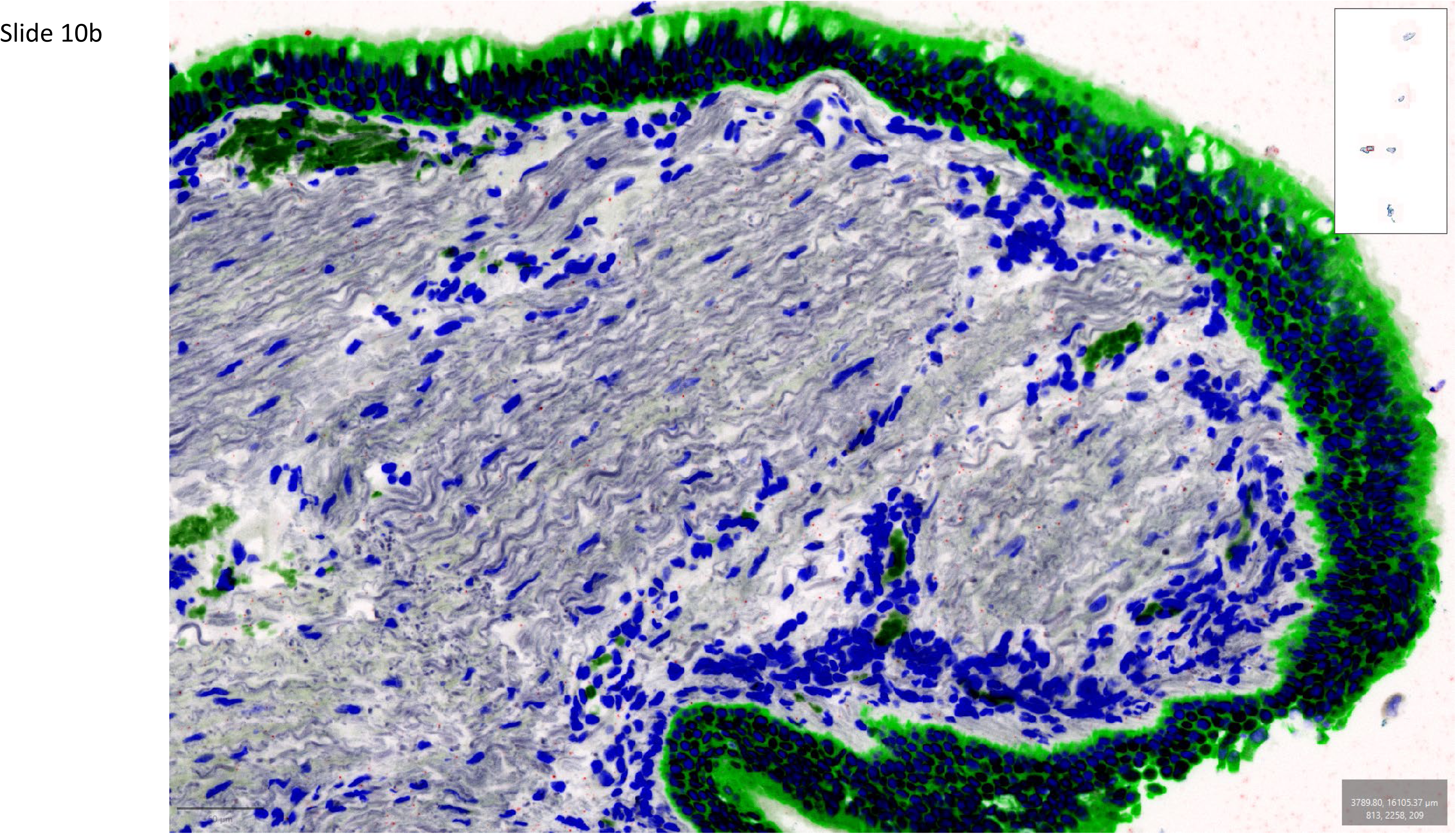
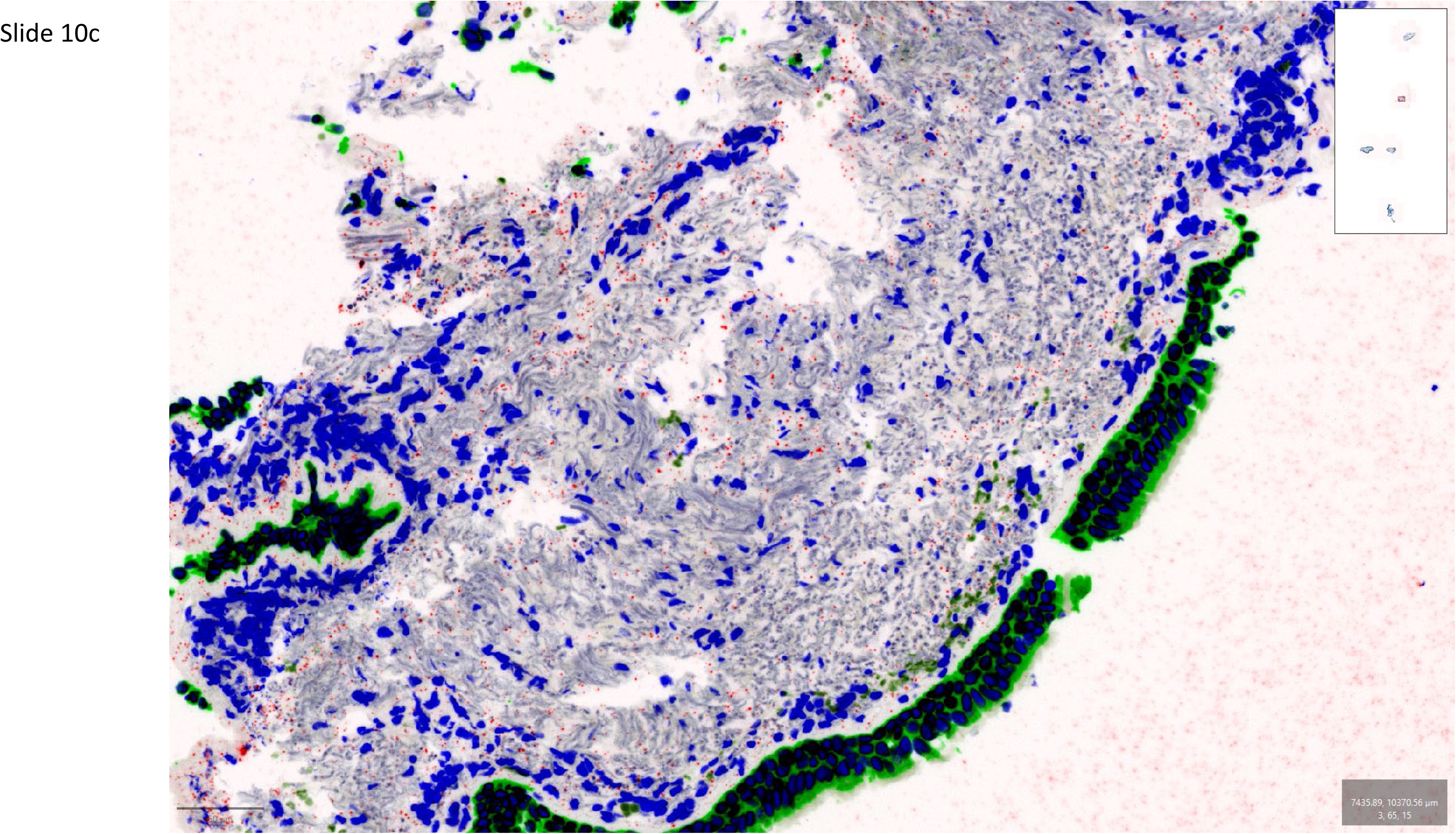
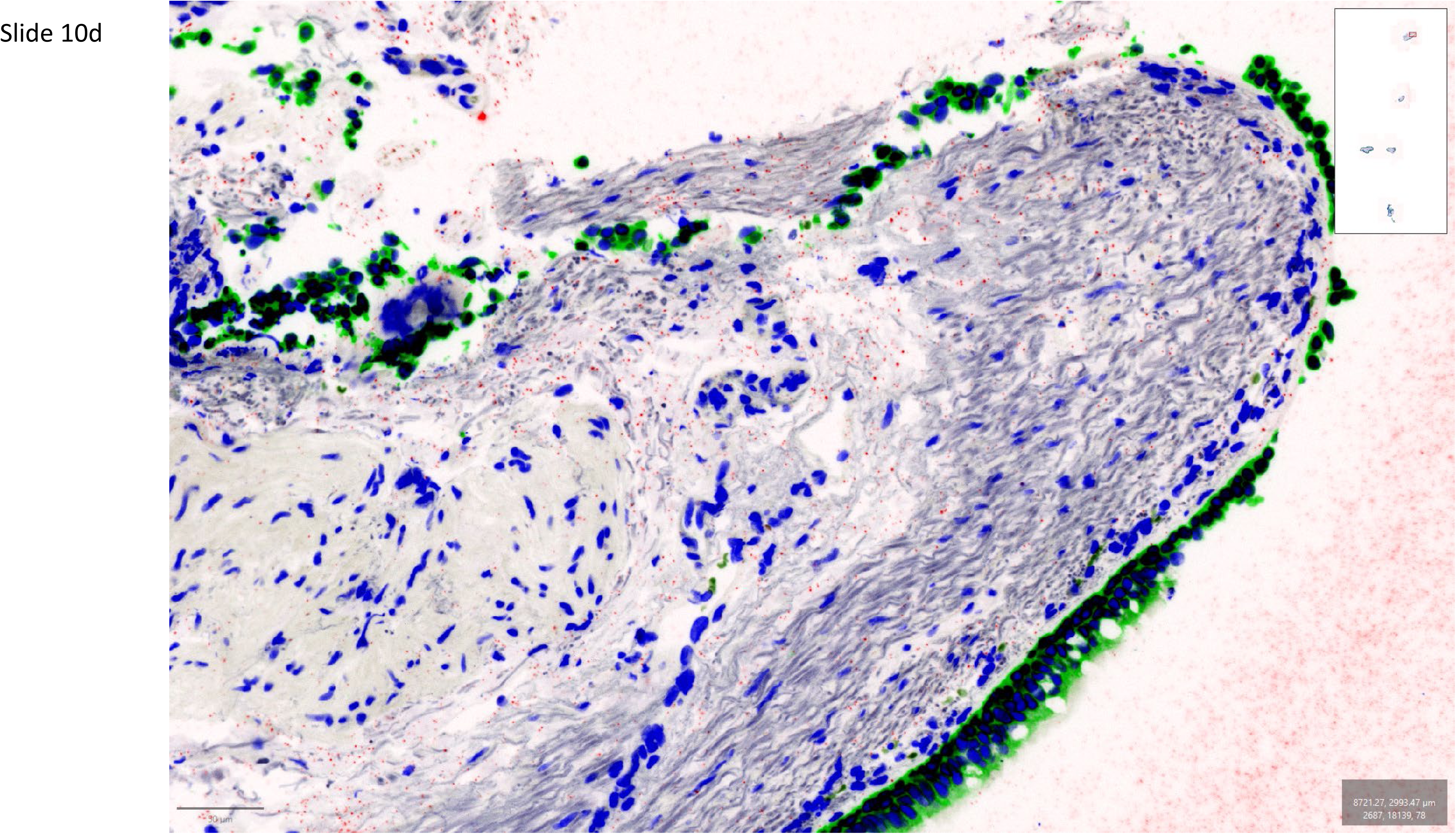
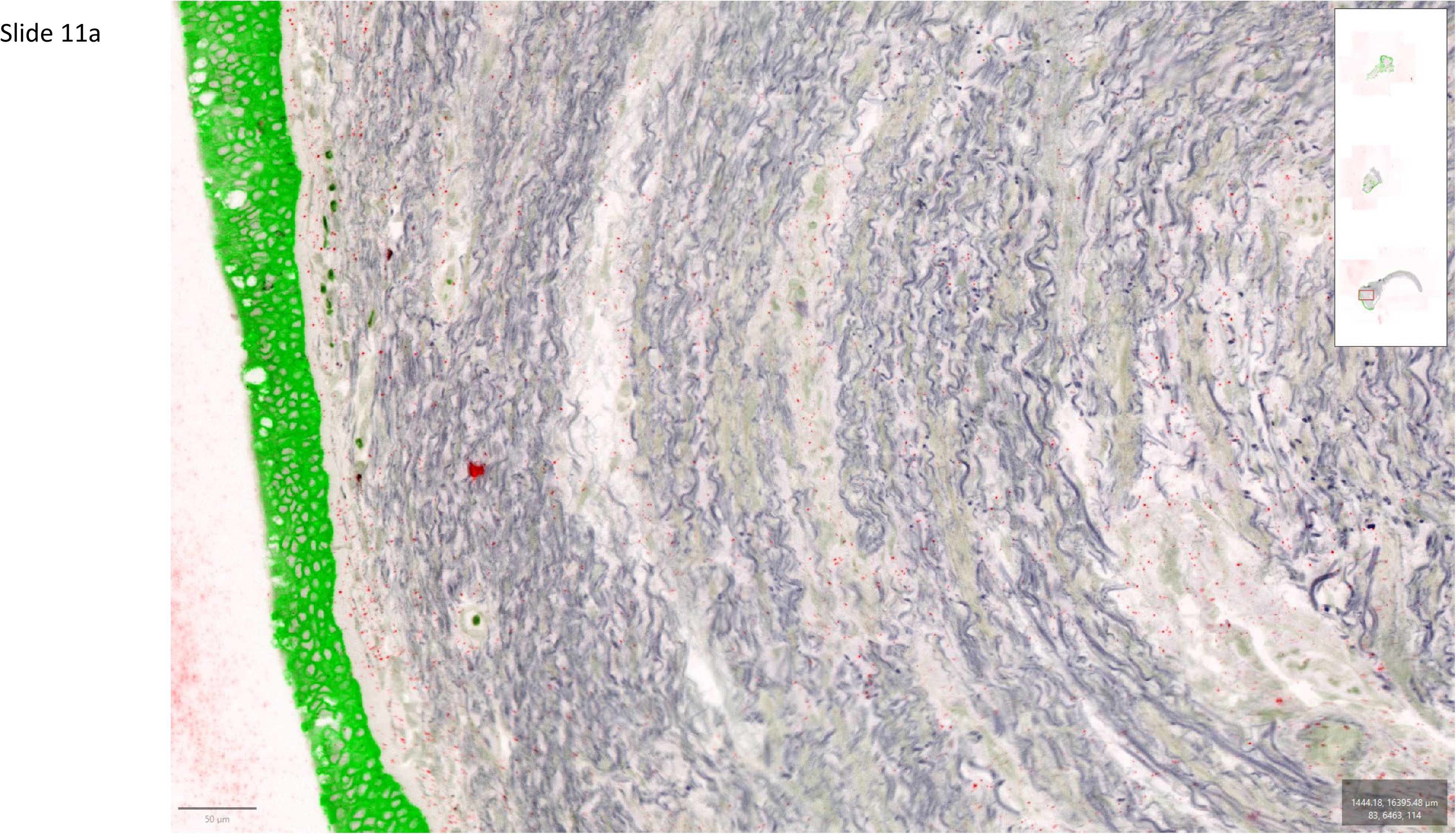
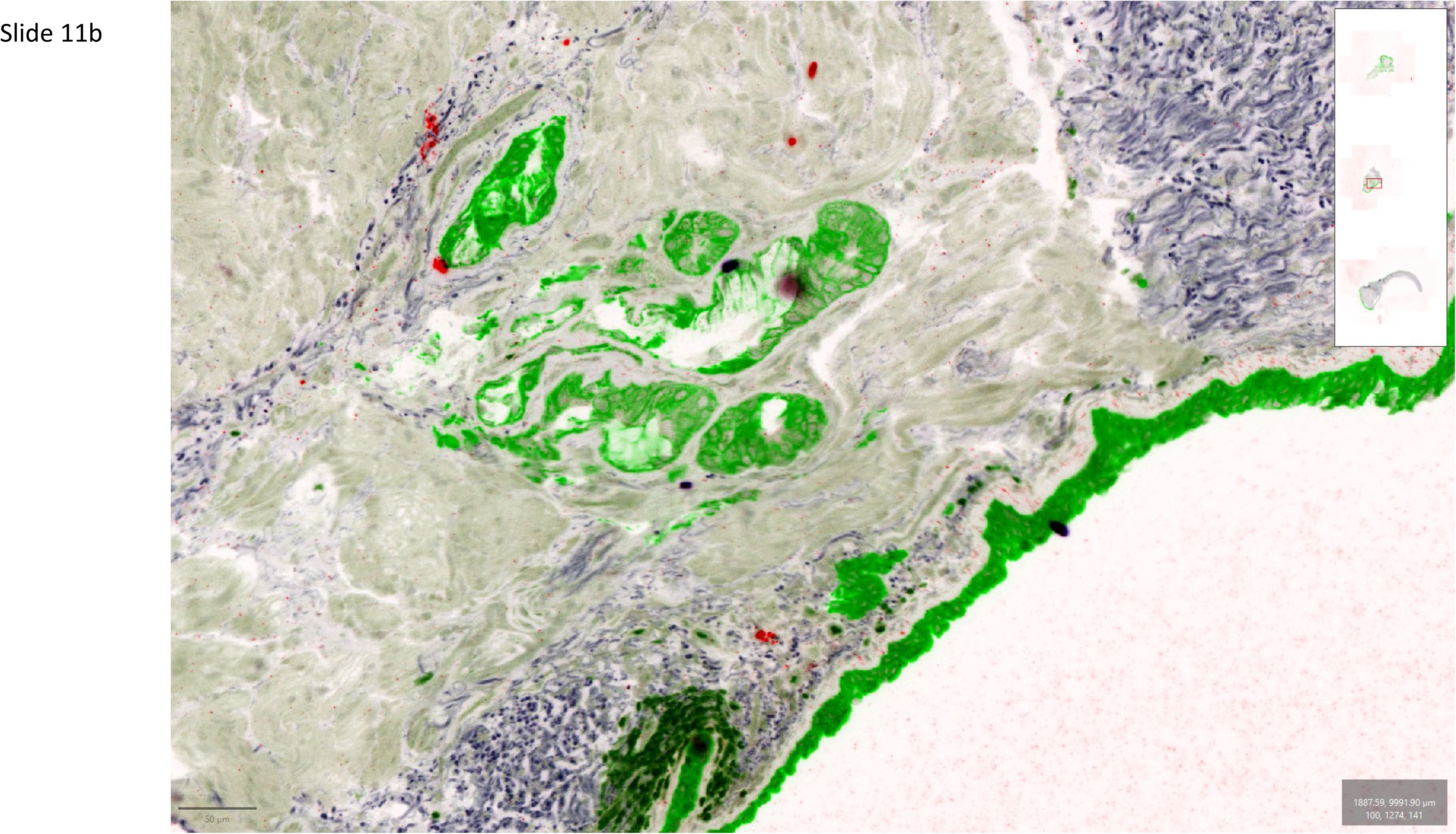
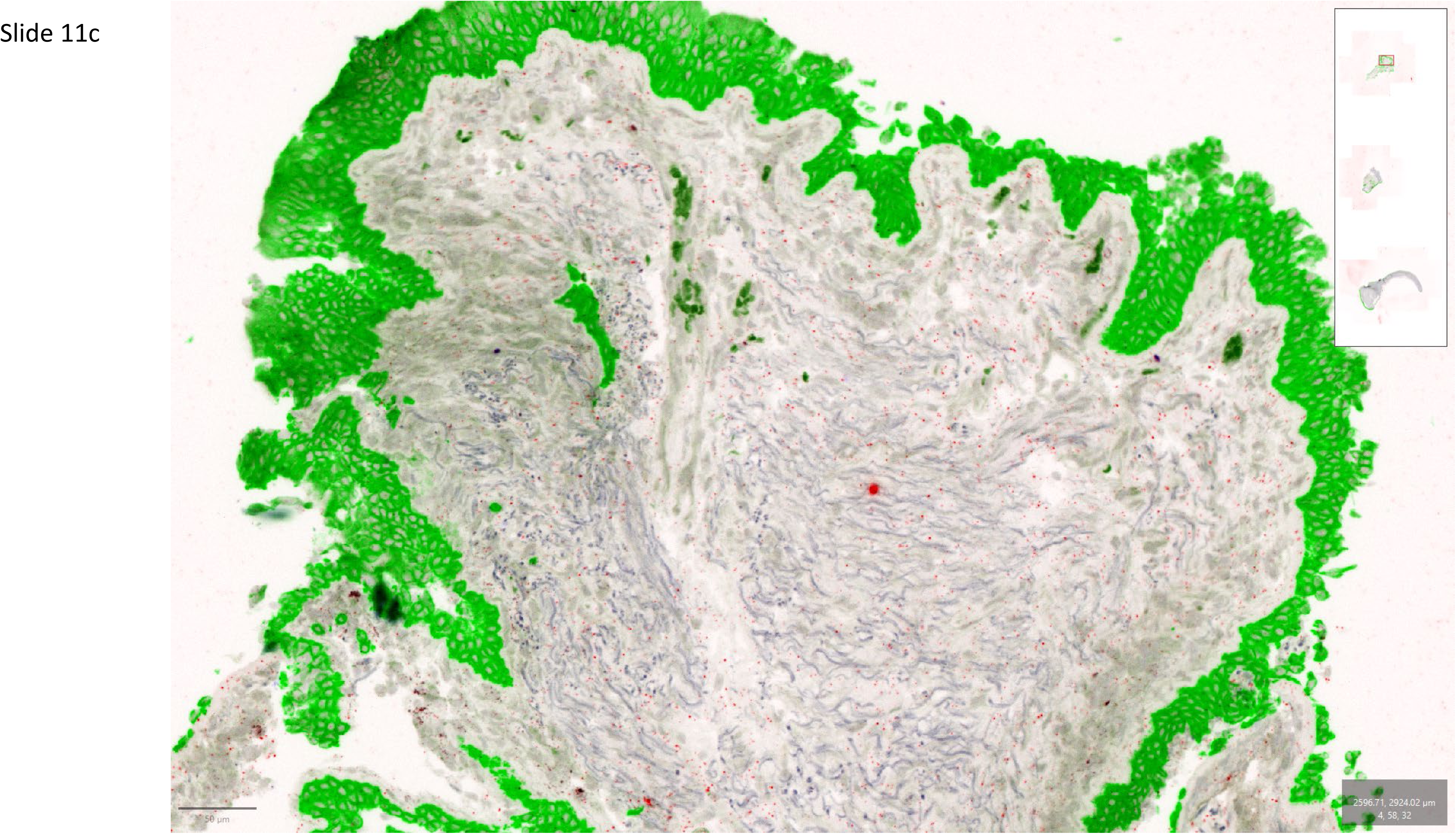
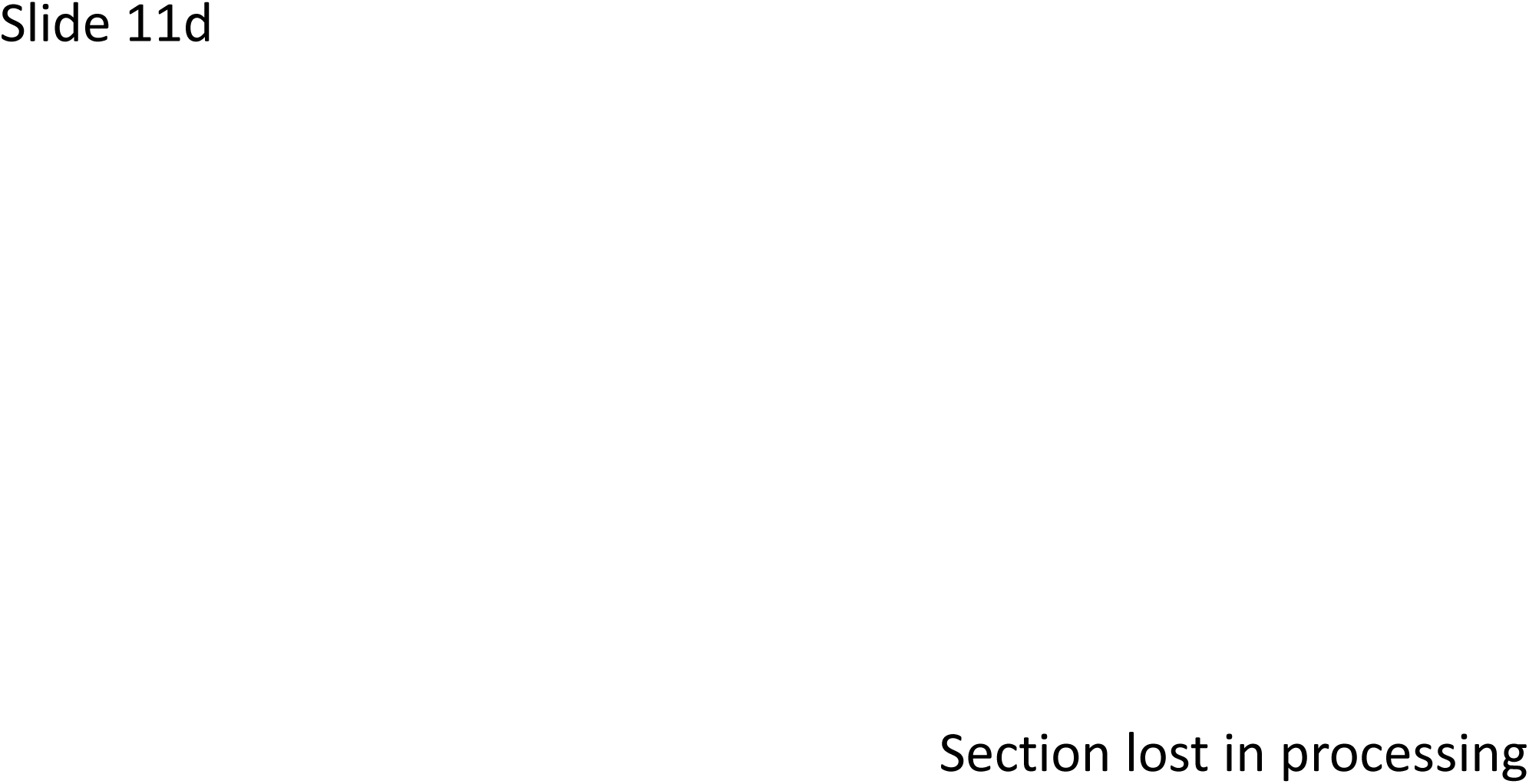

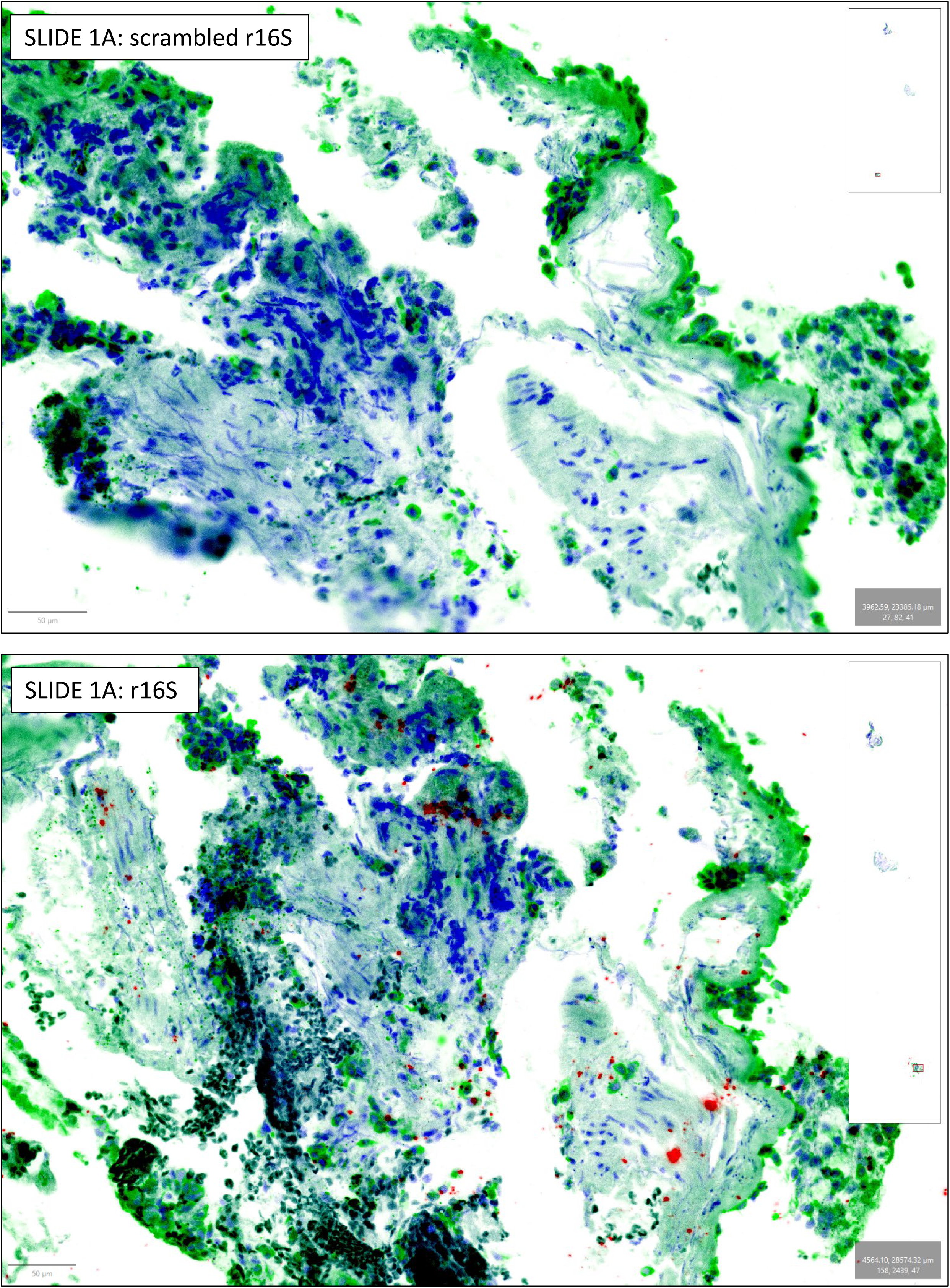
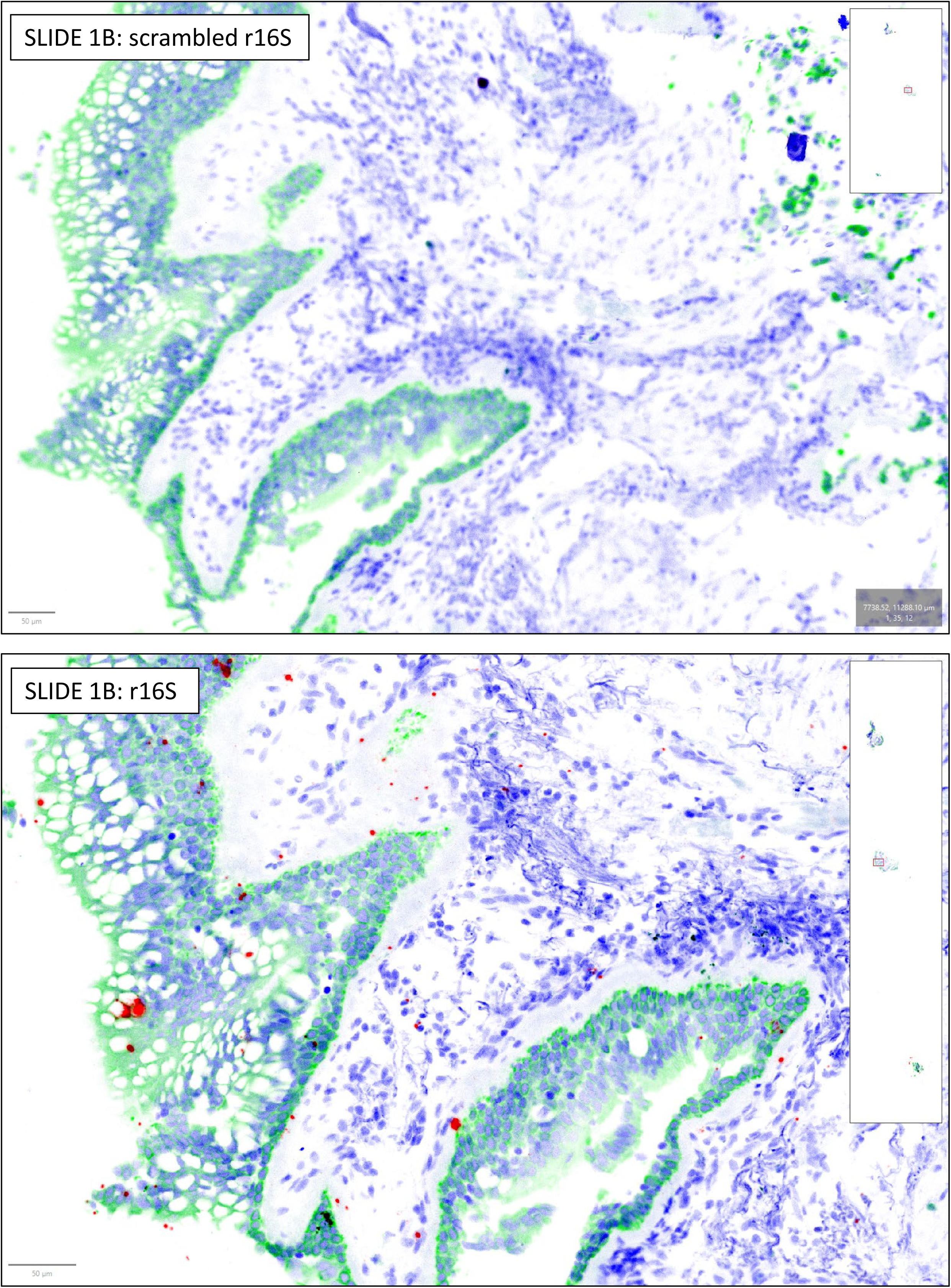
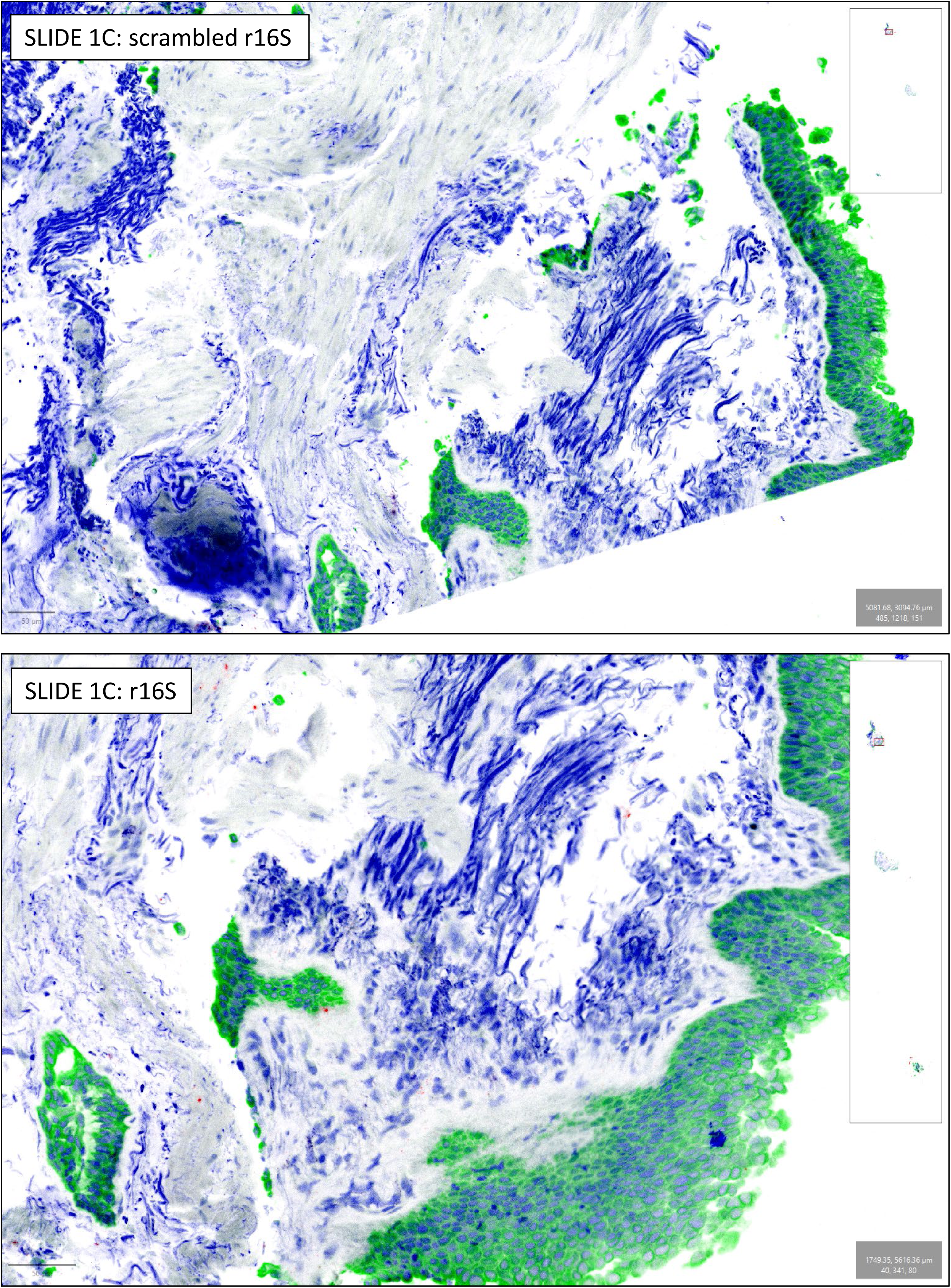
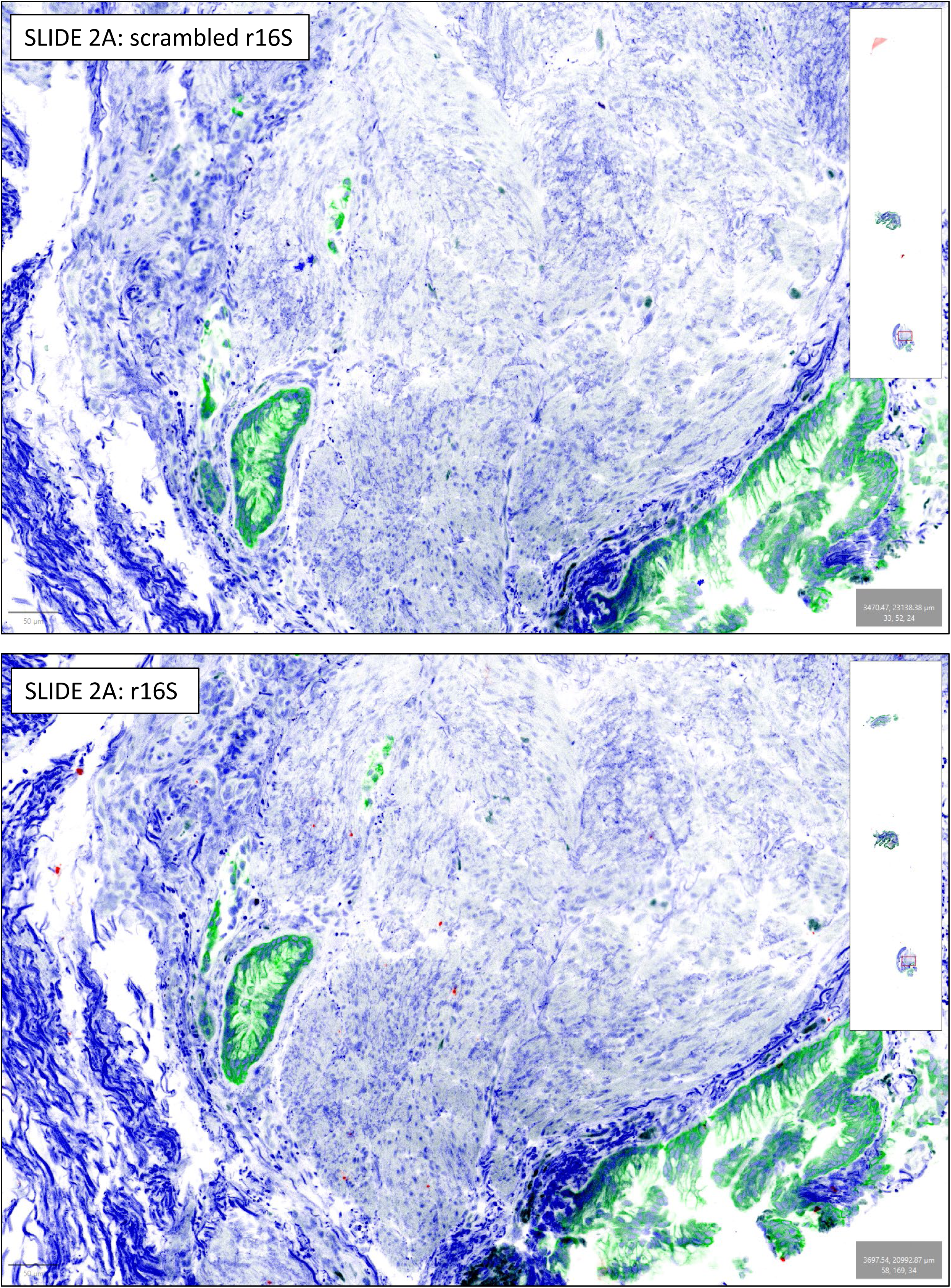
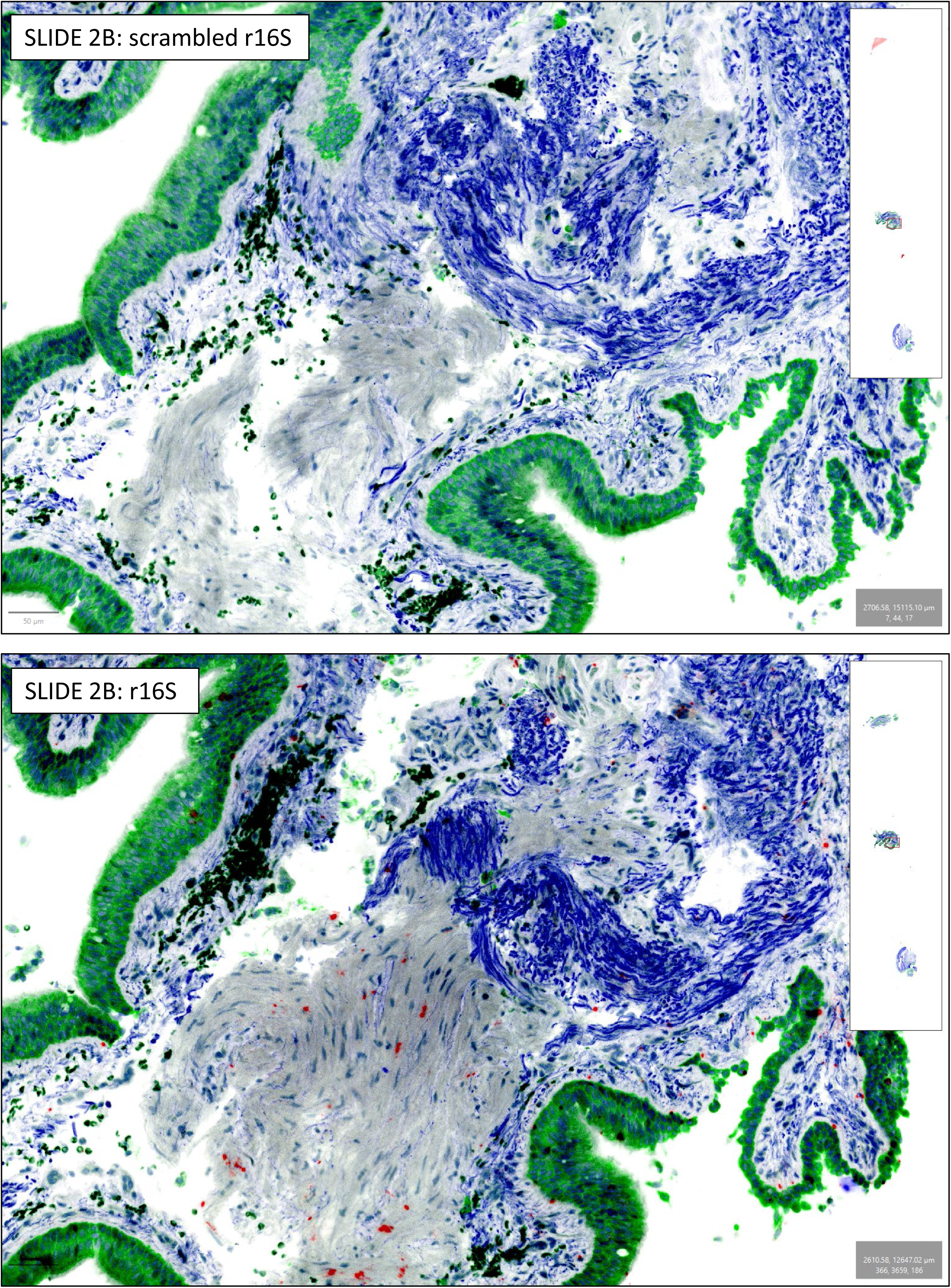
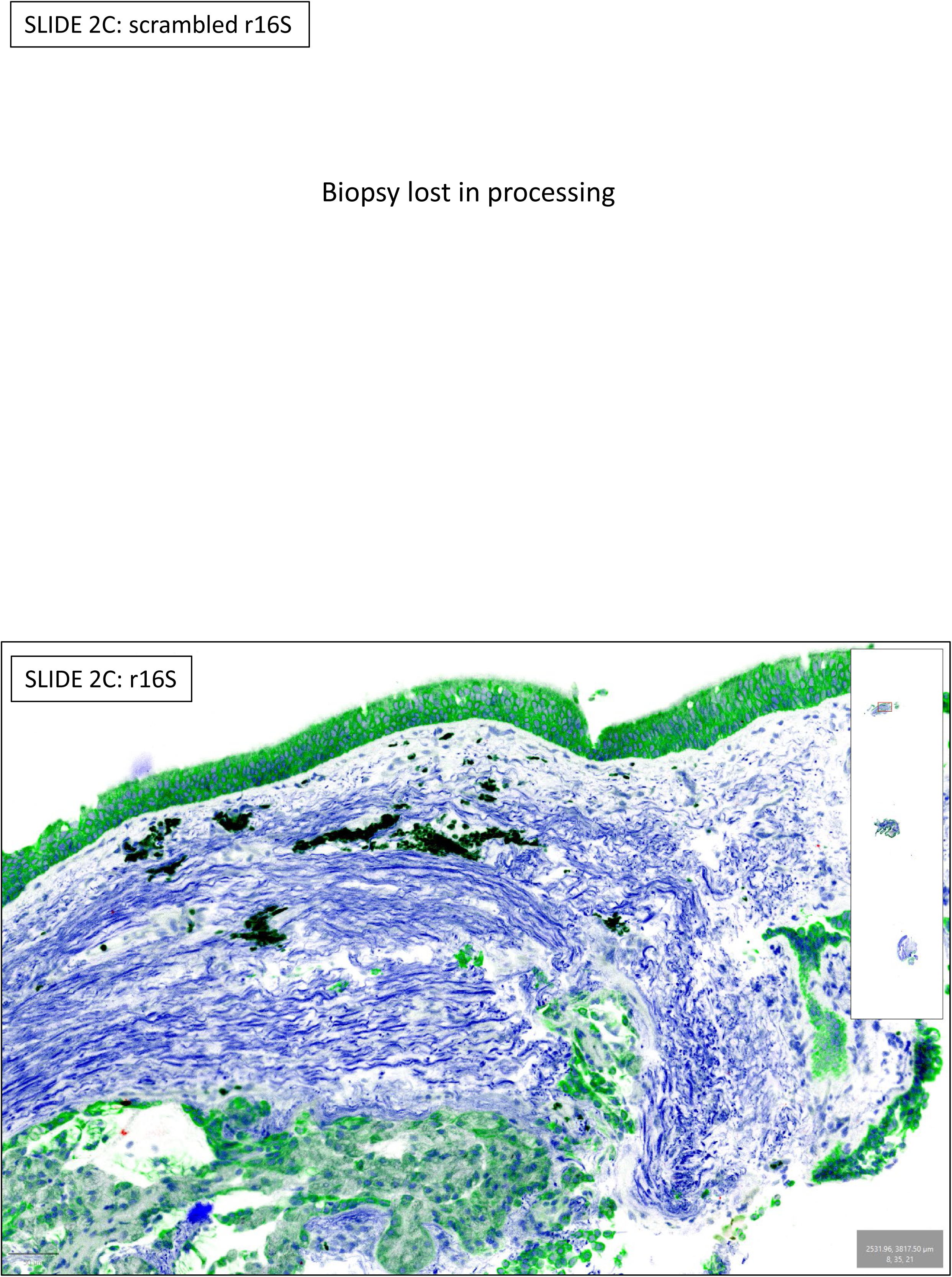
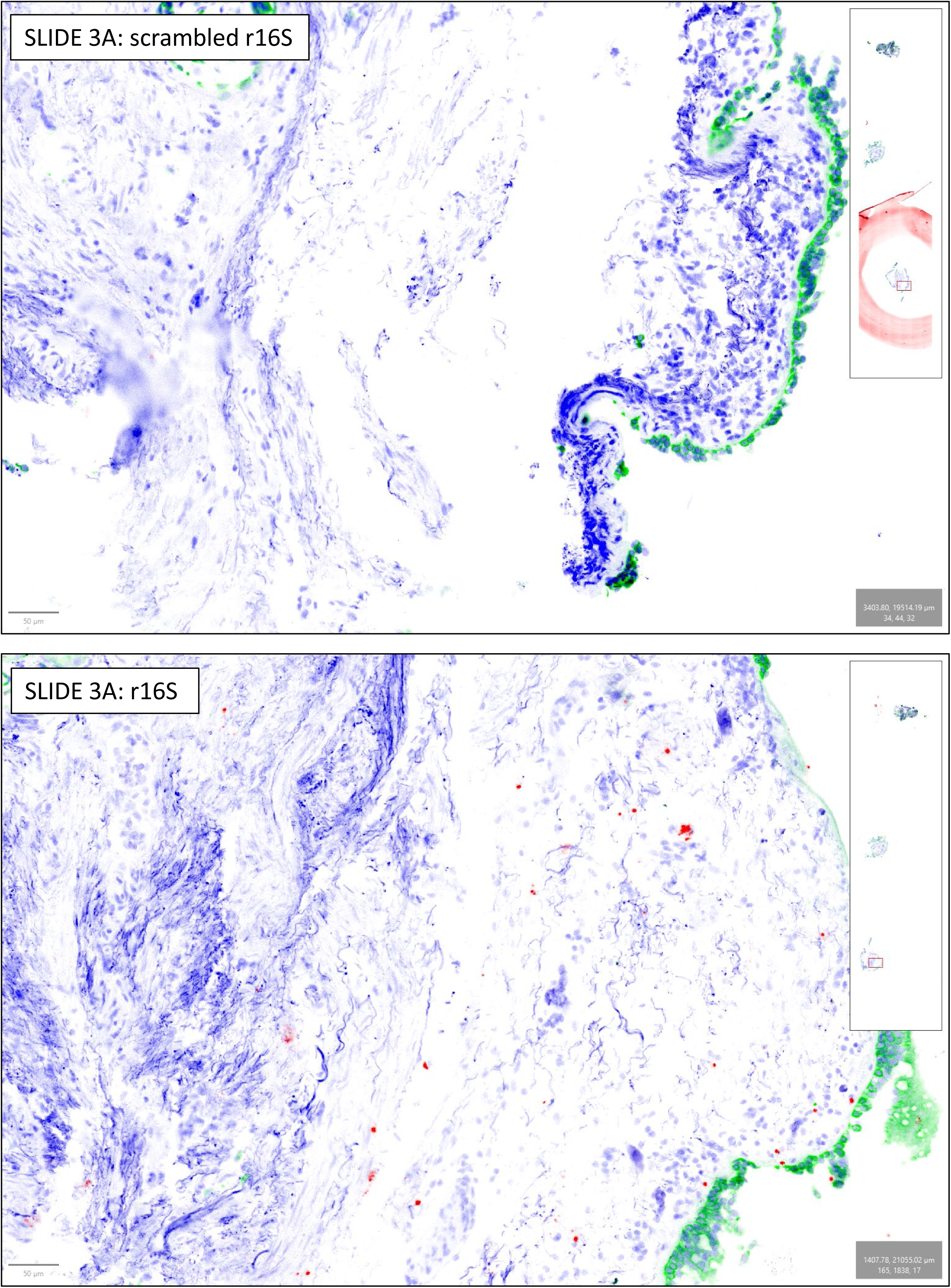
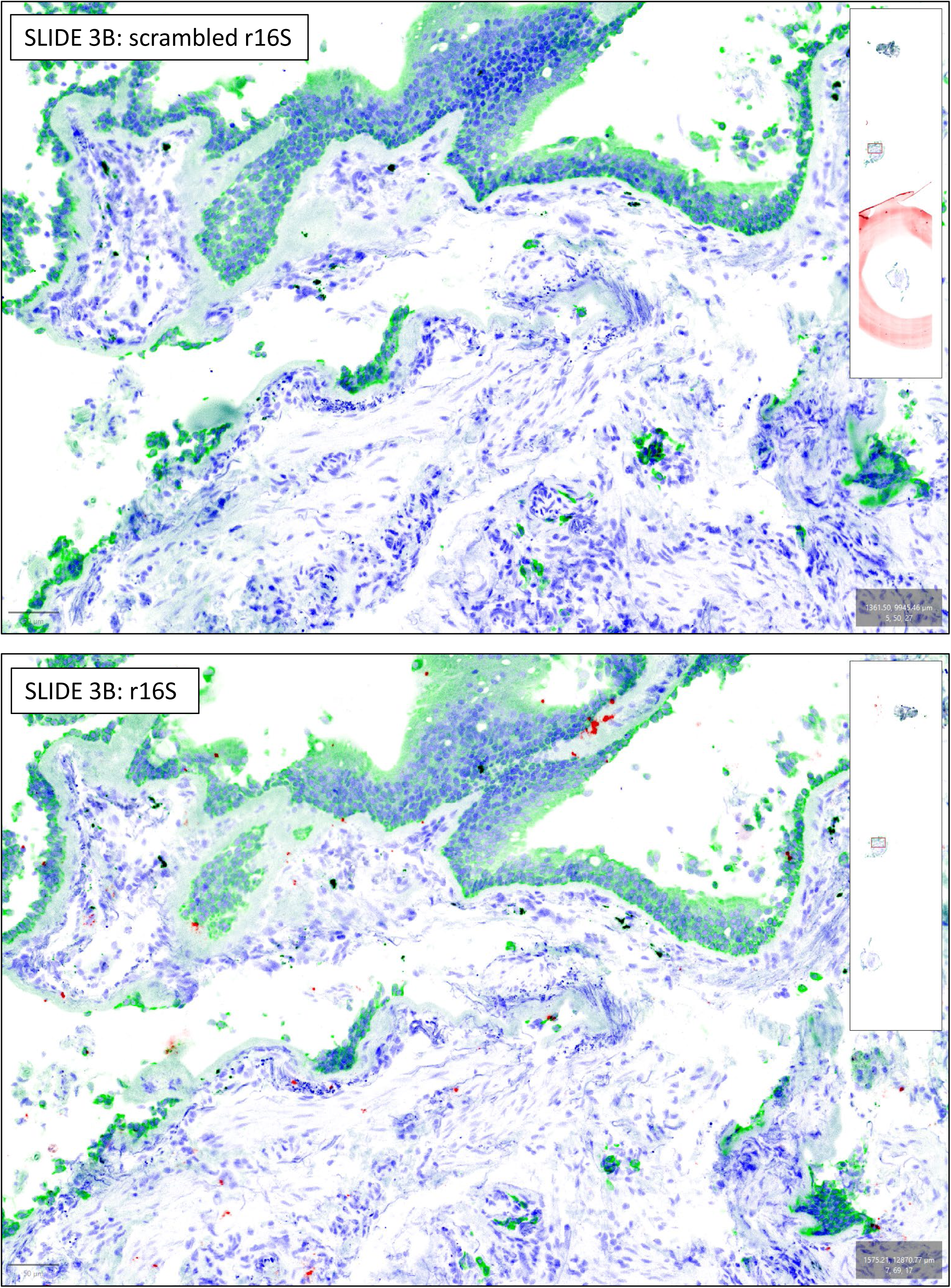
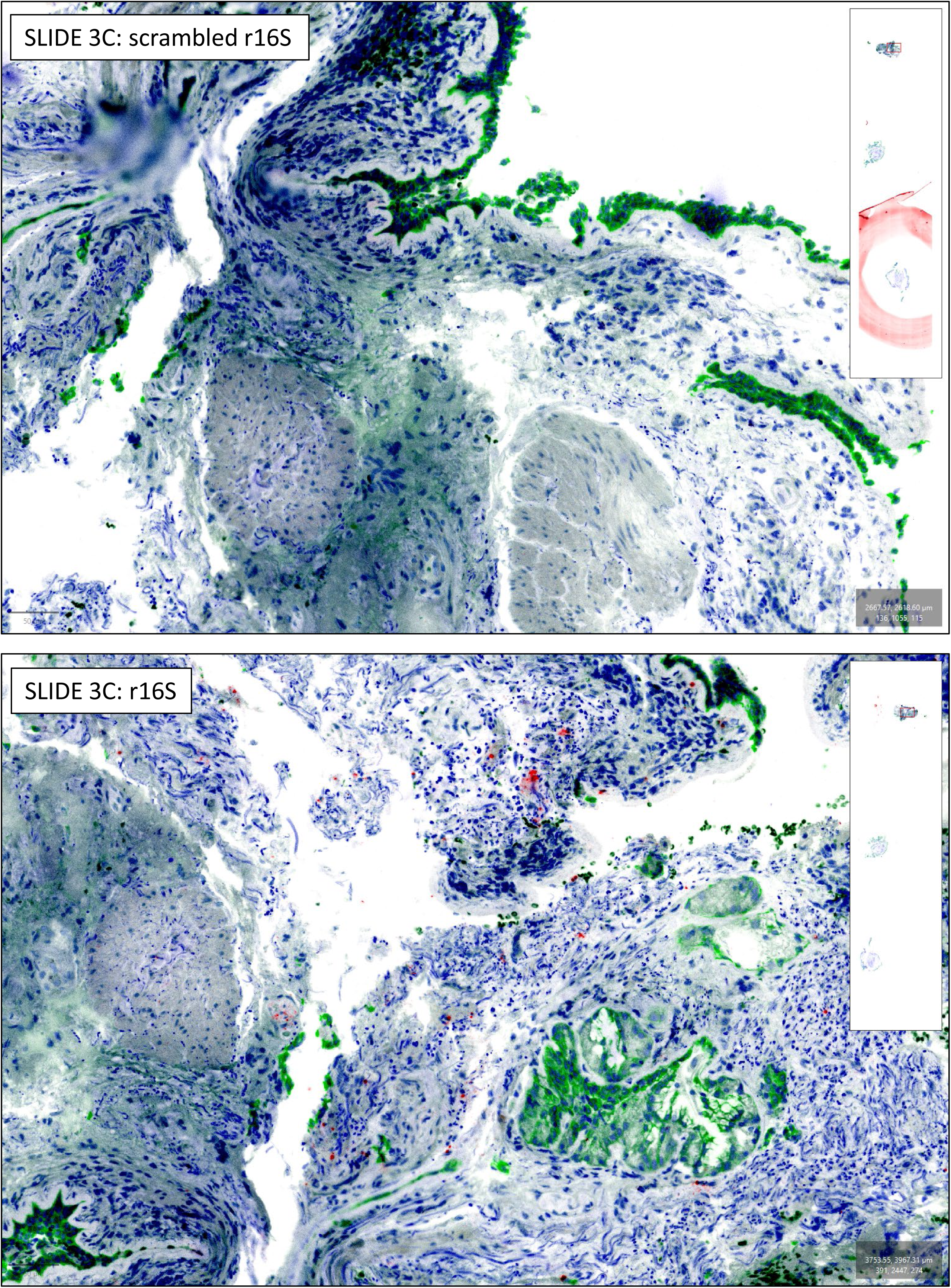

Network module eigengene phylogeny and correlations.

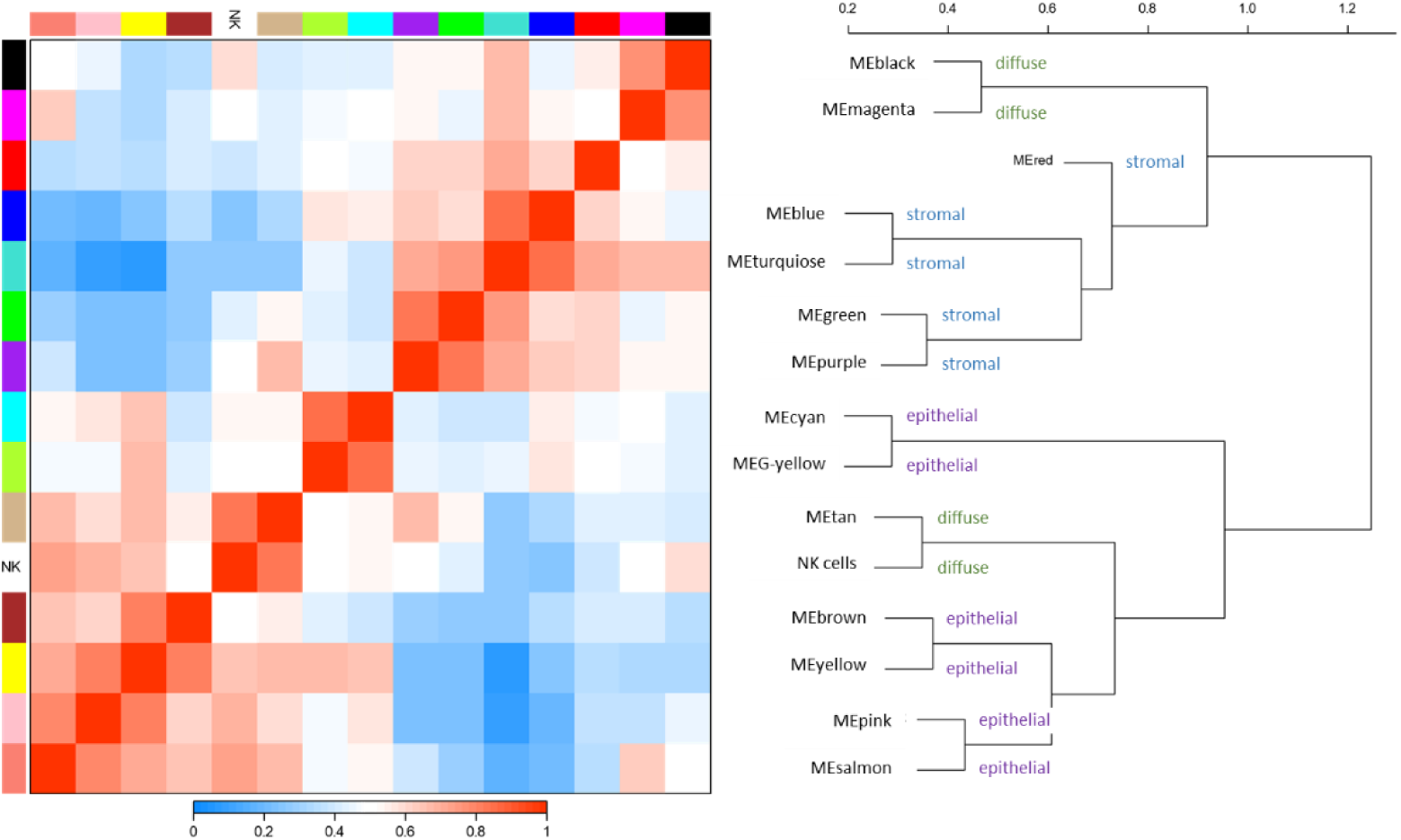

Strong correlations are seen between the eigenvectors of adjacent networks, indicating overlapping gene content. Stromal and epithelial networks are in different branches of the phylogenetic tree.

## References

1. S. T. Holgate, The sentinel role of the airway epithelium in asthma pathogenesis. Immunol Rev 242, 205–219 (2011).

2. ISAAC, Worldwide variation in prevalence of symptoms of asthma, allergic rhinoconjunctivitis, and atopic eczema: ISAAC. The International Study of Asthma and Allergies in Childhood (ISAAC) Steering Committee. Lancet 351, 1225–1232 (1998).

3. L. Pembrey et al., Asthma inflammatory phenotypes on four continents: most asthma is non-eosinophilic. International journal of epidemiology 52, 611–623 (2023).

4. M. F. Moffatt et al., A large-scale, consortium-based genomewide association study of asthma. N Engl J Med 363, 1211–1221 (2010).

5. M. Hilty et al., Disordered microbial communities in asthmatic airways. PLoS One 5, e8578 (2010).

6. T. Ichinohe et al., Microbiota regulates immune defense against respiratory tract influenza A virus infection. Proc Natl Acad Sci U S A 108, 5354–5359 (2011).

7. R. L. Brown, R. P. Sequeira, T. B. Clarke, The microbiota protects against respiratory infection via GM-CSF signaling. Nat Commun 8, 1512 (2017).

8. D. Yang et al., Many chemokines including CCL20/MIP-3alpha display antimicrobial activity. J Leukoc Biol 74, 448–455 (2003).

9. Y. Lu et al., Early-Life Antibiotic Exposure and Childhood Asthma Trajectories: A National Population-Based Birth Cohort. Antibiotics 12, 314 (2023).

10. M. J. Ege et al., Exposure to environmental microorganisms and childhood asthma. N Engl J Med 364, 701–709 (2011).

11. B. Sozanska, M. Blaszczyk, N. Pearce, P. Cullinan, Atopy and allergic respiratory disease in rural Poland before and after accession to the European Union. J Allergy Clin Immunol 133, 1347–1353 (2014).

12. R. J. Farrell, J. T. LaMont, Microbial factors in inflammatory bowel disease. Gastroenterology Clinics of North America 31, 41–62 (2002).

13. Y. J. Huang et al., Airway microbiota and bronchial hyperresponsiveness in patients with suboptimally controlled asthma. J Allergy Clin Immunol 127, 372–381 e373 (2011).

14. W. H. Man, W. A. de Steenhuijsen Piters, D. Bogaert, The microbiota of the respiratory tract: gatekeeper to respiratory health. Nat Rev Microbiol 15, 259–270 (2017).

15. M. A. Brown, M. Jabeen, G. Bharj, T. S. C. Hinks, Non-typeable Haemophilus influenzae airways infection: the next treatable trait in asthma? Eur Respir Rev 31, (2022).

16. J. L. Simpson et al., Airway dysbiosis: Haemophilus influenzae and Tropheryma in poorly controlled asthma. Eur Respir J 47, 792–800 (2016).

17. M. F. Jabeen et al., Identifying Bacterial Airways Infection in Stable Severe Asthma Using Oxford Nanopore Sequencing Technologies. Microbiol Spectr 10, e0227921 (2022).

18. Y.-d. Gao et al., Effect of Haemophilus influenzae, Streptococcus pneumoniae and influenza vaccinations on infections, immune response and asthma control in preschool children with asthma. Allergy n/a.

19. S. Johnston et al., Community study of role of viral infections in exacerbations of asthma in 9-11 year old children. BMJ 310, p1225–1229 (1995).

20. W. Cookson, M. Moffatt, G. Rapeport, J. Quint, A Pandemic Lesson for Global Lung Diseases: Exacerbations Are Preventable. Am J Respir Crit Care Med 205, 1271–1280 (2022).

21. E. E. Schadt, Molecular networks as sensors and drivers of common human diseases. Nature 461, 218–223 (2009).

22. E. M. Turek et al., Airway microbial communities, smoking and asthma in a general population sample. EBioMedicine 71, 103538 (2021).

23. P. Danaher et al., Advances in mixed cell deconvolution enable quantification of cell types in spatial transcriptomic data. Nature Communications 13, 385 (2022).

24. P. Langfelder, S. Horvath, WGCNA: an R package for weighted correlation network analysis. BMC Bioinformatics 9, 559 (2008).

25. B. Lin et al., Airway hillocks are injury-resistant reservoirs of unique plastic stem cells. Nature 629, 869–877 (2024).

26. British_Thoracic_Society, BTS/SIGN British Guideline for the management of asthma (SIGN 153) Available at: https://www.sign.ac.uk/sign-153-british-guideline-on-the-management-of-asthma.html. (2016).

27. A. Bourdin et al., Phenotyping of Severe Asthma in the Era of Broad-Acting Anti-Asthma Biologics. The Journal of Allergy and Clinical Immunology: In Practice 12, 809–823 (2024).

28. F. Demenais et al., Multiancestry association study identifies new asthma risk loci that colocalize with immune-cell enhancer marks. Nat Genet 50, 42–53 (2018).

29. M. F. Moffatt et al., Genetic variants regulating ORMDL3 expression contribute to the risk of childhood asthma. Nature 448, 470–473 (2007).

30. M. Caliskan et al., Rhinovirus wheezing illness and genetic risk of childhood-onset asthma. N Engl J Med 368, 1398–1407 (2013).

31. T. Nakatsuji et al., The microbiome extends to subepidermal compartments of normal skin. Nat Commun 4, 1431 (2013).

32. L. Bay et al., Universal Dermal Microbiome in Human Skin. mBio 11, 10.1128/mbio.02945-02919 (2020).

33. B. Lötstedt, M. Stražar, R. Xavier, A. Regev, S. Vickovic, Spatial host-microbiome sequencing reveals niches in the mouse gut. Nat Biotechnol, (2023).

34. Y. Ma et al., Intratumor microbiome-derived butyrate promotes lung cancer metastasis. Cell Reports Medicine 5, 101488 (2024).

35. L. Cuthbertson et al., Genomic attributes of airway commensal bacteria and mucosa. Commun Biol 7, 171 (2024).

36. J. A. Coultas, J. Cafferkey, P. Mallia, S. L. Johnston, Experimental Antiviral Therapeutic Studies for Human Rhinovirus Infections. J Exp Pharmacol 13, 645–659 (2021).

37. S.-J. Lee, I. Depoortere, H. Hatt, Therapeutic potential of ectopic olfactory and taste receptors. Nature Reviews Drug Discovery 18, 116–138 (2019).

38. H. Chen et al., Activation of STAT6 by STING Is Critical for Antiviral Innate Immunity. Cell 147, 436–446 (2011).

39. V. Mehraj, R. Ponte, J.-P. Routy, The Dynamic Role of the IL-33/ST2 Axis in Chronic Viral-infections: Alarming and Adjuvanting the Immune Response. EBioMedicine 9, 37–44 (2016).

40. S. P. Sanders, E. S. Siekierski, J. D. Porter, S. M. Richards, D. Proud, Nitric oxide inhibits rhinovirus-induced cytokine production and viral replication in a human respiratory epithelial cell line. J Virol 72, 934–942 (1998).

41. L. Cuthbertson et al., The impact of persistent bacterial bronchitis on the pulmonary microbiome of children. PLoS One 12, e0190075 (2017).

42. S. J. Salter et al., Reagent and laboratory contamination can critically impact sequence-based microbiome analyses. BMC biology 12, 87 (2014).

43. P. Bankhead et al., QuPath: Open source software for digital pathology image analysis. Sci Rep 7, 16878 (2017).

44. N. H. de Klerk, A. W. Musk, A. James, J. J. Glancy, W. O. Cookson, Comparison of chest radiograph reading methods for assessing progress of pneumoconiosis over 10 years in Wittenoom crocidolite workers. Br J Ind Med 47, 127–131 (1990).

