## Supplementary material for "Microbiome diversity, intra-mucosal bacteria and immune integration within normal and asthmatic airway mucosa": Subject details

### Frequencies

#### Asthma

| | | Frequency | Percent | Valid Percent | Cumulative Percent |
|---|---|---|---|---|---|
| Valid | 0 | 44 | 39.6 | 39.6 | 39.6 |
| | 1 | 67 | 60.4 | 60.4 | 100.0 |
| | Total | 111 | 100.0 | 100.0 | |

#### Severity

| | | Frequency | Percent | Valid Percent | Cumulative Percent |
|---|---|---|---|---|---|
| Valid | 0 | 44 | 39.6 | 40.0 | 40.0 |
| | 1 | 30 | 27.0 | 27.3 | 67.3 |
| | 2 | 36 | 32.4 | 32.7 | 100.0 |
| | Total | 110 | 99.1 | 100.0 | |
| Missing | System | 1 | 0.9 | | |
| Total | | 111 | 100.0 | | |

#### Sex

| | | Frequency | Percent | Valid Percent | Cumulative Percent |
|---|---|---|---|---|---|
| Valid | 1 | 57 | 51.4 | 51.4 | 51.4 |
| | 2 | 54 | 48.6 | 48.6 | 100.0 |
| | Total | 111 | 100.0 | 100.0 | |

#### Ethnicity

| | | Frequency | Percent | Valid Percent | Cumulative Percent |
|---|---|---|---|---|---|
| Valid | 1 | 98 | 88.3 | 88.3 | 88.3 |
| | 2 | 8 | 7.2 | 7.2 | 95.5 |
| | 3 | 5 | 4.5 | 4.5 | 100.0 |
| | Total | 111 | 100.0 | 100.0 | |

#### Smoking

| | | Frequency | Percent | Valid Percent | Cumulative Percent |
|---|---|---|---|---|---|
| Valid | 0 | 95 | 85.6 | 85.6 | 85.6 |
| | 1 | 16 | 14.4 | 14.4 | 100.0 |
| | Total | 111 | 100.0 | 100.0 | |

#### Denudation

| | | Frequency | Percent | Valid Percent | Cumulative Percent |
|---|---|---|---|---|---|
| Valid | 0 | 54 | 48.6 | 55.7 | 55.7 |

| | | | | | |
|---|---|---|---|---|---|
| | 1 | 43 | 38.7 | 44.3 | 100.0 |
| | Total | 97 | 87.4 | 100.0 | |
| Missing | System | 14 | 12.6 | | |
| Total | | 111 | 100.0 | | |

**Epithelialhyperplasia**

| | | Frequency | Percent | Valid Percent | Cumulative Percent |
|---|---|---|---|---|---|
| Valid | 0 | 71 | 64.0 | 73.2 | 73.2 |
| | 1 | 26 | 23.4 | 26.8 | 100.0 |
| | Total | 97 | 87.4 | 100.0 | |
| Missing | System | 14 | 12.6 | | |
| Total | | 111 | 100.0 | | |

**Gobletcellhyperplasia**

| | | Frequency | Percent | Valid Percent | Cumulative Percent |
|---|---|---|---|---|---|
| Valid | 0 | 80 | 72.1 | 82.5 | 82.5 |
| | 1 | 17 | 15.3 | 17.5 | 100.0 |
| | Total | 97 | 87.4 | 100.0 | |
| Missing | System | 14 | 12.6 | | |
| Total | | 111 | 100.0 | | |

**Basementmembrane**

| | | Frequency | Percent | Valid Percent | Cumulative Percent |
|---|---|---|---|---|---|
| Valid | 0 | 42 | 37.8 | 43.3 | 43.3 |
| | 1 | 55 | 49.5 | 56.7 | 100.0 |
| | Total | 97 | 87.4 | 100.0 | |
| Missing | System | 14 | 12.6 | | |
| Total | | 111 | 100.0 | | |

**Inflammation**

| | | Frequency | Percent | Valid Percent | Cumulative Percent |
|---|---|---|---|---|---|
| Valid | 0 | 43 | 38.7 | 45.3 | 45.3 |
| | 1 | 43 | 38.7 | 45.3 | 90.5 |
| | 2 | 9 | 8.1 | 9.5 | 100.0 |
| | Total | 95 | 85.6 | 100.0 | |
| Missing | System | 16 | 14.4 | | |
| Total | | 111 | 100.0 | | |

**Macrophages**

| | | Frequency | Percent | Valid Percent | Cumulative Percent |
|---|---|---|---|---|---|
| Valid | 0 | 93 | 83.8 | 97.9 | 97.9 |
| | 1 | 2 | 1.8 | 2.1 | 100.0 |
| | Total | 95 | 85.6 | 100.0 | |
| Missing | System | 16 | 14.4 | | |

| | | | |
|---|---|---|---|
| Total | | 111 | 100.0 |

#### Mastcells

| | | Frequency | Percent | Valid Percent | Cumulative Percent |
|---|---|---|---|---|---|
| Valid | 0 | 86 | 77.5 | 90.5 | 90.5 |
| | 1 | 9 | 8.1 | 9.5 | 100.0 |
| | Total | 95 | 85.6 | 100.0 | |
| Missing | System | 16 | 14.4 | | |
| Total | | 111 | 100.0 | | |

#### Eosinophils

| | | Frequency | Percent | Valid Percent | Cumulative Percent |
|---|---|---|---|---|---|
| Valid | 0 | 72 | 64.9 | 75.8 | 75.8 |
| | 1 | 22 | 19.8 | 23.2 | 98.9 |
| | 2 | 1 | 0.9 | 1.1 | 100.0 |
| | Total | 95 | 85.6 | 100.0 | |
| Missing | System | 16 | 14.4 | | |
| Total | | 111 | 100.0 | | |

#### Bacteria

| | | Frequency | Percent | Valid Percent | Cumulative Percent |
|---|---|---|---|---|---|
| Valid | 0 | 94 | 84.7 | 98.9 | 98.9 |
| | 1 | 1 | 0.9 | 1.1 | 100.0 |
| | Total | 95 | 85.6 | 100.0 | |
| Missing | System | 16 | 14.4 | | |
| Total | | 111 | 100.0 | | |

#### Descriptive Statistics

| Subjects | N | Minimum | Maximum | Mean | Std. Deviation |
|---|---|---|---|---|---|
| Age | 111 | 18 | 72 | 41.69 | 14.315 |
| Abs_Neu | 109 | 1.7 | 12.9 | 4.104 | 1.8234 |
| Abs_Eos | 109 | 0.00 | 0.94 | 0.2572 | 0.20014 |
| BMI | 111 | 19.07 | 35.35 | 25.8131 | 3.54285 |

| Biopsy bacterial scores | N | Minimum | Maximum | Mean | Std. Deviation |
|---|---|---|---|---|---|
| Epithelium_16S | 37 | 0 | 2 | 0.81 | 0.593 |
| Base Memb_16S | 37 | 0.0 | 3.0 | 1.905 | 0.9040 |
| Stroma_16S | 37 | 0.50 | 3.00 | 2.3784 | 0.76744 |
