## Supplementary material for "Microbiome diversity, intra-mucosal bacteria and immune integration within normal and asthmatic airway mucosa": Correlation analyses

### Table S2. Correlations between key variables

**Correlations**

| Phenotype¶ | | Asthma | Severity | Blood eosinophils | Shannon diversity* | Actinomyces 6974 | Selenomonas 3_19868 | *H. influenzae* | *T. whipplei* | Denudation | Goblet cell hyperplasia | Abnormal B. membrane | Visible eosinophils | Visible bacteria | Epithelium 16S score | BM 16S score | Stroma 16S score |
|---|---|---|---|---|---|---|---|---|---|---|---|---|---|---|---|---|---|
| Asthma | Pearson r | | 0.891 | 0.351 | -0.203 | -0.197 | -0.161 | 0.093 | 0.169 | 0.121 | 0.259 | 0.364 | 0.205 | 0.082 | 0.224 | -0.117 | -0.037 |
| | Sig. (2-tailed) | | 0.000 | 0.000 | 0.009 | 0.011 | 0.038 | 0.231 | 0.029 | 0.238 | 0.010 | 0.000 | 0.046 | 0.427 | 0.159 | 0.467 | 0.819 |
| | N | 111 | 110 | 109 | 167 | 167 | 167 | 167 | 167 | 97 | 97 | 97 | 95 | 95 | 41 | 41 | 41 |
| Severity | Pearson r | 0.891 | | 0.296 | -0.150 | -0.174 | -0.071 | 0.058 | 0.127 | 0.101 | 0.331 | 0.322 | 0.159 | 0.009 | 0.212 | -0.215 | -0.152 |
| | Sig. (2-tailed) | 0.000 | | 0.002 | 0.053 | 0.025 | 0.360 | 0.455 | 0.102 | 0.328 | 0.001 | 0.001 | 0.125 | 0.930 | 0.188 | 0.182 | 0.350 |
| | N | 110 | 110 | 108 | 166 | 166 | 166 | 166 | 166 | 96 | 96 | 96 | 94 | 94 | 40 | 40 | 40 |
| Blood eosinophils | Pearson r | 0.351 | 0.296 | | -0.085 | -0.131 | -0.212 | -0.027 | 0.139 | -0.018 | 0.085 | 0.177 | 0.271 | 0.001 | 0.340 | 0.038 | 0.112 |
| | Sig. (2-tailed) | 0.000 | 0.000 | | 0.281 | 0.097 | 0.007 | 0.737 | 0.076 | 0.862 | 0.414 | 0.087 | 0.009 | 0.995 | 0.032 | 0.816 | 0.492 |
| | N | 163 | 162 | 163 | 163 | 163 | 163 | 163 | 163 | 95 | 95 | 95 | 93 | 93 | 40 | 40 | 40 |
| Shannon diversity | Pearson r | -0.203 | -0.150 | -0.085 | | 0.320 | 0.270 | -0.093 | -0.454 | 0.134 | -0.113 | 0.045 | 0.107 | -0.354 | -0.044 | 0.269 | 0.491 |
| | Sig. (2-tailed) | 0.009 | 0.053 | 0.281 | | 0.000 | 0.000 | 0.229 | 0.000 | 0.108 | 0.176 | 0.593 | 0.207 | 0.000 | 0.722 | 0.028 | 0.000 |
| | N | 167 | 166 | 163 | 167 | 167 | 167 | 167 | 167 | 145 | 145 | 145 | 142 | 142 | 67 | 67 | 67 |
| Actinomyces 6974 | Pearson r | -0.197 | -0.174 | -0.131 | 0.320 | | 0.678 | 0.081 | -0.133 | -0.205 | -0.061 | -0.066 | -0.128 | -0.079 | 0.133 | 0.052 | 0.096 |
| | Sig. (2-tailed) | 0.011 | 0.025 | 0.097 | 0.000 | | 0.000 | 0.296 | 0.086 | 0.013 | 0.470 | 0.429 | 0.128 | 0.349 | 0.284 | 0.674 | 0.442 |
| | N | 167 | 166 | 163 | 167 | 167 | 167 | 167 | 167 | 145 | 145 | 145 | 142 | 142 | 67 | 67 | 67 |
| Selenomonas 3_19868 | Pearson r | -0.161 | -0.071 | -0.212 | 0.270 | 0.678 | | 0.058 | -0.074 | -0.223 | 0.028 | -0.075 | -0.215 | -0.074 | -0.298 | -0.089 | 0.034 |
| | Sig. (2-tailed) | 0.038 | 0.360 | 0.007 | 0.000 | 0.000 | | 0.460 | 0.341 | 0.007 | 0.740 | 0.372 | 0.010 | 0.382 | 0.014 | 0.473 | 0.783 |
| | N | 167 | 166 | 163 | 167 | 167 | 167 | 167 | 167 | 145 | 145 | 145 | 142 | 142 | 67 | 67 | 67 |
| *H. influenzae* | Pearson r | 0.093 | 0.058 | -0.027 | -0.093 | 0.081 | 0.058 | | 0.111 | -0.063 | 0.124 | -0.065 | -0.062 | 0.032 | 0.142 | -0.117 | -0.327 |
| | Sig. (2-tailed) | 0.231 | 0.455 | 0.737 | 0.229 | 0.296 | 0.460 | | 0.155 | 0.448 | 0.139 | 0.437 | 0.467 | 0.707 | 0.253 | 0.346 | 0.007 |
| | N | 167 | 166 | 163 | 167 | 167 | 167 | 167 | 167 | 145 | 145 | 145 | 142 | 142 | 67 | 67 | 67 |
| *T. whipplei* | Pearson r | 0.169 | 0.127 | 0.139 | -0.454 | -0.133 | -0.074 | 0.111 | | -0.187 | 0.089 | -0.015 | 0.003 | 0.316 | 0.200 | -0.345 | -0.355 |
| | Sig. (2-tailed) | 0.029 | 0.102 | 0.076 | 0.000 | 0.086 | 0.341 | 0.155 | | 0.024 | 0.288 | 0.857 | 0.968 | 0.000 | 0.104 | 0.004 | 0.003 |
| | N | 167 | 166 | 163 | 167 | 167 | 167 | 167 | 167 | 145 | 145 | 145 | 142 | 142 | 67 | 67 | 67 |
| Denudation | Pearson r | 0.121 | 0.101 | -0.018 | 0.134 | -0.205 | -0.223 | -0.063 | -0.187 | | -0.123 | 0.411 | 0.260 | -0.123 | 0.035 | 0.411 | 0.123 |
| | Sig. (2-tailed) | 0.238 | 0.328 | 0.862 | 0.108 | 0.013 | 0.007 | 0.448 | 0.024 | | 0.141 | 0.000 | 0.002 | 0.145 | 0.828 | 0.008 | 0.442 |
| | N | 97 | 96 | 95 | 145 | 145 | 145 | 145 | 145 | 145 | 145 | 145 | 142 | 142 | 41 | 41 | 41 |
| Goblet cell hyperplasia | Pearson r | 0.259 | 0.331 | 0.085 | -0.113 | -0.061 | 0.028 | 0.124 | 0.089 | -0.123 | | 0.017 | -0.032 | 0.295 | -0.061 | -0.289 | -0.170 |
| | Sig. (2-tailed) | 0.010 | 0.001 | 0.414 | 0.176 | 0.470 | 0.740 | 0.139 | 0.288 | 0.141 | | 0.841 | 0.701 | 0.000 | 0.706 | 0.067 | 0.287 |
| | N | 97 | 96 | 95 | 145 | 145 | 145 | 145 | 145 | 145 | 145 | 145 | 142 | 142 | 41 | 41 | 41 |
| Abnormal basement membrane | Pearson r | 0.364 | 0.322 | 0.177 | 0.045 | -0.066 | -0.075 | -0.065 | -0.015 | 0.411 | 0.017 | | 0.289 | -0.144 | 0.132 | 0.441 | 0.170 |
| | Sig. (2-tailed) | 0.000 | 0.001 | 0.087 | 0.593 | 0.429 | 0.372 | 0.437 | 0.857 | 0.000 | 0.841 | | 0.000 | 0.088 | 0.411 | 0.004 | 0.288 |
| | N | 97 | 96 | 95 | 145 | 145 | 145 | 145 | 145 | 145 | 145 | 145 | 142 | 142 | 41 | 41 | 41 |
| Visible eosinophils | Pearson r | 0.205 | 0.159 | 0.271 | 0.107 | -0.128 | -0.215 | -0.062 | 0.003 | 0.260 | -0.032 | 0.289 | | -0.071 | 0.050 | 0.270 | 0.239 |
| | Sig. (2-tailed) | 0.046 | 0.125 | 0.009 | 0.207 | 0.128 | 0.010 | 0.467 | 0.968 | 0.002 | 0.701 | 0.000 | | 0.398 | 0.755 | 0.087 | 0.133 |
| | N | 95 | 94 | 93 | 142 | 142 | 142 | 142 | 142 | 142 | 142 | 142 | 142 | 142 | 41 | 41 | 41 |
| Visible bacteria | Pearson r | 0.082 | 0.009 | 0.001 | -0.354 | -0.079 | -0.074 | 0.032 | 0.316 | -0.123 | 0.295 | -0.144 | -0.071 | | -0.075 | -0.234 | -0.302 |
| | Sig. (2-tailed) | 0.427 | 0.930 | 0.995 | 0.000 | 0.349 | 0.382 | 0.707 | 0.000 | 0.145 | 0.000 | 0.088 | 0.398 | | 0.640 | 0.140 | 0.055 |
| | N | 95 | 94 | 93 | 142 | 142 | 142 | 142 | 142 | 142 | 142 | 142 | 142 | 142 | 41 | 41 | 41 |
| Epithelium 16S score | Pearson r | 0.224 | 0.212 | 0.340 | -0.044 | 0.133 | -0.298 | 0.142 | 0.200 | 0.035 | -0.061 | 0.132 | 0.050 | -0.075 | | 0.344 | 0.090 |
| | Sig. (2-tailed) | 0.159 | 0.188 | 0.032 | 0.722 | 0.284 | 0.014 | 0.253 | 0.104 | 0.828 | 0.706 | 0.411 | 0.755 | 0.640 | | 0.028 | 0.575 |
| | N | 41 | 40 | 40 | 67 | 67 | 67 | 67 | 67 | 41 | 41 | 41 | 41 | 41 | 41 | 41 | 41 |
| Basement membrane 16S score | Pearson r | -0.117 | -0.215 | 0.038 | 0.269 | 0.052 | -0.089 | -0.117 | -0.345 | 0.411 | -0.289 | 0.441 | 0.270 | -0.234 | 0.344 | | 0.580 |
| | Sig. (2-tailed) | 0.467 | 0.182 | 0.816 | 0.028 | 0.674 | 0.473 | 0.346 | 0.004 | 0.008 | 0.067 | 0.004 | 0.087 | 0.140 | 0.028 | | 0.000 |
| | N | 41 | 40 | 40 | 67 | 67 | 67 | 67 | 67 | 41 | 41 | 41 | 41 | 41 | 41 | 41 | 41 |
| Stroma 16S score | Pearson r | -0.037 | -0.152 | 0.112 | 0.491 | 0.096 | 0.034 | -0.327 | -0.355 | 0.123 | -0.170 | 0.170 | 0.239 | -0.302 | 0.090 | 0.580 | |
| | Sig. (2-tailed) | 0.819 | 0.350 | 0.492 | 0.000 | 0.442 | 0.783 | 0.007 | 0.003 | 0.442 | 0.287 | 0.288 | 0.133 | 0.055 | 0.575 | 0.000 | |
| | N | 41 | 40 | 40 | 67 | 67 | 67 | 67 | 67 | 41 | 41 | 41 | 41 | 41 | 41 | 41 | 41 |

IBM SPPS Statistics 29.0.2.0 (171) datafile "Moffatt et al 16S mapfile with histology 16Score 2024_08_12.csv"

**¶Phenotypes**

- Clinical
- 16S rRNA community analyses
- H&E histology
- 16S probe hits microscopy

*Microbial abundances from all lower lobe and some upper lobe samples, so total n=167
