## Supplementary material for "Microbiome diversity, intra-mucosal bacteria and immune integration within normal and asthmatic airway mucosa": Stepwise predictions oaf asthma status

Stepwise regression of asthma predictors

**Variables in the Equation**

| | | B | S.E. | Wald | df | Sig. | Exp(B) | 95% C.I.for EXP(B) Lower | Upper |
|---|---|---|---|---|---|---|---|---|---|
| Step 1[a] | Basement membrane | 1.518 | 0.380 | 15.948 | 1 | 0.000 | 4.561 | 2.166 | 9.606 |
| | Constant | -0.208 | 0.264 | 0.618 | 1 | 0.432 | 0.813 | | |
| Step 2[b] | Shannon_diversity | -2.675 | 0.646 | 17.150 | 1 | 0.000 | 0.069 | 0.019 | 0.244 |
| | Basementmembrane | 2.081 | 0.464 | 20.136 | 1 | 0.000 | 8.012 | 3.228 | 19.881 |
| | Constant | 8.043 | 2.017 | 15.895 | 1 | 0.000 | 3110.816 | | |
| Step 3[c] | Abs_Eos | 3.783 | 1.465 | 6.669 | 1 | 0.010 | 43.967 | 2.489 | 776.571 |
| | Shannon_diversity | -2.637 | 0.686 | 14.784 | 1 | 0.000 | 0.072 | 0.019 | 0.274 |
| | Basement membrane | 1.875 | 0.482 | 15.128 | 1 | 0.000 | 6.524 | 2.535 | 16.785 |
| | Constant | 7.180 | 2.144 | 11.209 | 1 | 0.001 | 1312.650 | | |

a. Variable(s) entered on step 1: Basementmembrane.

b. Variable(s) entered on step 2: Shannon_diversity.

c. Variable(s) entered on step 3: Abs_Eos.

**Model Summary**

| Step | -2 Log likelihood | Cox & Snell R Square | Nagelkerke R Square |
|---|---|---|---|
| 1 | 162.543[a] | 0.116 | 0.159 |
| 2 | 136.086[b] | 0.270 | 0.371 |
| 3 | 127.633[b] | 0.314 | 0.431 |

a. Estimation terminated at iteration number 4 because parameter estimates changed by less than .001.

b. Estimation terminated at iteration number 6 because parameter estimates changed by less than .001.
