## Supplementary material for "Microbiome diversity, intra-mucosal bacteria and immune integration within normal and asthmatic airway mucosa": Visual scores for 16S foci

| Slide | Patient ID | Epithel_R1 | Base_Mem_R1 | Stroma_R1 | Epithel_R2 |
|---|---|---|---|---|---|
| 1a | 005 | 1 | 2 | 3 | 0 |
| 1b | 006 | 2 | 2 | 3 | 1 |
| 1c | 019 | 1 | 3 | 3 | 1 |
| 1d | 022 | 1 | 3 | 3 | 1 |
| 2a | 053 | 0 | 3 | 3 | 0 |
| 2b | 054 | 0 | 2 | 3 | 0 |
| 2c | 301 | 0 | 1 | 1 | 0 |
| 2d | 309 | | | | |
| 3a | 316 | 0 | 1 | 3 | 0 |
| 3b | 321 | 2 | 1 | 3 | 2 |
| 3c | 322 | 0 | 1 | 2 | 0 |
| 3d | 323 | 0 | 1 | 2 | 1 |
| 4a | 334 | 0 | 1 | 2 | 1 |
| 4b | 337 | 0 | 1 | 2 | 0 |
| 4c | 365 | 0 | 1 | 2 | 0 |
| 4d | 370 | 0 | 0 | 1 | 0 |
| 5a | 371 | 1 | 1 | 0 | 1 |
| 5b | 374 | 0 | 1 | 2 | 0 |
| 5c | 415 | 0 | 1 | 1 | 0 |
| 5d | 601 | 0 | 1 | 3 | 1 |
| 6a | 604 | 1 | 1 | 1 | 0 |
| 6b | 610 | 2 | 3 | 3 | 1 |
| 6c | 612 | 1 | 3 | 3 | 2 |
| 6d | 620 | 1 | 2 | 3 | 1 |
| 7a | 635 | 1 | 1 | 1 | 1 |
| 7b | 637 | 2 | 3 | 2 | 1 |
| 7c | 638 | 2 | 2 | 3 | 2 |
| 7d | 640 | 2 | 1 | 3 | 1 |
| 8a | 641 | 0 | 3 | 3 | 1 |
| 8b | 642 | 2 | 3 | 3 | 1 |
| 8c | 649 | 1 | 2 | 3 | 1 |
| 8d | 652 | 2 | 2 | 1 | 2 |
| 9a | 653 | 1 | 2 | 3 | 1 |
| 9b | 656 | 1 | 3 | 3 | 1 |
| 9c | 660 | 2 | 2 | 3 | 2 |
| 9d | 664 | 1 | 3 | 3 | 0 |
| 10a | 666 | 0 | 3 | 3 | 0 |
| 10b | 670 | 0 | 0 | 1 | 1 |
| 10c | 674 | 0 | 2 | 3 | 1 |
| 10d | 675 | 0 | 1 | 2 | 1 |
| 11a | 679 | | | | |
| 11b | 686 | 1 | 2 | 2 | 1 |
| 11c | 687 | 1 | 2 | 1 | 1 |
| 11d | 688 | 2 | 2 | 2 | 1 |

| Base_Mem_R2 | Stroma_R2 | Av_Epi | Av_BM | Av_Stroma |
|---|---|---|---|---|
| 2 | 3 | 0.5 | 2 | 3 |
| 2 | 3 | 1.5 | 2 | 3 |
| 3 | 3 | 1 | 3 | 3 |
| 3 | 3 | 1 | 3 | 3 |
| 3 | 3 | 0 | 3 | 3 |
| 2 | 3 | 0 | 2 | 3 |
| 1 | 1 | 0 | 1 | 1 |
| 1 | 3 | 0 | 1 | 3 |
| 1 | 3 | 2 | 1 | 3 |
| 1 | 3 | 0 | 1 | 2.5 |
| 1 | 2 | 0.5 | 1 | 2 |
| 1 | 2 | 0.5 | 1 | 2 |
| 1 | 2 | 0 | 1 | 2 |
| 1 | 2 | 0 | 1 | 2 |
| 0 | 2 | 0 | 0 | 1.5 |
| 0 | 1 | 1 | 0.5 | 0.5 |
| 1 | 2 | 0 | 1 | 2 |
| 1 | 1 | 0 | 1 | 1 |
| 1 | 3 | 0.5 | 1 | 3 |
| 1 | 2 | 0.5 | 1 | 1.5 |
| 3 | 3 | 1.5 | 3 | 3 |
| 3 | 3 | 1.5 | 3 | 3 |
| 2 | 3 | 1 | 2 | 3 |
| 1 | 2 | 1 | 1 | 1.5 |
| 3 | 2 | 1.5 | 3 | 2 |
| 2 | 3 | 2 | 2 | 3 |
| 1 | 3 | 1.5 | 1 | 3 |
| 3 | 3 | 0.5 | 3 | 3 |
| 3 | 3 | 1.5 | 3 | 3 |
| 2 | 3 | 1 | 2 | 3 |
| 2 | 1 | 2 | 2 | 1 |
| 2 | 3 | 1 | 2 | 3 |
| 3 | 3 | 1 | 3 | 3 |
| 3 | 2 | 2 | 2.5 | 2.5 |
| 3 | 3 | 0.5 | 3 | 3 |
| 3 | 3 | 0 | 3 | 3 |
| 1 | 1 | 0.5 | 0.5 | 1 |
| 2 | 3 | 0.5 | 2 | 3 |
| 1 | 2 | 0.5 | 1 | 2 |
| 2 | 2 | 1 | 2 | 2 |
| 2 | 2 | 1 | 2 | 1.5 |
| 2 | 2 | 1.5 | 2 | 2 |
